# SINE compounds activate exportin-1 degradation via an allosteric mechanism

**DOI:** 10.1101/2024.10.07.617049

**Authors:** Casey E. Wing, Ho Yee Joyce Fung, Bert Kwanten, Tolga Cagatay, Ashley B. Niesman, Maarten Jacquemyn, Mehdi Gharghabi, Brecht Permentier, Binita Shakya, Rhituparna Nandi, Joseph M. Ready, Trinayan Kashyap, Sharon Shacham, Yosef Landesman, Rosa Lapalombella, Dirk Daelemans, Yuh Min Chook

## Abstract

The nuclear export receptor exportin 1 (XPO1/CRM1) is often overexpressed in cancer cells, leading to the mislocalization of numerous cancer-related protein cargoes^1,2^. Selinexor, a covalent XPO1 inhibitor, and other Selective Inhibitor of Nuclear Export (SINEs) restore proper nuclear localization by blocking XPO1-cargo binding^2–7^. SINEs also induce XPO1 degradation via the Cullin-RING E3 ubiquitin ligase (CRL) substrate receptor ASB8^7^. Here we elucidate the mechanism underlying the high-affinity engagement of CRL5^ASB8^ with SINE-conjugated XPO1. Cryogenic electron microscopy (cryoEM) structures reveal that ASB8 binds to a cryptic site on XPO1, which becomes accessible only upon SINE conjugation. While molecular glue degraders typically interact with both CRL and the substrate^8–10^, SINEs bind to XPO1 without requiring interaction with ASB8 for efficient XPO1 degradation. Instead, an allosteric mechanism facilitates high affinity XPO1-ASB8 interaction, leading to XPO1 ubiquitination and degradation. ASB8-mediated degradation is also observed upon treatment of the endogenous itaconate derivate 4-octyl itaconate, which suggests a native mechanism that is inadvertently exploited by synthesized XPO1 inhibitors. This allosteric XPO1 degradation mechanism of SINE compounds expands the known modes of targeted protein degradation beyond the well-characterized molecular glue degraders and proteolysis targeting chimeras of CRL4.

## Main text

XPO1 transports hundreds to thousands of broadly functioning proteins containing a classical nuclear export signal (NES), and several types of RNAs from the nucleus to the cytoplasm of the cell^1^. XPO1 is a 123 kDa, HEAT repeat-containing, ring-shaped protein that is allosterically regulated by ligands such as the GTPase RAN and NESs of export cargoes. RAN^GTP^ binds the N-terminal HEAT repeats while NESs bind a hydrophobic groove on the convex side of the XPO1 ring that is formed by the four helices of HEAT repeats 11 and 12 (h11A, h11B, h12A and h12B) (**Figure 1a**). 89 XPO1 entries in the Protein Data Bank (PDB) reveal that XPO1 adopts two primary conformations. In its unliganded state, XPO1 has a closed NES-binding groove whereas the binding of RAN^GTP^ and NES/cargo stabilize an open groove conformation (**Figure 1a,b**). The positive cooperativity of RAN^GTP^ and NES binding to XPO1 results in assembly of functional nuclear export complexes in the nucleus where RAN^GTP^ concentration is high, and to their disassembly in the cytoplasm where RANGAP catalyzes the GTPase activity of RAN^1,11^.

**Figure 1.**
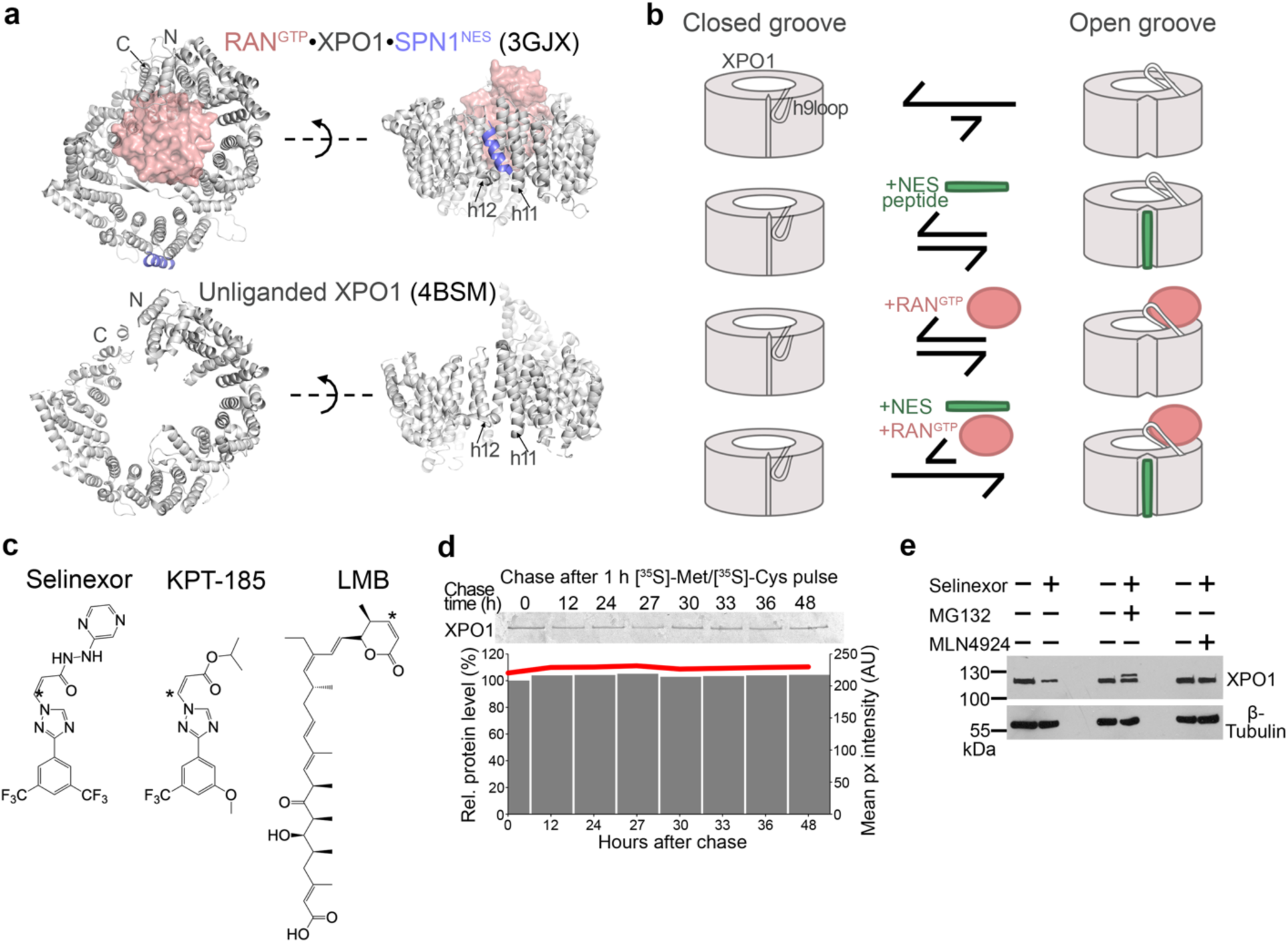
Selinexor/KPT-185 induces XPO1 degradation via a CRL mediated mechanism. **a**, Structures of a ternary RAN^GTP^•XPO1•SPN1 complex (top) and unliganded XPO1 (bottom). **b**, A schematic depicting allosteric regulation of XPO1 by its ligands, the GTPase RAN^GTP^ and the cargo’s NES, and positive cooperativity between the ligands. **c**, Chemical structures of selinexor, KPT-185 and Leptomycin B (LMB). **d**, Pulse-chase experiment in HT1080 cells, with a 1 h pulse of [^35^S]-Met/[^35^S]-Cys and XPO1 IP-ed over 48 h. **e**, XPO1 level probed after treatment of HT1080 cells with selinexor, proteasome inhibitor MG-132 and neddylation inhibitor MLN4924.

XPO1 cargoes include tumor suppressor proteins, cell cycle regulators, and apoptotic factors and oncogene mRNAs^1^. XPO1 is overexpressed in many cancers and is associated with poor prognosis due to nuclear depletion of many tumor suppressor and pro-apoptosis cargoes and increased cytoplasmic accumulation of many oncogene mRNAs. Aberrant localization of XPO1 cargoes may result in evasion of apoptosis and/or stimulate oncogenesis, conferring survival advantage to cancer cells (reviewed in^2^).

XPO1 inhibition as anti-cancer strategy included the unsuccessful Phase 1 clinical trial of the toxic lactone polyketide natural product inhibitor Leptomycin B (LMB)^2,12–14^. Smaller synthetic XPO1 inhibitors termed selective inhibitors of nuclear export (SINE) (Karyopharm Therapeutics Inc.) that are derived from the drug-like *N*-azolylacrylate XPO1 inhibitors^15,16^, effectively target a variety of cancers, and one KPT-SINE, selinexor/XPOVIO® (KPT-330), has been approved by the FDA for treatment of multiple myeloma and diffuse large B-cell lymphoma^2,3,14,17–23^. SINE compounds and LMB have α,β-unsaturated groups that undergo Michael addition reaction to covalently conjugate to Cys528 in the XPO1 NES-binding groove, directly blocking XPO1-NES/cargo interactions and hence nuclear export^15,20,24–29^.

Besides inhibiting nuclear export, many SINE compounds also induce XPO1 degradation^4–6,20^, a property not shared by LMB (**Extended Data Figure 1a**)^30^. The chemically distinct covalent XPO1 inhibitor felezonexor (CBS9106/SL-801, Stemline Therapeutics) also induces XPO1 degradation, which is suggested to be mediated by CRLs^30,31^. Our recent findings from CRISPR loss-of-function screens have revealed that the E3 ligase substrate receptor Ankyrin and SOCS box containing protein 8 (ASB8) mediates selinexor-induced XPO1 degradation^7^.

### Selinexor-induced XPO1 degradation is mediated by a Cullin Ring E3 Ligase

Several covalent XPO1 inhibitors such as selinexor, its analogs KPT-185 and KPT-8602 (eltanexor), as well as the distinct inhibitor felezonexor, cause dose– and time-dependent reduction of XPO1 levels in cells (**Extended Data Figure 1b**)^4,5,20,22,30,32,33^. All known covalent XPO1 inhibitors including selinexor, KPT-185 or LMB (**Figure 1c** shows chemical structures of some) conjugate to XPO1’s Cys528 side chain, and the mutation of this residue confers resistance to the inhibitors^15,24–29,34^. Transfection of XPO1 C528T mutation prevents SINE conjugation and SINE-induced degradation of the XPO1 mutant in cells (**Extended Data Figure 1c**), in agreement with previous findings with endogenously mutated XPO1 C528S^7^.

In the absence of inhibitor, pulse chase studies show that endogenous XPO1 in HT1080 cells is very long-lived, with XPO1 protein levels stable even after 48 hours (**Figure 1d**). Proteasome inhibitor MG132 (like bortezomib treatment as shown by others^4,35^) and neddylation inhibitor MLN4924 prevents SINE-induced XPO1 degradation, suggesting involvement of the proteasome and a CRL, respectively (**Figure 1e, Extended Data Figure 1d,e**). All the above results show that SINE-conjugated XPO1 (SINE-XPO1) is targeted by CRL for degradation by the proteasome, in agreement with our previous results^7^.

### CRL5^ASB8^ binds and ubiquitinates SINE-XPO1 for degradation

We recently reported that overexpressing the CRL substrate receptor ASB8 enhances selinexor-induced XPO1 degradation, while knocking out ASB8 inhibits this degradation^7^. ASB8 is a member of the human ANKyrin repeat and SOCS box containing protein (ASB) family. It contains an N-terminal helix (N^helix^), seven ANKyrin repeats (ANK1-7, each with antiparallel helices α1, α2) and a C-terminal SOCS box (AF-Q9H765-F1, **Figure 2a**)^36^. We hypothesize that, like other ASB proteins, the ASB8 SOCS box binds the heterodimeric Elongin B-Elongin C (ELOB/C) CRL adaptor, which in turn binds Cullin 5 (CUL5) (**Figure 2a**)^36,37^. CUL5 is anticipated to bind the RING-box protein 2 (RBX2), which recruits E2 enzymes, and neddylation is anticipated to activate the CRL5^ASB8^ complex to ubiquitinate SINE-XPO1 (**Figure 2a**)^38,39^. The ARIH2 E3 enzyme has been reported to prime ubiquitination of other CUL5 substrates^40–42^.

**Figure 2.**
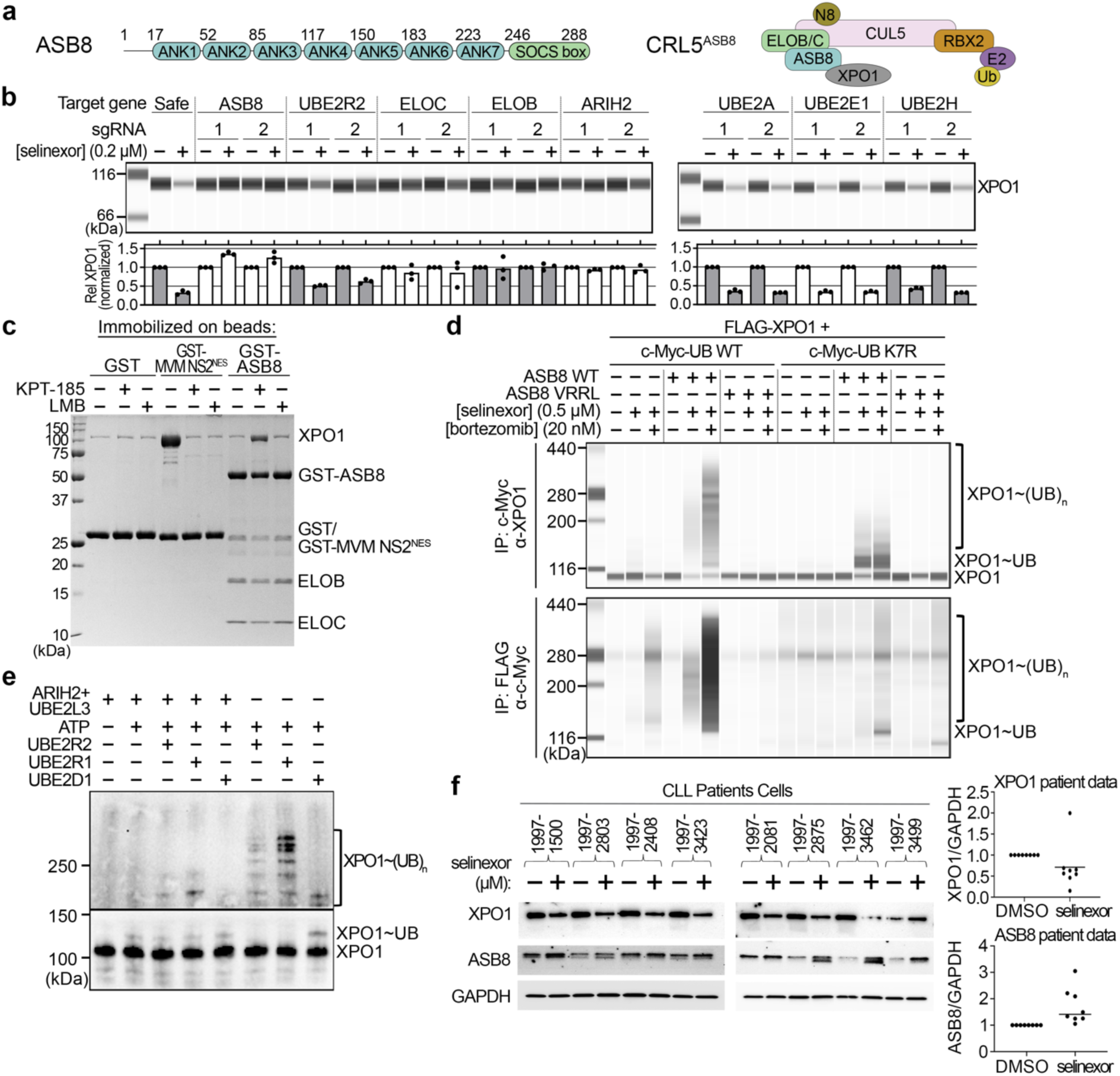
CRL5^ASB8^ binds and degrades selinexor/KPT-185-XPO1. **a**, Domain organization of ASB8 and schematic of the inhibitor-XPO1•CRL5^ASB8^ complex. **b**, Rescue of SINE-XPO1 degradation in HAP1 cells by CRISPR knockout of either a CRL5^ASB8^ component or E2 ligase, assessed by Simple Western. XPO1 levels were normalized to total protein content and plotted relative to untreated conditions. Results are from three independent experiments. **c**, Pull-down assay of immobilized GST-ASB8•ELOB/C (negative control, immobilized GST) with XPO1 pre-treated with inhibitors or DMSO showing that the compounds inhibit XPO1-NES interaction. Bound proteins are visualized by Coomassie SDS-PAGE, unbound proteins in **Extended Data Figure 2f**. **d**, Ubiquitination of XPO1 in HEK293T cells was assessed by immunoprecipitation of c-Myc (upper panel) or FLAG fusions (lower panel) from cell lysates, followed by Simple Western analysis using α-XPO1 (upper panel) or α-c-Myc (lower panel) antibodies. Additional blots in **Extended Data Figure 4a,b**. **e**, *In vitro* mono-ubiquitination and poly-ubiquitination of XPO1 by ARIH2/UBE2L3 and/or UBE2R2, UBE2R1 and UBE2D1, respectively (visualized by α-XPO1 in WB). **f**, Expression level of XPO1 and ASB8 by WB in HG3 cells (**Extended Data Figure 2g**) or CLL cells from patients treated with DMSO or 0.5 μM selinexor for 24 h. Quantitation of XPO1 and ASB8 protein levels in CLL patient cells shown on the right. Loading control is GAPDH.

Apart from ASB8, the involvement of other components and interactors of the CRL5^ASB8^ complex on SINE-induced XPO1 degradation had not been investigated. Here, we show that CRISPR and siRNA knockdowns of ELOB, ELOC, CUL5 and RBX2 all resulted in decreased SINE-XPO1 degradation, further confirming the role of CRL5^ASB8^ in this process (**Figure 2b**, **Extended Data Figure 2a-d**). We also investigated the roles of the ARIH2 E3 enzyme and several E2 enzymes identified by our previous CRISPR loss-of-function screens (**Figure 2b, Extended Data Figure 2a**). Indeed, knockout of ARIH2 rescues selinexor-induced XPO1 degradation suggestive of an E3-E3 ligase interaction^41,42^. Knockout of the E2 UBE2R2 partially rescues degradation, indicating a role in supporting XPO1 ubiquitination.

Previously, we co-immunoprecipitated XPO1 with ASB8 from cellular extracts in a selinexor dose dependent manner^7^. However, it was unclear whether ASB8 binds directly to selinexor-XPO1. To investigate this, we examined the direct interaction of SINE-XPO1 with a recombinant ASB8•ELOB/C complex. Immobilized GST-ASB8•ELOB/C successfully pulled down SINE-XPO1 but not LMB-XPO1 nor unliganded XPO1, demonstrating direct and specific binding of ASB8 to SINE-XPO1 (**Figure 2c**, **Extended Data Figure 2e,f**). Isothermal titration calorimetry (ITC) analysis reveals a dissociation constant (K*_D_*) of 29 [3.3, 88] nM for ASB8•ELOB/C binding to SINE-XPO1 (**Extended Data Figure 3a**).

Next, we investigated the ability of CRL5^ASB8^ to ubiquitinate XPO1 both in cells and *in vitro*. Co-transfection of myc-ubiquitin, FLAG-XPO1 and Tag100-ASB8 led to the appearance of higher molecular weight poly-ubiquitinated XPO1 protein bands, but only when cells were treated with selinexor and MG132 (**Figure 2d**, **Extended Data Figure 4a,b**). To confirm these bands were genuine poly-ubiquitinated XPO1, we included the ubiquitin K7R mutant, which is not capable of ubiquitin chain elongation, thus resulting primarily in mono-ubiquitinated protein. *In vitro* ubiquitination assays using recombinant neddylated CUL5•RBX2, ASB8•ELOB/C, ARIH2 and its E2 enzyme UBE2L3 successfully primed mono-ubiquitination of XPO1 in the presence of selinexor but not LMB (**Figure 2e**, **Extended Data Figure 4c,d**). E2 enzymes UBE2R2, UBE2R2 and UBE2D1 were able to poly-ubiquitinate selinexor-XPO1 to varying degrees, both with and without priming by ARIH2-UBE2L3 (**Figure 2e**).

Upon validating that CRL5^ASB8^ ubiquitinates SINE-XPO1 to promote its degradation, we examined selinexor-induced XPO1 degradation and ASB8 level in primary samples from patients with chronic lymphocytic leukemia (CLL) and in the CLL cell line HG3 (**Figure 2f, Extended Data Figure 2g**). Selinexor induces degradation of XPO1 in primary CLL cells and in HG3 cells (**Figure 2f, Extended Data Figure 2g**). A modest increase in ASB8 levels was observed following selinexor treatment, although the significance of this change remains unclear.

### CryoEM structures of SINE-XPO1 bound to the ASB8•ELOB/C complex

To understand how SINE-XPO1 engages CRL5^ASB8^, we assembled complexes of selinexor-XPO1 and KPT-185-XPO1 bound to the ELOB/C-bound ASB8^τιN16^ (N^helix^ removal decreased aggregation) for cryoEM structure determination. We solved the RAN^GTP^-bound selinexor-XPO1•ASB8•ELOB/C structure to 3.3 Å resolution and the KPT-185-XPO1•ASB8•ELOB/C structure to 3.4 Å resolution (**Extended Data Table 1** and **Extended Data Figures 5 and 6**). For comparison, we also solved cryoEM structures of unliganded and selinexor-bound XPO1 at 2.9 Å and 3.4 Å resolution, respectively. These structures are very similar to previously published crystal structures of unliganded XPO1 (PDB: 3VYC, 4BSM, 4FGV) and selinexor-XPO1 (6TVO and 7L5E)^28,43–46^ (**Extended Data Table 1** and **Extended Data Figures 7 and 8**).

The cryoEM maps for RAN^GTP^•selinexor-XPO1•ASB8•ELOB/C and KPT-185-XPO1•ASB8•ELOB/C are very similar. The only notable difference is RAN-binding to the N-terminal HEAT repeats of XPO1, which stabilizes them. These HEAT repeats are dynamic and not resolved in the KPT-185-XPO1•ASB8•ELOB/C map, due to the absence of the GTPase (**Figure 3a,b**). We first built the KPT-185-XPO1•ASB8•ELOB/C structure using the AlphaFold model of ASB8, the ASB9-bound ELOB/C structure (PDB: 6V9H)^37^ and the cryoEM structure of selinexor-XPO1 from this study (**Figure 3b**). Next, we built the RAN^GTP^•selinexor-XPO1•ASB8•ELOB/C structure using ASB8•ELOB/C from the KPT-185-XPO1•ASB8•ELOB/C structure, and XPO1 and RAN^GTP^ from previous crystal structures (6TVO, 3M1I)^26,47^ (**Figure 3a**).

**Figure 3.**
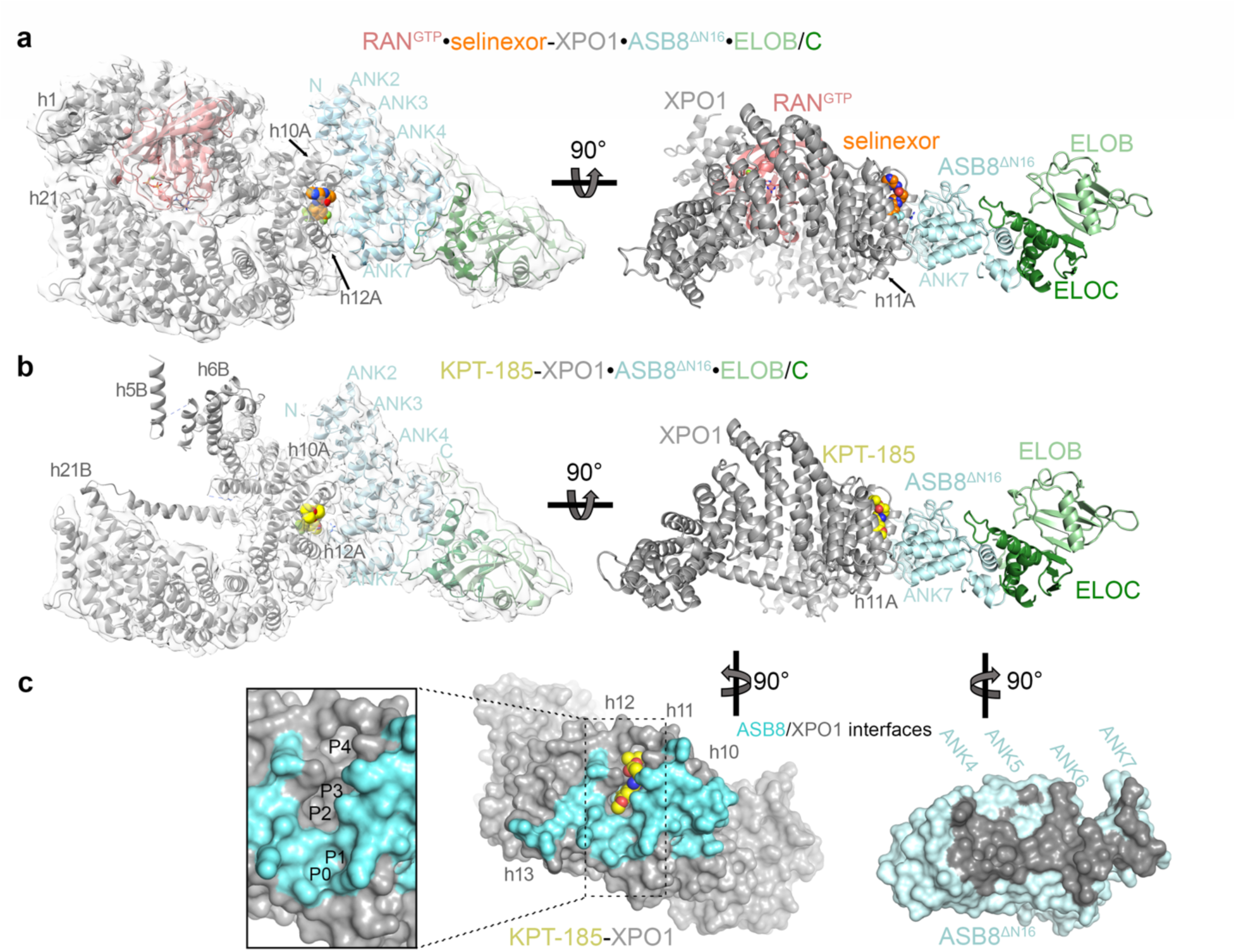
CryoEM structures of SINE-XPO1•ASB8 complexes. **a**, CryoEM map and refined structure of RAN^GTP^ (pink)•selinexor (orange spheres)-XPO1 (gray)•ASB8^ΔN16^ (cyan)•ELOB/C (light/dark green). **b**, CryoEM map and refined structure of KPT-185 (yellow spheres)-XPO1 (gray)•ASB8^ΔN16^ (cyan)•ELOB/C (light/dark green). **c**, XPO1 and ASB8 surfaces in grey and light cyan, respectively. Butterfly views show the ASB8 interface of XPO1 (cyan) and the XPO1 interface of ASB8 (dark grey). An insert shows the XPO1 groove with KPT-185 removed to reveal the P0-4 hydrophobic pockets.

ASB8•ELOB/C binds similarly to both selinexor-XPO1 and KPT-185-XPO1 (**Figure 3a,b**). The concave surface of the ASB8 ANK domain solenoid (ANK2-7) abuts the convex surface of XPO1 at HEAT repeats h10-h12, which includes the inhibitor-bound groove. Meanwhile, the ASB8 C-terminal SOCS box packs against the convex surface of its ANK5 and ANK6 repeats, similar to ASB9^37^ (**Figure 3a-c**, **Extended Data Figure 9a**). The ASB8 SOCS box binds the ELOB/C heterodimer in the same manner as the ASB9 SOCS binds ELOB/C. Additionally, alignment of the ASB8-bound CRL adaptor allowed the overlay of the RBX1 bound NEDD8-CUL5 (**Extended Data Figure 9b**)^37,42,48,49^. The two structures are very similar in the regions where selinexor-XPO1 and KPT-185-XPO1 bind to ASB8•ELOB/C, with Cα r.m.s.d. values of 0.7 Å for XPO1 central HEAT repeats h10-12 and 0.5 Å for ASB8•ELOB/C. The N– and C-terminal HEAT repeats of the two XPO1 solenoids differ slightly (Cα r.m.s.d. 1.9 Å) because of the RAN^GTP^ that is bound to the selinexor-XPO1•ASB8•ELOB/C complex (**Figure 3a,b**).

KPT-185 and selinexor bind XPO1 similarly regardless of whether ASB8•ELOB/C is bound. The map feature of the pyrazinylpropenehydrazide arm of selinexor is weak in the new selinexor-XPO1 cryoEM structure (**Extended Data Figure 8e**). The different orientations of the arm in the RAN^GTP^•selinexor-XPO1•ASB8•ELOB/C cryoEM structure (**Extended Data Figure 5d**) and the previously reported selinexor-XPO1 crystal structure (7L5E)^28^ suggest flexibility (**Extended Data Figure 8f**). In contrast, the isopropyl acrylate arm of KPT-185, which binds deeper in the XPO1 groove, is positioned in a similar manner in both cryoEM and crystal structures (4GMX)^24^ (**Extended Data Figures 6e**). The core triazole and substituted phenyl scaffolds of selinexor and KPT-185 bind the XPO1 groove similarly in the cryoEM and crystal structures. The SINE-bound XPO1 grooves in the cryoEM structures adopt the same open conformation as those in the crystal structures of XPO1 bound to the synthetic SINE inhibitors, natural product lactone polyketide inhibitors and NES peptides (**Extended Data Figure 10**).

### Interactions between ASB8 and the selinexor/KPT-185-bound XPO1

For structure analysis, we focus on the KPT-185-XPO1•ASB8•ELOB/C structure for interaction analysis because of its overall structure statistics (**Figure 4a,b, Extended Data Table 1**). The interface in the RAN^GTP^•selinexor-XPO1•ASB8•ELOB/C structure is very similar (**Extended Data Figure 11a,b**).

**Figure 4.**
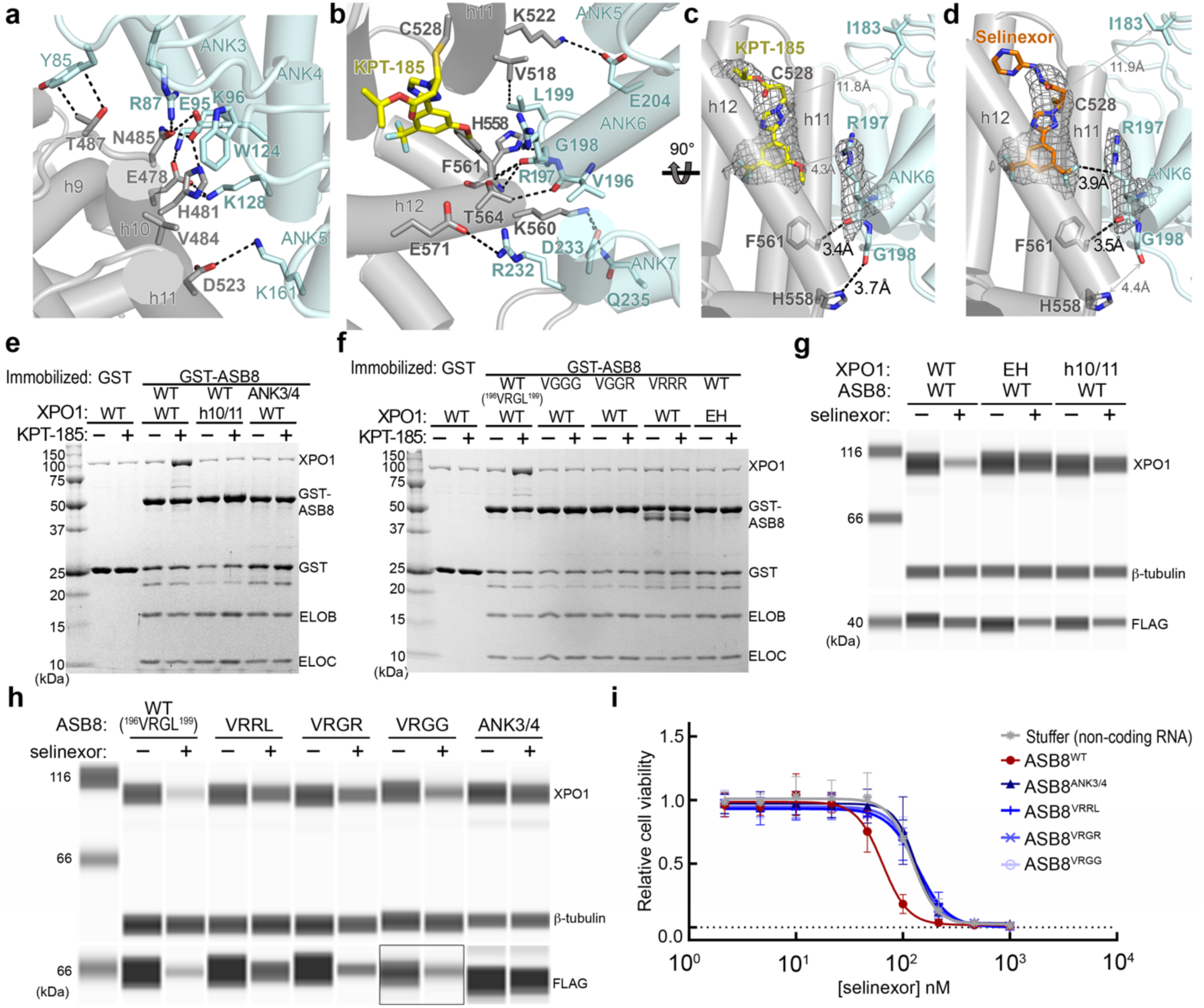
The SINE-XPO1-ASB8 interaction interface. **a**, The XPO1-ASB8 h10/11-ANK3/4 interface. Dashed lines show contacts <4.0 Å. XPO1 and ASB8 side chains with bolded labels were mutated to generate XPO1^h10/11^ and ASB8^ANK3/4^ mutants, respectively, for experiments in panel **(e)**. **b**, The XPO1-ASB8 h11/12-ANK6/7 interface. Dashed lines show contacts <4.0 Å. **c-d**, Side views of interfaces in **b** for the KPT-185-XPO1•ASB8•ELOB/C (**c**) and RAN^GTP^•selinexor-XPO1•ASB8•ELOB/C (**d**) structures. CryoEM maps are shown as gray meshes at the inhibitors and the nearby ASB8 R197. **e,f**, Pull-down assay of immobilized GST-ASB8•ELOB/C with XPO1 and KPT-185 or DMSO, visualized by Coomassie staining. Mutations of ASB8 or XPO1 target the h10/11-ANK3/4 (**e**) and the h11/12-ANK6/7 interfaces (**f**). Unbound proteins shown in **Extended Data Figure 11c**. **g**, HEK293T cells were transfected with XPO1 WT, XPO1^EH^ or XPO1^h10/11^ alone (**Extended Data Figure 11e**) or together with FLAG-tagged ASB8 WT and treated overnight with selinexor; XPO1 degradation assessed by Simple Western. Additional data shown in **Extended Data Figure 11e**. **h**, HEK293T cells were transfected with FLAG-tagged ASB8 WT (^196^VRGL^199^), ASB8^ANK3/4^ and ASB8 mutants VR<u>R</u>L, VRG<u>R</u> or VRG<u>G</u> followed by selinexor treatment overnight and Simple Western analysis of XPO1. Boxed bands indicate samples blotted separately. Representative WB images in **g** and **h** (loading control, β-tubulin). Additional data shown in **Extended Data Figure 11f**. **i**, Cell viability assay on HAP1 cells with stable overexpression of ASB8 WT or one of the ASB8 mutants. Data points were normalized to untreated cells and represent mean ±SD of four separate experiments.

ASB8 covers 993 Å^2^ of XPO1 spanning h10-h12 and ASB8-XPO1 contacts are concentrated at two ends of this interface (**Figure 3c**). At one end, XPO1 h10 and h11 (h10/11) make numerous contacts with ANK repeats 3 and 4 (ANK3/4) of ASB8 (**Figure 4a**). At the other end, the XPO1 groove (h11/12) interacts with ANK repeats 6 and 7 (ANK6/7) (**Figure 4b**). Most of the KPT-185 molecule bound in the XPO1 groove is far from ASB8, and the methoxy group of KPT-185 is 4.3 Å away from the ASB8 R197A side chain at the nearest point (**Figure 4c**). Similarly, most of the conjugated selinexor is distant from ASB8, with only one putative contact (3.9 Å) observed between one of selinexor’s trifluoromethyl (TFM) group and the ASB8 R197 side chain (**Figure 4d**).

The interface between XPO1 h10/11 and ASB8 ANK3/4 involves a variety of polar, electrostatic, and hydrophobic interactions (**Figure 4a**). We mutated six XPO1 residues at this interface (E478A, H481A, V484G, N485G, T487G, and D523G; named XPO1^h10/11^ mutant), which substantially decreased KPT-185 dependent binding to GST-ASB8•ELOB/C (**Figure 4e**). Similarly, mutation of the ASB8 contact residues (R87A, E95A, K96A, W124A, and K128A; named ASB8^ANK3/4^ mutant) similarly decreased binding to KPT-185-XPO1.

At the other XPO1-ASB8 interface, the ASB8 ANK6/7 repeats bind part of the XPO1 groove (**Figure 4b**). The XPO1 groove is lined with five hydrophobic pockets P0-4 (**Figure 3c**), and SINE inhibitors bind across the P2-P4 pockets. ASB8 residues^196^ VRGL^199^ (at the last helix turn of the ANK6 α1 helix and the ensuing loop) interacts with XPO1 pockets P0 and P1 (**Figure 4b**). The ASB8 L199 side chain binds the XPO1 P1 pocket while the main chain carbonyl groups of ASB8 ^196^VRG^198^ make multiple contacts with the main and side chains of the XPO1 h12A helix (**Figure 4b-d**). Nearby, ASB8 ANK7 residues R232 and D233 make electrostatic interactions with XPO1 K560 and E571, respectively (**Figure 4b**). E571 is an oncogenic mutation site^50–53^. Mutations of XPO1 V518 and F561 within the ASB8-binding site to Asp and His, respectively (mutant named XPO1^EH^), transformed the hydrophobic P0 and P1 pockets to a charged surface, and substantially reduced ASB8 binding (**Figure 4f**). Mutations of ASB8 ^196^VRGL^199^ to VGGG, VGGR or GRGG, to destabilize the last helical turn of the ANK6 α1 helix, or to VRRR to generate steric clash with the XPO1 P1 pocket, also decreased ASB8-XPO1 interactions (**Figure 4f**, **Extended Data Figure 11c,d**).

All these interface mutations also reduced selinexor-induced XPO1 degradation in cells, indicating that the interaction with ASB8 is crucial for this process. Indeed, transient overexpression of ASB8 and XPO1 wild type (WT) combined with selinexor treatment, typically leads to extensive XPO1 degradation but mutations at the XPO1-ASB8 interface impair this degradation ability (**Figure 4g,h, Extended Data Figure 11e,f**). Moreover, while overexpression of ASB8 WT sensitizes cells to selinexor, this sensitizing effect is absent in cells expressing any of the mutant ASB8 proteins (**Figure 4i**). Collectively, the structural, biochemical, and cellular results above reveal the importance of interactions between inhibitor-bound XPO1 h10/11 with ASB8 ANK3/4 and between the P0 and P1 pockets of the KPT-185-bound XPO1 groove with ASB8 ANK6 residues ^196^VRGL^199^ for engagement of inhibitor-XPO1 by CRL5^ASB8^ for XPO1 degradation.

### Selinexor/KPT-185 reveals a cryptic ASB binding site in the XPO1 groove

The XPO1 interface with ASB8 is a three-dimensional degron that spans the convex surface HEAT repeats h10-h12 (**Figure 3a-c**). This conformation at h10/11 is not unique to selinexor/KPT-185-bound XPO1 since XPO1 repeats h10 and h11 adopt the same conformation in many different XPO1 structures; Cα r.m.s.d. of residues 469-552 is 0.6 – 1.0 Å when comparing unliganded, NES-, inhibitor-, RANBP3– and nucleoporin-bound structures (**Figure 5a**)^25,54–56^. The XPO1 side chains that contact ASB8 ANK3/4 are all solvent-accessible in the different structures. The ASB8-binding site at h10/11 is thus not unique to selinexor/KPT-185-bound XPO1. However, the XPO1 groove, which is made up of HEAT repeats h11 and h12, is different depending on the bound ligand (**Figure 5a-f**). Our cryoEM structure of unliganded human XPO1 and three crystal structures of unliganded XPO1 from several organisms all show that the grooves are closed (**Figure 5b, Extended Data Figure 12**). The ASB8-binding site is thus inaccessible to unliganded XPO1.

**Figure 5.**
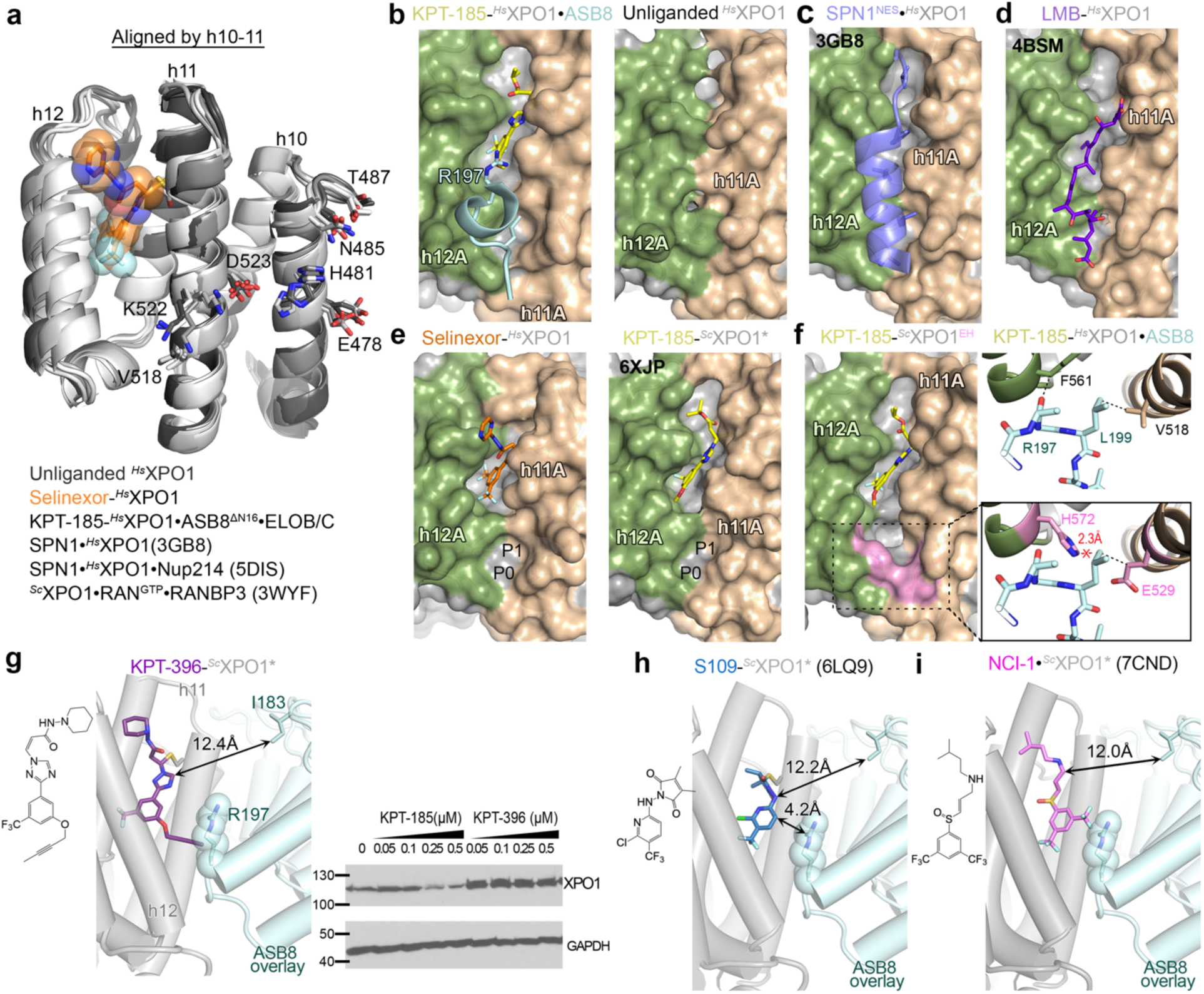
The ASB8 binding site. **a**, Repeats h10-12 XPO1 of several structures drawn as cartoons and aligned at h10/11. Selinexor (spheres) and h10/11 side chains that contact ASB8 are drawn as sticks. **b**, The left panel shows the NES/inhibitor-binding groove of KPT-185•XPO1•ASB8 is open and occupied by KPT-185 and ASB8. The different colors of helices h11A (brown), h12A (green) and behind them, h11B and h12B (grey) show the open *vs*. closed grooves. Panels **c-f** use the same orientation and colors. The right panel shows the unliganded XPO1 groove, which is closed (additional views in **Extended Data Figure 11**). **c-e**, XPO1 grooves bound to SPN1^NES^ (blue; 3GB8) (**c**), LMB (purple; 4HAT) (**d**), selinexor (**e**, left) and KPT-185 (6XJP) (**e**, right). **f**, The KPT-185-bound groove of the *^Sc^*XPO1^EH^ mutant. An insert shows overlayed ASB8 and the likely steric clash, compared to KPT-185-*^Hs^*XPO1•ASB8•ELOB/C above. The sequence alignment of human and yeast XPO1 shown in **Extended Data Figure 15a**. **g**, The chemical structure of KPT-396, and the KPT-396-bound XPO1 structure with ASB8 overlayed. XPO1 level when cells were treated with DMSO, Selinexor or KPT-396; XPO1 detected by WB and GAPDH used as loading control. **h-i**, Same as in **g**, showing S109 (6LQ9) (**h**) and NCI-1 (7CND) (**i**).

The cargo-bound XPO1 groove is occupied by the cargo’s NES (**Figure 5c**). These NES peptides have diverse sequences and adopt a variety of conformations when binding to the XPO1 groove, but all characterized NESs minimally bind across the P1-P3 pockets (**Figure 5c**, **Extended Data Figure 13**)^57,58^. They all occupy the ASB8-binding site, blocking it; thus, XPO1-NES binding is incompatible with ASB8 binding. LMB and other lactone polyketide inhibitors also occupy the entire XPO1 groove leaving no room for ASB8 to bind, which aligns with LMB not inducing XPO1 degradation (**Figure 5d**)^25,30^.

SINE compounds like selinexor and KPT-185 are smaller than LMB. SINE compounds make many hydrophobic interactions with the open XPO1 groove (**Extended Data Figure 14a,b**), but they occupy only P2-P4^20,24,26–28^, leaving the P0 and P1 pockets exposed (**Figure 5e**). The P0 and P1 pockets, along with the nearby XPO1 residues at the N-terminal ends of helixes h11A and h12A, form the three-dimensional degron unique to the SINE-bound exportin.

Perturbation of this XPO1 surface, such as the XPO1^EH^ mutation, abrogated SINE-induced XPO1 degradation (**Figure 4f,g**). The crystal structure of the KPT-185 bound *Sc* equivalent of the XPO1^EH^ mutant (*^Sc^*XPO1^EH^ with V529E/F572H mutations equivalent to V518E/F561H of *^Hs^*XPO1) shows a polar/charged groove with shallower P0 and P1 pockets that are not compatible with ASB8 binding (**Figure 5g, Extended Data Figure 15a-d** and **Extended Data Table 2**). When we utilized SINE analogs with larger phenyl substituents, the ASB8 binding site was also perturbed. KPT-396, containing a larger oxypropyne substituent than selinexor’s TFM, is unable to induce XPO1 degradation (**Figure 5g**). A 2.4 Å resolution crystal structure of KPT-396-XPO1 shows the flexible KPT-396 oxypropyne (no electron density) extending toward solvent where it would clash with the ASB8 R197 side chain (**Figure 5g**, **Extended Data Figure 15e-g** and **Extended Data Table 2**).

We also aligned the KPT-185-XPO1•ASB8•ELOB/C structure with those of XPO1 bound to the SINE eltanexor^20^ and two chemically completely unrelated XPO1 inhibitors, felezonexor analog S109 and the non-covalent inhibitor NCl-1^59^ (**Figure 5h,i, Extended Data Figure 16a**). S109 and eltanexor do not seem clash with the overlayed ASB8, consistent with their previously reported abilities to induce XPO1 degradation. Pull down assays demonstrate that felezonexor– and eltanexor-XPO1 indeed bind ASB8•ELOB/C **(Extended Data Figure 16b)**. NCI-1 has not been reported to cause XPO1 degradation^55^. Structural alignment reveals a clash with ASB8 suggesting it might not trigger CRL5^ASB8^-mediated XPO1 degradation (**Figure 5i**).

Although the ASB8-binding site at one end of the XPO1 groove is clearly unique to selinexor/KPT-185-bound XPO1 (**Figure 5b,e**), the adjacent site at h10/11 remains conformationally invariant across all known XPO1 states, suggesting that the exportin should have low affinity for ASB8, even without inhibitors, when the XPO1 groove is closed. Indeed, ITC reveals a K*_D_* of 11 μM for ASB8-XPO1 binding in the absence of SINE (**Extended Data Figure 3b**), but this is too weak for productive ubiquitination and degradation^60^.

### SINEs induce ASB8-binding via an allosteric mechanism

All SINE compounds bind covalently to XPO1 and engage in extensive non-covalent interactions with it (**Figure 4c,d, Extended Data Figure 14**). In contrast, KPT-185 does not contact ASB8 while selinexor makes a single van der Waals contact with the ASB8 R197 side chain. This mode of XPO1-degrader-E3 ligase interaction is substantially different from those of characterized molecular glue degraders that bridge substrate and E3 ligase through multiple direct contacts to both proteins (**Extended Data Figure 17**)^8–10,61–68^. SINEs primarily function by indirectly or allosterically opening the XPO1 groove, thereby exposing a distinct cryptic site that binds ASB8 (**Figure 6a**).

**Figure 6.**
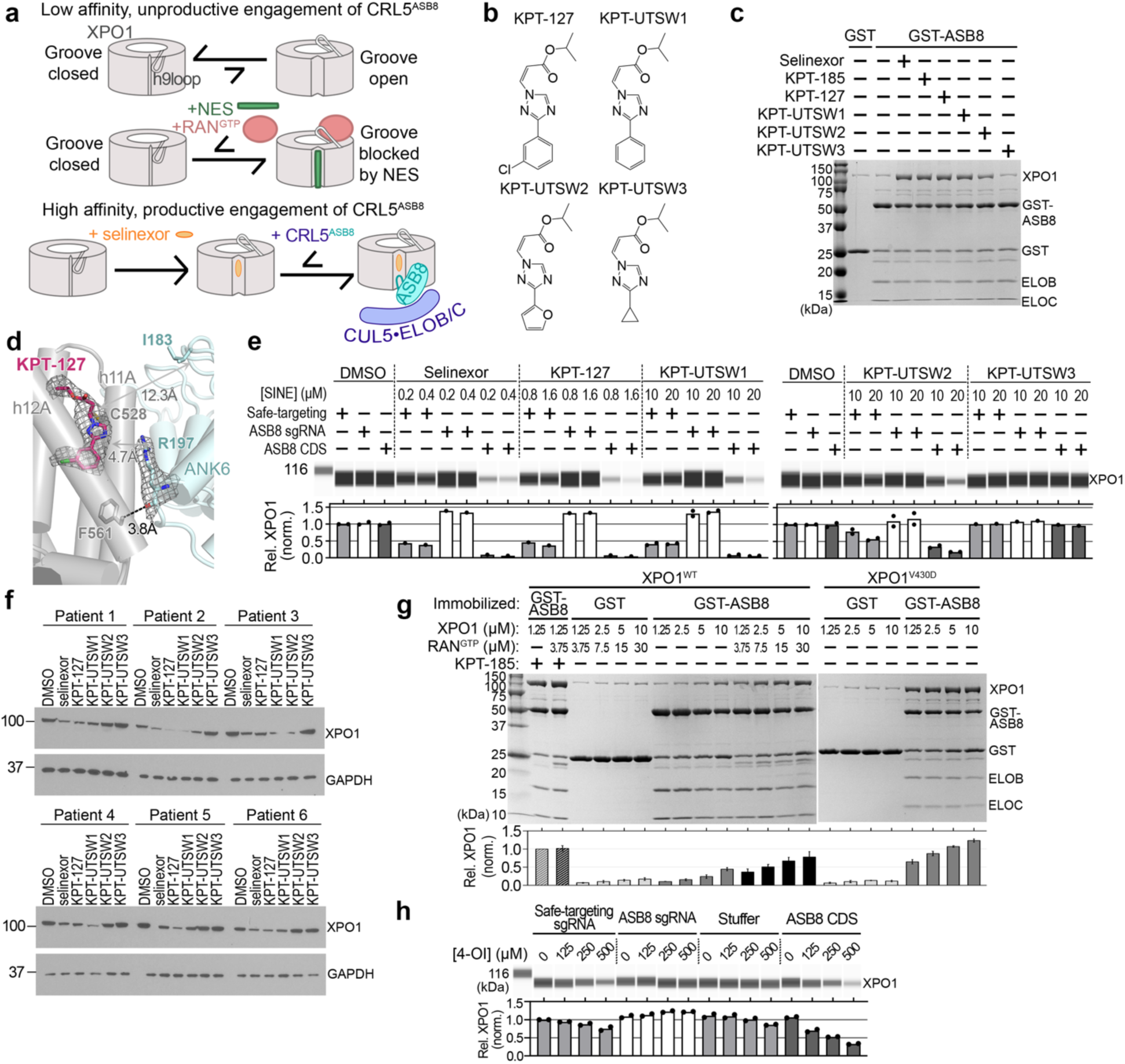
SINEs induce XPO1 degradation via an allosteric mechanism. **a**, The ASB8 binding site at the XPO1 groove is occluded when 1) XPO1 is unliganded and the closed groove is stabilized by the h9 loop, or 2) the groove opens and binds NES as RAN^GTP^ disengages the h9 loop. XPO1s with blocked ASB8 binding site have low affinity and unproductive engagement of CRL5^ASB8^. In contrast, selinexor binds covalently to occupy half of the XPO1 groove, keeping it open and exposing a cryptic ASB8 binding site to produce high affinity productive engagement of CRL5^ASB8^. **b**, Chemical structures of SINE analogs that are smaller than selinexor/KPT-185. **c**, Pull-down assays to assess different SINE-XPO1 complexes binding to ASB8. Unbound proteins in **Extended Data Figure 18c**. **d**, CryoEM structure of KPT-127-XPO1•ASB8^ΔN16^•ELOB/C. Map density of KPT-127 and ASB8 R197A is shown as gray mesh. **e**, XPO1 degradation in HAP1 cells by minimal SINE analogs. Cells were treated with with twofold and fourfold the IC_50_ (as determined in a 72-hour cell viability assay, see **Extended Data Figure 18f**), with a maximum of 10 µM and 20 µM. Representative Simple Western blots are shown. Quantitative analysis from two independent experiments. **f**, XPO1 degradation in primary CLL cells from patients were treated with 0.5 uM selinexor or 1 uM KPT-127, KPT-UTSW1, KPT-UTSW2 or KPT-UTSW3 for 24 hours. Whole cell lysates were prepared 24 hours post-treatment. Loading control is GAPDH. **g**, Pull-down assays to assess binding of ASB8 to XPO1 without SINE, via allosteric activation by RAN^GTP^ or the V430D mutation in the h9 loop. Quantification of band intensity from N = 3 experiment is plotted below as mean ± s.d. **h**, Degradation of XPO1 by 4-OI in HAP1 cells. Cells were treated overnight with 4-OI followed by lysis and immunodetection of XPO1. XPO1 levels were normalized to total protein content and plotted relative to lane 1. Data collected from two independent experiments. Additional data shown in **Extended Data Figure 23**.

We employed two approaches to investigate the allosteric mechanism of the SINE degraders. First, we utilized small SINE analogs to confirm that, upon SINE binding to XPO1, ASB8 interacts with XPO1 independently of any interaction with compound. Second, we examined ASB8’s ability to bind to XPO1 stabilized in the open-groove conformation, without inhibitors. KPT-127 contains only one substituent on its phenyl^69,70^ while KPT-UTSW1 has no phenyl substituent; KPT-UTSW2 contains a furanyl instead of the phenyl group and KPT-UTSW3 has a cyclopropyl instead of the phenyl group (**Figure 6b,c**). Compared to KPT-185 or selinexor, KPT-127 exhibits similarly efficient conjugation kinetics, whereas KPT-UTSW1 is slower but still successfully conjugates to XPO1 after overnight incubation (**Extended Data Figure 18a**). Both KPT-127-XPO1 and KPT-UTSW1-XPO1 complexes successfully pulled down ASB8, induced mono-ubiquitination by ARIH2-UBE2L3 *in vitro*, and induced XPO1 degradation in cells (**Figure 6c,e, Extended Figures 4d** and **18d**). CryoEM structures confirmed no SINE-ASB8 contacts (**Figure 6d, Extended Data Figures 19 and 20**). These results clearly demonstrate that interaction with the compound is not required for ASB8 binding to liganded XPO1 or for its degradation activity.

The two smallest SINE analogs, KPT-UTSW2 and KPT-UTSW3, did not effectively conjugate XPO1 even after overnight incubation (**Extended Data Figure 18b**). However, despite inefficient conjugation, KPT-UTSW2 can enhance XPO1-ASB8 interaction and induce XPO1 degradation in cells (**Figure 6c, Extended Data Figure 18d**). We suspect that although both KPT-UTSW2 and ASB8 each bind weakly to XPO1, they act with positive cooperativity enhancing each other’s interaction with XPO1.

We also took a different approach to remove the one selinexor-ASB8 contact (**Figure 4d**) by mutating ASB8 R197 to an alanine (ASB8 R197A mutation). ASB8 R197A binds selinexor-ASB8 with high affinity (ITC, K*_D_* 58 [7.8, 225] nM, **Extended Data Figure 3c**). However, the cryoEM structure of selinexor-XPO1•ASB8^R197A^•ELOB/C yielded inconclusive results due to major rearrangement of the inhibitor (**Extended Data Figure 21**).

Next, we leveraged XPO1’s intrinsic allostery to investigate the allosteric effect of SINEs on ASB8 binding. RAN^GTP^ binds at the N-terminal HEAT repeats of XPO1. RAN^GTP^-binding shifts the equilibrium of the exportin from a closed to an open groove conformation (**Figure 1a,b**), which enhances NES binding, and we hypothesize this also increases ASB8 binding. Indeed, XPO1/RAN^GTP^ titration and pull-down by immobilized GST-ASB8•ELOB/C (with ITC being in the presence of RAN^GTP^) showed greater ASB8 recruitment compared to XPO1 alone (**Figure 6g, Extended Data Figure 22a**). To further confirm the ability of ASB8 binding to the XPO1 open groove without compound, we stabilized XPO1 in its open groove conformation by introducing the well-characterized V430D (V441D in yeast) mutation in its h9 loop. The h9 loop of XPO1 stabilizes the closed groove by interacting with its h11B and h12B helices (**Figure 1b**, **Extended Data Figure 7d**); V430D disrupts this interaction and shifts XPO1 to its open groove state and enhances NES-binding^47,58^. XPO1 V430D effectively pulls down ASB8 without compound (**Figure 6g** and **Extended Data Figure 22a,b**).

Altogether, these results demonstrate that direct contact between SINEs and ASB8 is not required for high affinity formation of the SINE-XPO1•CRL5^ASB8^ complex and subsequent degradation of XPO1, strongly supporting an allosteric mechanism of action. Groove opening is key for ASB8 recruitment. Therefore, upon SINE binding the NES-binding groove is stabilized in its open conformation. This conformational change exposes a cryptic ASB8-binding site at the opposite end of the SINE binding site and facilitates high affinity ASB8 interaction with XPO1 and hence degradation.

### Cellular ligands and CRL5^ASB8^-mediated XPO1 degradation

It has been reported that secondary metabolites such as prostaglandins, itaconate and likely other metabolites bind to Cys528 in the XPO1 groove^71–74^. Also, nitric oxide can modify the XPO1 groove via S-nitrosylation of Cys528^75^. However, the biological role for these modifications is not known. We hypothesize that these modifications modulate XPO1 activity. Indeed, XPO1 transports numerous cargo proteins with diverse functions and maintaining proper XPO1 levels and function ensures that these proteins are correctly localized which is vital for cellular homeostasis. Therefore, it is reasonable to consider that biological mechanisms regulate XPO1 protein level. We have previously reported anti-viral effects of the secondary metabolite itaconate and demonstrated that the itaconate derivative 4-octyl itaconate (4OI) blocks nuclear export of viral RNP by inhibiting XPO1^72^. Itaconate is a metabolite produced in the tricarboxylic acid cycle and plays an important role in macrophages upon infection and inflammation. Here we demonstrate that 4OI-induced XPO1 degradation is ASB8 dependent (**Figure 6h** and **Extended Data Figure 23)**. This result suggests that targeted ASB8-mediated degradation is a native biological process that is exploited by SINE.

### Therapeutic implications

Many studies have shown that KPT-SINEs effectively block XPO1-NES/cargo binding and nuclear export^7,24,29,76^. Nuclear export inhibition occurs rapidly, within 30 minutes of compound addition, whereas XPO1 degradation starts approximately 6 hours after treatment^4^, suggesting that the primary driver of antitumor activity of the compounds is inhibition of nuclear export. Additional support for this model comes from the reversible XPO1 inhibitor FR-027, which potently inhibits nuclear export but does not induce XPO1 degradation and still demonstrates potent antitumor efficacy^77^. This suggests that while degradation may contribute additively to antitumor activity, this role may be non-essential when nuclear export inhibition is potent. Preclinical studies, including those in Multiple Myeloma (MM), further support this concept. These studies show that proteasome inhibitors such as bortezomib or carfilzomib can be combined with selinexor without compromising nuclear export inhibition or its associated antitumor effects^78–80^.

Although the contribution of selinexor-induced XPO1 degradation to antitumor activity is secondary, excessive XPO1 depletion may contribute to systemic toxicity and increase adverse effects in patients. Over-depletion of XPO1 from healthy cells due to SINE-induced XPO1 degradation may impair the recovery of normal tissues between dosing. This hypothesis warrants future investigation.

In conclusion, although SINEs primarily exert their antitumor activity through inhibition of cargo binding rather than degradation, the role of CRL5^ASB8^-mediated XPO1 degradation of XPO1 should be considered in future drug design. Although our understanding of the CRL5^ASB8^-complex and its regulation of XPO1 is still emerging, understanding the allosteric mechanism of XPO1’s degraders provide a new entry point for modulating XPO1 activity in therapeutic contexts. Targeting XPO1’s nuclear export function through direct inhibition or protein degradation presents diverse therapeutic avenues. Optimal strategy may include degraders that reduce XPO1 protein levels without impairing export, inhibitors that selectively block export without promoting degradation, or dual-action – each offering potential to balance efficacy with reduced toxicity.

## Acknowledgements

We thank the Structural Biology Laboratory and the CryoEM Facility at UTSW (supported by the Cancer Prevention & Research Institute of Texas or CPRIT RP220582) as well as the Pacific Northwest Center for Cryo-EM (PNCC; with assistance from Sean Mulligan; supported by NIH grant R24GM154185) for cryoEM data collection, and the Advanced Photon Source, operated for the DOE Office of Science by Argonne National Laboratory under Contract No. DE-AC02-06CH11357 for X-ray crystallography data collection. We thank Lotte Bral, Eva Libaers and Kim Janssens for excellent technical support. We thank H. Niederstrasser and B. A. Posner of the High-Throughput Screening Core Facility of UTSW (National Institutes of Health (NIH) shared instrumentation grant 1S10OD018005-01 to B.A.P. and the Simmons Cancer Center Support grant – Arteaga 2P30CA142543-11) for siRNA analysis. We thank Ning Zheng for advice on performing *in vitro* ubiquitination assays and the Elizabeth Komives laboratory for gifting plasmids necessary for overexpression of ASB8 and other proteins necessary for the *in vitro* ubiquitination assays. We are grateful to the patients who provided blood for the above studies and to the OSU Comprehensive Cancer Center Leukemia Tissue Bank (supported by NCI P30 CA016058) for sample procurement. This work was funded by the National Institute of General Medical Sciences of the NIH under R35GM144137 (Y.M.C.) and the Molecular Biophysics Training Program T32GM131963 (C.E.W.), the National Cancer Institute of the NIH under R01 CA214046 (R.L.), the Welch Foundation grant I-1532 (Y.M.C.), CPRIT grants RP180410, RP150053 and RP170170 (Y.M.C.), support from the Alfred and Mabel Gilman Chair in Molecular Pharmacology (Y.M.C.), Eugene McDermott Scholar in Biomedical Research (Y.M.C.), the FWO (Fonds Wetenschappelijk Onderzoek) grant G0E6517N (D.D), the HHMI International Fellowship (H.Y.J.F.) and the Gilman Special Opportunities Award (H.Y.J.F.).

## Author Contributions

The project was conceived by Y.M.C with inputs from T.C., H.Y.J.F. and C.E.W.. The manuscript was written by Y.M.C., C.E.W. and H.Y.J.F. with inputs from D.D., B.K. and R.L. Cellular experiments were performed T.G., and biochemical experiments by C.E.W., A.B.N. and B.S.. CryoEM and crystal structures were determined and analyzed by C.E.W. and H.Y.J.F.. Cellular experiments involving XPO1 inhibition, ubiquitination and degradation were performed by B.K., M.J. and B.P., with input from D.D.. Cellular experiments with CLL cells were performed by M.G., with input from R.L.. <u>KPT-UTSW1, KPT-UTSW2 and KPT-UTSW3 were synthesized by R.N., with input from J.R.R..</u> KPT-185, selinexor, KPT-396 and KPT-127 were provided by Karyopharm Therapeutics, with input by Y.L. and S.S.

## Supplementary Materials

Extended Data Figures 1 to 23

Extended Data Tables 1 to 3

Supplemental Methods

## Methods

### Constructs and expression vectors

Full-length wild type XPO1 was PCR amplified and subcloned into either into pciNeo-3xFLAG (gift from Dr. Xiaodong Wang, National Institute of Biological Sciences, Beijing) or into pTagRFP-C (Evrogen) mammalian expression vector to express ^FLAG^XPO1 or ^RFP^XPO1 fusion protein, respectively. XPO1 point mutant C528T variant was constructed by site directed PCR mutagenesis using the pciNeo-^FLAG^XPO1 construct. In addition, full-length attB-flanked ^FLAG^XPO1 was generated by PCR amplified from pCI-neo-^FLAG^XPO1 construct and was introduced into pDONR233(Thermo Fisher Scientific) to create an entry vector for LR Gateway Reaction (Thermo Fisher Scientific). Subsequently pDONR233-^FLAG^XPO1 and destination vector pcDNA5/FRT/TO vector (Thermo Fisher Scientific) were used in LR reaction (Thermo Fisher Scientific) as manufacturer’s instructions to generate pcDNA5/FRT-^FLAG^XPO1 FRT integration vector. Ubiquitin B (UBB) in pENTR221 vector was purchased as an Ultimate ORF clone (Thermo Fisher Scientific, clone ID IOH56688), PCR amplified and inserted into pcDNA3.1-HA (gift from Oskar Laur (Addgene plasmid # 128034; http://n2t.net/addgene:128034; RRID:Addgene_128034)) to express ^HA^Ub fusion protein. All constructs were made using standard molecular procedures and validated by direct Sanger sequencing.

### Cell culture and transfection

Human fibrosarcoma HT1080 were grown in complete Dulbecco’s modified Eagle’s medium (D-MEM, Thermo Fisher Scientific) supplemented with 10% heat inactivated fetal calf serum (FCS) penicillin/streptomycin (100 U/ml) and 2 mM glutamine). Flp-In T-REx-293 Cell Line (Thermo Fisher Scientific) was maintained in complete D-MEM supplemented with 15 μg/ml blasticidin and 100 μg/ml zeocin (InvivoGen). HEK 293 T-REx ^FLAG^XPO1 stable cell line maintained in the complete D-MEM culturing medium supplemented with 10 µg/ml blasticidin and 50 µg/ml hygromycin (InvivoGen). All transfections were carried out using either Lipofectamine2000 (plasmid) or Lipofectamine RNAiMAX (siRNA) according to manufacturer’s instructions.

HEK293T (ATCC) and HeLa Kyoto (Gifted by Prof. Johan Van Lint, KU Leuven) were cultivated in DMEM (Gibco) supplemented with 10% FCS and 20 µg/mL gentamicin. HAP1 (Horizon Discovery) cells were grown in IMDM (Gibco) supplemented with 5% FCS and 20 µg/mL gentamicin. Cells were cultivated at 37 °C and 5% CO_2_.

### Immunoblotting

Briefly, samples of proteins were prepared using either RIPA buffer (25 mM Tris-HCl, pH 7.5, 150 mM NaCl, 1% NP-40, 1% sodium deoxycholate and 0.1% SDS and 1x cOmplete protease inhibitors (Roche) or NET-150 (150 mM NaCl, 20 mM Tris at pH.8.0, 1 mM EDTA, 1x NaF/Na-orthovanadate (10 mM/ 1mM), 0.5 % NP-40, 1x cOmplete protease inhibitors). 10 µg portion of total lysates were resolved by SDS-PAGE and transferred to a PVDF membrane. The membrane was blocked for 1 h in 5% non-fat milk and incubated with primary antibody for another 1 h at room temperature. The membranes were then incubated with horseradish peroxidase-conjugated anti-mouse or anti-rabbit IgG (1:10,000; Thermo Fisher Scientific). Finally, the membranes were developed using a Kodak medical X-ray processor (Kodak, Rochester, NY, USA) or Gel Doc XR+ Gel Documentation System (Promega).

### Generation of the HEK 293 T-REx ^FLAG^XPO1 stable cell line

Flp-In T-REx-293 cells (Thermo Fisher Scientific) were co-transfected with Flp recombinase expression vector pOG44 (Thermo Fisher Scientific) and pcDNA5/FRT-^FLAG^XPO1 or pcDNA5/FRT construct. To select stable HEK 293 T-Rex ^FLAG^XPO1 cell lines derived from the host cell line by FLP mediated integration into the FRT docking site, zeocin was substituted with 50 μg/ml hygromycin B. ^FLAG^XPO1 expression was induced by supplementing growth media with 0.5µg/ml doxycycline (Sigma Aldrich).

### Pulse-chase labeling of endogenous XPO1 protein in HT1080 cells

HT1080 cells were grown 50% confluency and incubated with high glucose, no glutamine, no methionine, no cystine DMEM (Gibco, #21013024) supplemented with 1x L-glutamine (Gibco, #35050061) for 2 h. The cells were then pulsed for 1 h with 200 μCi/ml of [35S]-Met(methionine)/[35S]-Cys(Cysteine) (Perkin Elmer, NEG072002MC) and chased for 0, 12, 24, 27, 30, 33, 36 and 48 h. After each chase period the cells were lysed in RIPA buffer and XPO1 protein was recovered from whole lysates by immunoprecipitation using mouse monoclonal XPO1 antibody (Santa Cruz Biotech, sc-5595) and subjected to SDS-polyacrylamide gel electrophoresis. Gel was dried for 2 h and exposed to 35S intensifying screen and X-OMAT film in 70 °C for 24 h. The band corresponding to XPO1 was identified by autoradiography, and band intensities were calculated using ImageJ^81^.

### *In vivo* XPO1 degradation by SINE compound treatment

Human fibrosarcoma HT1080 cells and tetracycline-inducible HEK 293 T-REx ^FLAG^XPO1 cells were cultured in complete D-MEM and grown to 80% confluence. Where indicated, HT1080 cells were transiently transfected with the indicated RFP or FLAG-*^Hs^*XPO1 constructs and recovered in fresh media for 12 h before KPT-185 treatment. Unless otherwise indicated, cells were treated with 500 nM KPT-185, 1 µM selinexor or 10 nM LMB, and cells were grown for another 18-72 h. When needed MG132 or MLN4924 was added to growth media to 200 µM or at 100 µM final concentration. Cells were collected by centrifugation and lysed on ice for 30 min in RIPA buffer. Immunoblotting was used to detect XPO1 (CRM1 rabbit polyclonal H-300; sc-5595, Santa Cruz or CRM1 mouse monoclonal (C-1): sc-74454, Santa Cruz) and β-actin or β-tubulin (Loading Control Antibody Sampler Kit #5142, Cell Signaling Technology) to confirm equal protein loading.

### Detection of ^HA^UB-XPO1 conjugation

HEK 293 T-REx ^FLAG^XPO1 cells were reverse transfected with pcDNA3.1-^HA^UB or pcDNA3.1-HA and grown in complete D-MEM in the presence of doxycycline. Cells were then treated with DMSO, 0.5 µM KPT-185 and 200 µM MG132, or 200 µM MG132 alone for additional 18 h before lysis. Cells were collected by centrifugation and lysed on ice for 30 min in RIPA buffer. ^HA^UB∼^FLAG^XPO1 conjugation and total ^FLAG^XPO1 proteins were immunodetected by using primary antibodies for HA tag (Thermo Fisher Scientific, #26183) or FLAG M2 tag (Sigma Aldrich, F3165), respectively.

### *In vitro* deubiquitination assay

HEK 293 T-REx ^FLAG^XPO1 cells were grown in complete D-MEM in the presence of doxycycline to 80% confluence. Cells were then treated with 1 µM selinexor and 0.2 µM MG132 for additional 18 h before lysis. Cells were collected by centrifugation and lysed on ice for 30 min in NET-150 buffer. ^FLAG^XPO1 protein was recovered from whole lysates by immunoprecipitation using anti-FLAG M2 magnetic beads (Sigma Aldrich) and eluted by competition using the 3X FLAG peptide (APExBio) in TE buffer pH 8.0 (10 mM Tris, 1 mM EDTA). Subsequently, half of the elute was incubated with vOTU in DUB buffer (50 mM NaCl, 50 mM Tris, pH 7.5 and 50 mM DTT) for 30 min at RT. Immunoblotting with FLAG M2 antibody (Sigma Aldrich, F3165) was used to detect ^FLAG^XPO1 levels.

### siRNA knockdowns

ON-TARGETplus siRNA targeting the mRNAs for NAE1 (MU-006401-02) and CUL5 (MU-019553-01) was purchased from Dharmacon, Horizon Discovery, and MISSION esiRNA targeting the mRNAs for ELOB (EHU150621), ELOC (EHU903091), RBX1 (EHU055711) and RBX2 (EHU094071) were purchased from Sigma Aldrich. The negative control scrambled siRNA was Dharmacon, Horizon Discovery (D-001210-01). siRNAs were transfected into cells with Lipofectamine RNAiMAX (Thermo Fisher Scientific), according to the manufacturer’s instructions at 30 nM final concentration. The most efficient knockdown at protein levels was seen 48 h without compromising cell’s integrity. XPO1 degradation by KPT-185 treatment were performed at day 3 after knockdown. 18 h post-KPT-185 treatment, cells were lysed in RIPA buffer and immunoblotting was performed using primary antibodies: XPO1 (CRM1 rabbit polyclonal H-300; sc-5595, Santa Cruz or CRM1 mouse monoclonal (C-1): sc-74454), NAE1 (Cell Signaling Technology, #14321), ELOB (Proteintech, 12450-1-AP), ELOC (Bethyl Laboratories, A304-008A), CUL5 (Bethyl Laboratories, A302-173A), RBX1 (Cell Signaling Technology, #11922), RBX2 (Sigma Aldrich, HPA036995) and GAPDH (Loading Control Antibody Sampler Kit #5142, Cell Signaling Technology).

### Patient Sample Processing, Cell Lines, and Cell Culture

Blood was obtained from healthy volunteers or patients with CLL following written informed consent under a protocol approved by the Institutional Review Board of The Ohio State University (OSU; Columbus, OH) in accordance with the Declaration of Helsinki. All patients examined had CLL as defined by the 2008 International Workshop on Chronic Lymphocytic Leukemia criteria, and cells were isolated and cultured as previously described^24^. HG-3 (DSMZ, Germany), cells were cultured in RPMI 1640 (Gibco) supplemented with 10% fetal bovine serum (FBS, VWR) and 1% penicillin/streptomycin/L-glutamine (P/S/G; Gibco) at 37 °C and 5% CO_2_. Cell lines were validated via short tandem repeat analysis by The Ohio State University Genomic Services Core. For experiments cells were resuscitated, cultured in RPMI (Invitrogen) supplemented with 10% fetal bovine serum (FBS), and used within 3 to 4 weeks from thawing. All cell lines were confirmed to be *Mycoplasma* negative using the MycoAlert Mycoplasma Detection Kit from Lonza according to the manufacturer’s instructions and used within 3 to 4 weeks from thawing. Cells were lysed in RIPA buffer. Proteins were separated by SDS-PAGE and blots probed with commercially available antibodies and detected by addition of Enhanced chemiluminescence (ECL) substrate (Pierce) followed by quantification by Chemi-Doc system with Quantity One software (Bio-Rad Laboratories). XPO1 used SC-74454, 1:1000 dilution (experiments in **Figure 2f** and **Extended Data Figure 2g**) or 1:500 dilution (experiments in **Figure 6f** and **Extended Data Figure 18d**), full ECL. ASB8 used Invitrogen MA5-26984, 1:500 dilution, 50% ECL; GAPDH using SC-47724 ab, 1:1000 dilution, 50% ECL.

### Protein expression and purification

Full-length XPO1 was purified according to established protocol^82^. Human ASB8 cDNA in pENTR221 vector was purchased as an Ultimate ORF clone (Thermo Fisher Scientific, clone ID IOH3117), PCR amplified and inserted into pGEX-4T3 (Amp^R^) vector with a TEV protease site. Human ELOB (full-length) and ELOC (residues 17-112) were obtained in pACYC (Cam^R^) (gift from Dr. Elizabeth Komives). ASB8 plasmids were co-transformed into BL21-gold(DE3) *E. coli* competent cells containing pACYC-ELOB/C for co-expression of GST-ASB8•ELOB/C. Cultures were grown in LB media at 37 °C to an OD_600_ ∼0.7 and induced with 0.4 μM IPTG at 18 °C for 18 hours. Cells were pelleted by centrifugation at 4,500 x g for 30 minutes at 4 °C, resuspended in buffer (50 mM HEPES pH 7, 150 mM NaCl, 10% glycerol, 2 mM DTT, 5 mM EDTA, 5 mM EGTA) supplemented with 200 μg/mL Pefabloc, 10 μg/mL leupeptin, and 1 mM benzamidine protease inhibitors, and then lysed with the EmulsiFlex-C5 cell homogenizer (Avestin). GST-ASB8•ELOB/C was purified over Glutathione Sepharose 4B (GSH; GE Healthcare) resin followed by size exclusion chromatography (SEC) over a Superdex200 10/300 Increase column (Cytiva) pre-equilibrated in TB buffer (20 mM HEPES pH 7.4, 110 mM KOAc, 2 mM Mg(OAc)_2_, 10% glycerol, 2 mM DTT). ASB8^ΔN16^ and all ASB8 mutants were expressed and purified using the same protocol. GST-ASB8^ΔN16^•ELOB/C was cleaved from GSH-Sepharose resin by TEV protease for 1 h at room temperature prior to SEC. RAN^GTP^ was purified according to previous published method protocol^82^, his tag was not cleaved.

### *In vitro* binding assays

*In vitro* pull-down binding assays were performed by incubating 0.5 µM GST, GST-NES^MVM^ ^NS2^, or GST-ASB8 (WT or mutants) ± 1 µM XPO1 (WT or mutants) ± DMSO, 2 µM inhibitor (KPT-185, KPT-127, selinexor, eltanexor, felezonexor or LMB) or 5 μM inhibitor (KPT-UTSW1, –UTSW2 or –UTSW3) with 20 µL GSH-Sepharose beads (bead bed volume) (Cytiva) in a 200 µL reaction in TB Buffer for 30 min at 4°C. All inhibitors were pre-incubated with XPO1 for 30 minutes at 4°C with rotation, unless otherwise specified. For direct titration binding assays, 1 μM GST or GST-ASB8•ELOB/C ± 2.5 μM KPT-185 + XPO1 titrated from 1.25 – 10 μM ± RAN^GTP^ titrated from 3.75 – 30 μM was used. For XPO1 V430D validation pull-down assays, 0.5 μM GST or GST-NES + 0.5 μM XPO1 ± 1.5μM RAN^GTP^ was used. All samples were washed three times with 500 µL TB buffer, and bound proteins were separated and visualized by SDS-PAGE gel stained with Coomassie stain. All binding assays were repeated in triplicates. For quantification of direct titration binding assays, the ratio of XPO1/GST-ASB8 peak area was determined first and then normalized to the XPO1 WT + KPT-185 lane in each gel.

### Isothermal titration calorimetry

XPO1 and ASB8^ΔN16^•ELOB/C were dialyzed into ITC buffer (20 mM HEPES pH 7.5, 110 mM KOAc, 2 mM Mg(OAc)_2_, 10% v/v glycerol, 2 mM TCEP). ITC experiments were performed in a Micro-Cal PEAQ-ITC (Malvern Panalytical, Worcestershire, UK) calorimeter; it has a stirred 206.2 μL reaction cell held at 20°C (for reactions with selinexor) or 10°C (for reactions without selinexor). The first injections were 0.5 μL, followed by twenty 1.9 μL injections with a stirring rate of 750 rpm. XPO1 was used at 10 μM in the ITC cell; 70–190 μM ASB8^ΔN16^•ELOB/C were used in the syringe. For experiments with selinexor, 30 μM selinexor was added to XPO1 and ASB8^ΔN16^•ELOB/C after dialysis. All ITC experiments were performed in duplicate except when noted. ITC data were integrated and baseline corrected using NITPIC^83^. The integrated data were globally analyzed in SEDPHAT^84^ using a model considering a single class of binding sites. Thermogram and binding figures were plotted in GUSSI^85^.

### Purification of proteins for *in vitro* ubiquitination assay

Dr. Elizabeth Komives also gifted us the following plasmids for protein expression of the components used in the *in vitro* ubiquitination assay: pET28a-GB1-TEV-*^Hs^*CUL5, pET11a-*^Mm^*RBX2, pGex4T1-*^Hs^*NAE1/*^Hs^*UBA3, pET28a-NEDD8, pET28a-TEV-UBE2F, pET28a-TEV-UBE2L3 and pET11a-MBP-TEV-ARIH2. All plasmids were expressed individually in BL21-gold(DE3) *E. coli* competent cells except for pET28a-GB1-TEV-*^Hs^*CUL5 and pET11a-*^Mm^*RBX2 which were co-expressed at the same time. Cultures were grown in LB media at 37 °C with the appropriate antibiotic to an OD_600_ ∼0.7 and induced with 0.4 μM IPTG at 18 °C for 18 hours. Cells were pelleted by centrifugation at 4,500 x g for 30 minutes at 4 °C, and resuspended in lysis buffer supplemented with 200 μg/mL Pefabloc, 10 μg/mL leupeptin, and 1 mM benzamidine protease inhibitors and frozen in – 80°C. Lysis buffer for His-CUL5/RBX2, His-UBE2L3 and His-MBP-ARIH2 contained 50 mM HEPES pH 7.5, 150 mM NaCl, 10 mM imidazole pH 8.0, 2 mM β-mercaptoethanol, 10% glycerol. His-NEDD8 and His-UBE2F used the same buffer but with Tris pH 8.0 instead of HEPES pH 7.5. For GST-NAE1/UBA3, lysis buffer contained 50 mM HEPES pH 7.0, 150 mM NaCl, 2 mM DTT, 10% glycerol + protease inhibitors. All lysates were thawed and lysed with the EmulsiFlex-C5 cell homogenizer (Avestin) and clarified with 48,400 RCF at 4°C for 40 min.

Supernatant of His-CUL5/RBX2 and His-UBE2L3 were individually incubated with Ni NTA beads (Qiagen) for 30 mins at 4 °C, and the beads were washed with buffer containing 50 mM HEPES 7.5, 150 mM NaCl, 25 mM imidazole pH 8.0, 2 mM β-mercaptoethanol, 10% glycerol, before elution with 50 mM HEPES 7.5, 150 mM NaCl, 250 mM imidazole pH 8.0, 2 mM β-mercaptoethanol, 10% glycerol. The eluted proteins were concentrated to ∼15 mL and incubated with 1 mg of TEV protease for 1 at room temp to remove the His tags. The reaction is diluted and put through Ni-NTA beads to remove the TEV, his tags and uncut proteins. For CUL5/RBX the TEV reaction was diluted with lysis buffer described above, and flowthrough containing CUL5/RBX2 is further concentrated and purified by SEC in SEC buffer containing 25mM HEPES pH 7.5, 150mM NaCl, 2.5mM MgCl, 10% glycerol, 1mM DTT. The fractions containing the least degradation species of CUL5 were concentrated, flash frozen and stored at –80 °C. For UBE2L3, the TEV reaction was diluted with buffer containing 25 mM HEPES 7.0, 25 mM NaCl, 2 mM DTT, 10% glycerol and the flowthrough containing UBE2L3 was further purified using HiTrap SP (Cytiva) using a salt gradient of 25 mM to 1M in the buffer. A single peak containing clean UBE2L3 was concentrated, flash frozen and stored at –80°C.

Supernatant of His-NEDD8 and His-UBE2F were also similarly subjected to Ni-NTA purification as described above for His-CUL5/RBX2 and His-UBE2L3 but Tris pH 8.0 was used in the buffers. Elution from Ni-NTA column containing His-NEDD8 was concentrated and diluted with buffer containing 25 mM Tris pH 8.0, 25 mM NaCl, 2 mM DTT, 10% glycerol and purified using HiTrap Q (Cytiva). Elution from Ni-NTA column containing His-UBE2F was concentrated to ∼15 mL and incubated with 1 mg of TEV protease for 1 h at room temp. The reaction is diluted with buffer containing 25 mM Tris pH 8.0, 25 mM NaCl, 2 mM DTT, 10% glycerol and put through Ni-NTA beads. The flowthrough containing UBE2F was further purified using HiTrap Q. For both His-NEDD8 and UBE2F, most of the proteins remained in the flowthrough while other contaminants bound to the Q column, therefore the flowthroughs were concentrated, flash frozen and stored at –80 °C.

Supernatant containing GST-NAE1/UBA3 was incubated with GSH Sepharose beads (Cytiva) for 30 mins at 4 °C, washed with the lysis buffer and eluted with lysis buffer supplemented with 30 mM GSH. The elution was concentrated and further purified in a HiLoad 16/600 200pg (Cytiva) column using SEC buffer mentioned above. The peak containing the cleanest GST-NAE1/UBA3 was concentrated, flash frozen and stored at – 80 °C. Supernatant containing His-MBP-ARIH2 was passthrough Amylose beads (New England Biolabs) for binding. Amylose beads were then washed with buffer containing 20 mM HEPES 7.5, 150 mM NaCl, 10% glycerol, 1 mM DTT and His-MBP-ARIH2 was eluted with the same buffer supplemented with 20 mM maltose. The elution was concentrated to ∼15 mL and incubated with 1 mg of TEV protease for 1 h at room temp to cleave the His-MBP tag. The reaction was put through Ni-NTA beads and washed with lysis buffer. The flowthrough containing ARIH2 is then concentrated to be purified in in a HiLoad 16/600 200pg column using SEC buffer mentioned above. The peak corresponding to ARIH2 was concentrated, flash frozen and stored at –80 °C.

To generate neddylated-CUL5-RBX2, ∼2 mg of GST-NAE1/UBA3 and ∼0.2 mg His-NEDD8 was incubated (0.8:1 molar ratio) in SEC buffer supplemented with 10 mM ATP-Mg(OAc)_2_ in room temp for 15 min in a 1 mL reaction. Then CUL5-RBX2 and UBE2F are added in a 1:1:1 ratio to His-NEDD8 in a 2 mL reaction with additional 10 mM ATP-Mg(OAc)_2_ for another 30 mins at room temp. The reaction was supplemented with 10 mM imidazole pH 7.8 and bound to Ni-NTA beads to capture His-NEDD8∼CUL5/RBX2. The beads were washed and eluted with the same buffers used for CUL5/RBX2 His-tag purification. The elution was concentrated to 1 mL and injected to a Superdex 200 Increase 10/300 GL column for a final purification step in the SEC buffer.

### *In vitro* ubiquitination assay

For the poly-ubiquitination assay, 10 µM XPO1 was first incubated with 20 µM RAN^GTP^ and compounds or DMSO for 15 mins at room temp in buffer containing 25 mM HEPES pH 7.5, 150 mM NaCl, 10% glycerol. Reactions containing 20 µM ubiquitin (Sigma-Aldrich Cat# U5382-1MG, stock solution made with protease inhibitors), 0.2 µM E1 (Ubiquitylation Assay Kit, Abcam Cat# ab139467), 5 mM ATP-MgCl, 1 µM UBE2L3, 1.5 µM ARIH2, 1 µM neddylated CUL5-RBX2, 1 µM ASB8 FL-ELOB/C and 2 µM pre-incubated XPO1-RAN^GTP^-selinexor complexes were assembled in 10 µL in the aforementioned buffer and incubated for 30 min at room temp. Then 2.5 µM UBE2R2 (Active Motif Cat# 82002), 2 µM UBCH3 (Ubiquitylation Assay Kit, Abcam Cat# ab139467) or 2 µM UBCH5a (Sigma-Aldrich Cat# 23-029-M) was added and reaction was incubated for another 2 h at room temp. Reaction was stopped with SDS-PAGE sample buffer and boiled. Sample was diluted 40X and 1 µL sample was loaded for SDS-PAGE and transfer. Membrane was cut into two pieces at 150 kDa to ensure efficient visualization of the higher molecular weight species. Two blots were imaged separately by XPO1 antibody (Novus, NB100-79802, 1:2000 dilution).

For other assays shown in Extended Data, reactions contained 2.5 µM ubiquitin, 0.1 µM E1, 5 mM ATP-Mg (all three from Ubiquitylation Assay Kit, Abcam Cat# ab139467), 1 µM UBE2L3, 2 µM ARIH2, 1 µM neddylated CUL5-RBX2, 1 µM ASB8 FL-ELOB/C and 1 µM pre-incubated XPO1-inhibitor or DMSO complexes. Reactions were incubated for 2 h at room temp and stopped with SDS-PAGE sample buffer and boiled. Sample was diluted 40X and 1 µL sample was loaded for western blot visualization with XPO1 antibody (Novus, NB100-79802, 1:2000 dilution).

### CryoEM sample preparation and data collection

For KPT-185/KPT-127/KPT-UTSW1-XPO1•ASB8^ΔN16^•ELOB/C and selinexor-XPO1•ASB8^ΔN16/R197A^•ELOB/C, 2mg of XPO1 was mixed with GST-ASB8^ΔN16^•ELOB/C or GST-ASB8^ΔN16/R197A^•ELOB/C and the indicated inhibitors (1:2:2 molar ratio) and 1 mg of TEV protease in TB buffer (20 mM HEPES pH 7.4, 110 mM KOAc, 2 mM Mg(OAC)_2_, 2 mM DTT, 10% glycerol) for 1 hour at room temperature, then passed over GSH-Sepharose resin. Flowthrough was concentrated to 1 mL. For RAN^GTP^•selinexor-XPO1•ASB8^ΔN16^•ELOB/C, 2mg of XPO1 was mixed with ASB8^ΔN16^-ELOB/C with no GST tag, RAN^GTP^ and KPT-185 (1:2:2:2 molar ratio). All protein samples were purified over a Superdex200 10/300 Increase column (Cytiva) pre-equilibrated in cryoEM buffer (20 mM HEPES pH 7.4, 150 mM NaCl, 2 mM Mg(OAc)_2_, 2 mM TCEP). 4 µL of complex sample (0.4 mg/ml of KPT-185-XPO1•ASB8^ΔN16^•ELOB/C supplemented with 2-fold excess KPT-185, or 0.5 mg/ml of RAN^GTP^•selinexor-XPO1•ASB8^ΔN16^•ELOB/C supplemented with 2-fold excess ASB8^ΔN16^-ELOB/C and selinexor) was applied to a 400-mesh copper grid (Quantifoil R1.2/1.3). For KPT-127/KPT-UTSW1-XPO1•ASB8^ΔN16^•ELOB/C or selinexor-XPO1•ASB8^ΔN16/R197A^•ELOB/C, 4 μL of 0.8 mg/ml complex sample supplemented with 2-fold excess inhibitor was applied to a 300-mesh copper grid (C-Flat 1.2/1.3). For unliganded XPO1, XPO1 was buffer exchanged into cryoEM buffer and diluted to 3.4 mg/mL and supplemented with 0.025% Tyloxapol (BioVision). For selinexor-XPO1, XPO1 was diluted to 3 mg/mL with two times molar excess of Selinexor and supplemented with 0.025% Tween-20 (BioVision). 4 µL of each sample were applied on Quantifoil R1.2/1.3, 300 mesh copper grids. All grids were pre-glow-discharged using a PELCO easiGlow glow discharge apparatus (Ted Pella) at 30 mA for 80 s. Excess sample was blotted and plunge frozen in liquid ethane using the Vitrobot Mark IV System (Thermo Fisher) at 4 °C with 95% humidity.

One RAN^GTP^•selinexor-XPO1•ASB8^ΔN16^•ELOB/C grid was used for 24 h data collection at the UT Southwestern Cryo-Electron Microscopy Facility (CEMF) on a Titan Krios microscope (Thermo Fisher) at 300 kV with a 10 eV slit-width Bioquantum energy filter and a Gatan K3 detector in non-correlated double sampling (CDS) super-resolution mode using SerialEM software^86^. One KPT-185-XPO1•ASB8^ΔN16^•ELOB/C grid and one selinexor-XPO1 grid were used for 18 and 24 h data collection, respectively, at the CEMF with CDS-mode. One unliganded XPO1 grid and one selinexor-XPO1•ASB8^ΔN16/R197A^•ELOB/C grid were used for 24 h data collection on a Titan Krios 300kV microscope equipped with a Falcon 4 detector and a 10 eV slit-width Selectris X energy filter at the CEMF. One KPT-127-XPO1•ASB8^ΔN16^•ELOB/C grid was used for 24 h data collection on a Titan Krios 300kV microscope equipped with a Falcon 4i detector and a 10 eV slit-width Selectris X energy filter at the Pacific Northwest Center for CryoEM (PNCC). Two KPT-UTSW1-XPO1•ASB8^ΔN16^•ELOB/C grids were used for 48 h data collection on a Titan Krios 300kV microscope equipped with a Gatan K3 detector and a 10 eV slit-width Bioquantum energy filter at the CEMF. More details are listed in **Extended Data Tables 1 and 3**.

### CryoEM data processing and structure determination

All data processing was performed using the software cryoSPARC v4.2, 4.4 or 4.6^87^. Movies were imported, subjected to patch motion correction and patch CTF estimation. For RAN^GTP^•selinexor-XPO1•ASB8^ΔN16^•ELOB/C, KPT-185-XPO1•ASB8^ΔN16^•ELOB/C and selinexor-XPO1 dataset, a ½ F-crop factor was applied during patch motion correction.

For KPT-185-XPO1•ASB8^ΔN16^•ELOB/C (**Extended Data Figure 6a**), initial blob picking was performed on 795 micrographs and followed by two rounds of 2D classification to obtain templates used for template picking on the full dataset, which resulted in 2,386,555 particles. After two rounds of 2D classification, 627,555 particles were split into three classes for ab-initio reconstruction and further classified using heterogeneous refinement. One class had a 4.96 Å map showing helical features resembling XPO1 and ASB8 (193,789 particles). To obtain more particles, these particles were used to train Topaz^88^, which picked 1,066,266 particles. After rounds of 2D classification, the remaining 770,623 particles were used for ab-initio reconstruction and heterogeneous refinement, seeking four classes. One class (205,462 particles) resembled the initial map used to train Topaz and was subsequently refined to 3.37 Å using Non-Uniform (NU) refinement with optimization of the per-exposure-group CTF parameters (defaults and spherical aberration).

For RAN^GTP^•selinexor-XPO1•ASB8^ΔN16^•ELOB/C (**Extended Data Figure 5a**), template was generated using the initial model and used for template picking on full dataset, which resulted in 1,424,491 particles. After 1 round of 2D classification and 3 rounds of 3D heterogenous refinement, particle stack of 171,837 particles were used for Topaz training, which yielded 1,821,140 particles. After 1 round of 2D classification and 6 additional rounds of 3D heterogenous refinement (with 2 classes to clean up junk particles), 78,241 particles were used in NU after global CTF refinement with default parameters for 3 iterations, which yielded the best resolution map at 3.25 Å.

For unliganded XPO1 micrographs (**Extended Data Figure 7a**), blob picker was used to select 3,643,569 initial particles. After 4 rounds of 2D classification, 851,026 particles were used to obtain 3 *ab initio* models, which were subsequently submitted to heterogeneous refinement. ∼36% of the particles partitioned into a map that looks like intact XPO1, which were used to train Topaz for picking 1,245,354 initial particles. After 1 round of 2D classification, particles were classified into the previous 3 classes by heterogeneous refinement. 375,108 particles (50% out of the final 753,568 particles) were intact and subjected to final NU refinement with optimizations of per-particles defocus and per-group CTF parameters (Tilt, Trefoil, Spherical Aberration and Anisotropic Magnification) to yield final map of 2.93 Å resolution.

For selinexor-XPO1 micrographs (**Extended Data Figure 8a**), templates were generated using unliganded XPO1 particles and template picker was used to select 4,541,251 initial particles. After 7 more rounds of 2D classification, 688,481 particles were used to obtain 4 *ab initio* models, which were submitted to heterogeneous refinement.

∼42% of the particles partitioned into a map that looks like intact XPO1. The good particles were used to train Topaz for picking, which extracted 1,197,263 initial particles. After 3 rounds of 2D classification, particles were classified into the previous 4 classes by heterogeneous refinement. 431,664 particles (52% out of the final 829,704 particles) were subjected to local CTF refinement and then a final NU refinement with optimizations of the per-particle defocus and per-group CTF parameters (Tilt and Trefoil only) to yield final map of 3.37 Å resolution.

For selinexor-XPO1•ASB8^ΔN16/R197A^•ELOB/C (**Extended Data Figure 21b**), initial blob picking was performed on 2,792 micrographs and followed by two rounds of 2D classification to obtain templates used for template picking on the full dataset, which resulted in 900,655 particles. After one round of 2D classification, 300,890 particles were split into two classes for ab-initio reconstruction and further classified using heterogeneous refinement. One class had a 5.67Å map showing helical features resembling XPO1 and ASB8 (300,890 particles). This map was used to create templates for template picking on the full dataset (6,975 micrographs). After thirteen rounds of heterogenous refinement, the remaining 433,178 particles were subjected to NU refinement, global CTF refinement with optimization of spherical aberration. A final NU refinement yielded a final map of 2.49 Å.

For KPT-127-XPO1•ASB8^ΔN16^•ELOB/C (**Extended Data Figure 19a**), blob picking yielded 3,769,085 particles. After two rounds of 2D classification, 1,701,241 particles were further classified using two rounds of 3D heterogenous refinement of two classes, with the selinexor-XPO1•ASB8^ΔN16/R197A^•ELOB/C final map as a starting volume for one class, and a decoy class generated by killing an ab-initio reconstruction job. The desired class contained 498,390 particles, which were used to train Topaz for particle picking, yielding 1,092,211 particles. After one round of 2D classification, the 969,083 remaining particles were further classified by eight rounds of heterogenous refinement using the same starting map and decoy map. The final set of 118,442 particles were subjected to NU refinement followed by three rounds of global CTF refinement with optimization of spherical aberration and EWS curvature. A final round of NU refinement yielded a map of 3.28 Å.

For the first dataset of KPT-UTSW1-XPO1•ASB8^ΔN16^•ELOB/C (**Extended Data Figure 20a**), the selinexor-XPO1•ASB8^ΔN16/R197A^•ELOB/C final map was used to generate templates for template picking, yielding 3,314,896 particles. 2,875,712 particles remained after two rounds of 2D classification. The particles were split into three classes by heterogenous refinement, using the template map and two decoy maps as starting volumes and resulting in 875,959 particles. The same templates were used for the second dataset of KPT-UTSW1-XPO1•ASB8^ΔN16^•ELOB/C, yielding 4,056,850 particles. One round of 2D classification gave 3,697,926 particles, which were split into four classes by heterogenous refinement: one class with XPO1 bound to ASB8•ELOB/C using the selinexor-XPO1• ASB8^ΔN16/R197A^•ELOB/C final map, unliganded XPO1 final map, and two decoy maps as starting volumes. The XPO1•ASB8•ELOB/C class containing 830,258 particles was further refined through six rounds of heterogenous refinement with one XPO1•ASB8•ELOB/C class and one decoy class. One round of 2D classification resulted in 250,354 particles used to train Topaz for particle picking, which yielded 1,205,849 particles. The particles from the two datasets were combined and subjected to nine rounds of heterogenous refinement using the template volume and a junk volume to start. The final 215,025 particles were used for NU refinement, followed by three rounds of global CTF refinement and a final round of NU refinement, resulting in a final map of 4.21Å.

### CryoEM model building, refinement and analysis

All initial models (specified in **Extended Data Tables 1 and 3**) were roughly docked into unsharpened maps using UCSF Chimera or ChimeraX^89,90^ and then subjected to real-space refinement with global minimization and rigid body restraints in Phenix^91^ and manual model building using ISOLDE^92^ and Coot^93^. Model validation was performed in Phenix. For KPT-185-XPO1•ASB8^ΔN16^•ELOB/C, deepEMhancer^94^ was used to generate maps to assist in modeling of N– and C-terminal HEAT repeats of XPO1. For selinexor-XPO1, KPT-185-XPO1•ASB8^ΔN16^•ELOB/C and RAN^GTP^•selinexor-XPO1•ASB8^ΔN16^•ELOB/C, the Cys528-selinexor/KPT-185 conjugation was restrained to be 1.8 Å long during real-space refinement in Phenix to aid placement of the inhibitor and the cysteine sidechain. For selinexor-XPO1, the location of selinexor was optimized by manual building in the final step as the density for the ligand does not cover the whole inhibitor. Structures were analyzed using PyMOL 2.4 and 2.5^95^, PDBe PISA for the binding interface^96^. and the APBS electrostatic plugin for 3D structure and electrostatic analysis^95,97^. All figures were generated using PyMOL or ChimeraX.

### **X-** ray structure determination

Protein purification, complex formation, crystallization of KPT-185-*^Sc^*XPO1^EH^•RAN^GTP^•RANBP1 and KPT-396-*^Sc^*XPO1•RAN^GTP^•RANBP1 were all performed according to published protocol^98^. KPT-185 was used in 2 times molar excess while KPT-396 was used at 5 times excess. Both complexes were concentrated to ∼10 mg/mL. Crystals were formed overnight using hanging-drop method in crystallization condition of 17% PEG3350, 100 mM Bis-Tris (pH 6.6) and 200 mM ammonium nitrate and cryoprotected with 23% PEG3350 and 12% (v/v) glycerol for freezing. X-ray diffraction data was collected at Advanced Photon Source – Structural Biology Center 19ID beamline and processed using HKL3000^80^. Structure models were generated by refining 4GMX as starting model against the collected data using PHENIX^91,99,100^. X-ray/stereochemistry and X-ray/ADP weight were optimized by phenix.refine in later steps. Manual building was performed in Coot^101^.

### *In vivo* XPO1 degradation assay with Simple Western

To establish HAP1 knockout cells, two sgRNAs per gene from the Brunello knockout library were selected^102^, cloned into plentiGuide-Puro (Addgene #59263)^103^ and individually packaged into lentivector^7^. Next, HAP1-Cas9 cells were stably transduced by spinfection followed by selection with 1 µg/mL puromycin until the no-lentivector control was completely dead. For overexpression, HAP1 cells were stably transduced with a lentivector expressing the ASB8 coding sequence. HAP1 cells carrying an endogenous XPO1 C528S mutation were established using CRISPR-mediated homology-directed repair^7,29^. To induce XPO1 degradation, cells were treated with selinexor, SINE analogs or 4-octyl itaconate. Cells were treated with compound overnight unless indicated otherwise.

HEK293T cells (ATCC) were transiently (co-)transfected with pDNA expressing XPO1 or ASB8 (WT or mutant) using Turbofectin 8.0 (OriGene) according to the manufacturer’s protocol. After 24 h, media were replaced with fresh media containing selinexor as indicated and cells were incubated for another 24 h. HAP1 and HEK293T cells were subsequently collected and lysed in RIPA buffer (Thermo Fisher Scientific) containing 1× HALT Protease Inhibitor Cocktail (Thermo Fisher Scientific). We used a Simple Western size assay (JESS, Bio-Techne) for immunodetection of XPO1 (NB100-79802, Novus, 1:1000), FLAG (MAB8529, Bio-Techne, 1:50) and β-tubulin (NB600-936, Novus, 1:40). Data analysis was done with Compass for Simple Western software (Bio-Techne). We used RePlex and Total Protein Detection modules to correct XPO1 peak area for variations in sample loading. Alternatively, we corrected by dividing by β-tubulin peak area.

### *In vivo* ubiquitination

HEK293T cells were co-transfected as indicated with pDNA expressing FLAG-XPO1, Tag100-ASB8 (WT or mutant) and c-Myc-UBC (WT or K7R) using Turbofectin 8.0 (OriGene) according to the manufacturer’s protocol. After 24 h, media were replaced with fresh media containing selinexor and bortezomib as indicated. After 6 h of incubation, cells were collected and lysed in RIPA buffer (Thermo Fisher Scientific) containing 1× HALT Protease Inhibitor Cocktail (Thermo Fisher Scientific). Whole-cell lysates were first pre-cleared by incubation with Dynabeads Protein A (Thermo Fisher Scientific) on a rotary wheel at 4 °C for 4 h. Next, the non-bound fraction was incubated with either anti-FLAG M2 or anti-c-Myc magnetic beads (Sigma-Aldrich) at 4°°C overnight. Bound proteins were eluted by boiling in 1× Sample Buffer (Bio-Techne) with 1% SDS at 95 °C for 10 min. We used Simple Western detection of XPO1 (IP with anti-FLAG: NB100-79802, Novus Biologicals, 1:1000; IP with anti-c-Myc: sc-74454, Santa Cruz Biotechnology, 1:50), c-Myc (NB600-336, Novus Biologicals, 1:50), ASB8 (NBP1-83620, Novus Biologicals, 1:50) or total protein (using RePlex and Total Protein Detection with chemiluminescence).

### Cell viability assay

HAP1 cells (Horizon Discovery) were transduced at low multiplicity of infection (MOI<0.1) with a lentiviral vector for stable overexpression of either wild type or mutant ASB8. Cells were treated with a serial dilution of selinexor for a period of three days. Next, 20 µL MTS reagent (CellTiter 96 Aqueous Non-Radioactive Cell Proliferation Assay from Promega) was added and cells were incubated for an additional 2 h after which absorbance was measured at 490 nm with a Safire 2 microplate reader (Tecan). For each drug concentration, the absorbance values of three replicate wells were averaged, blank corrected and normalized to those of untreated cells. Dose-response curves were obtained by fitting a non-linear, four-parameter regression model to relative cell viability values using GraphPad Prism 10.

### Synthesis of smaller triazole acrylate inhibitors

The full methods for synthesis^104^ and characterization of KPT-UTSW1, KPT-UTSW2 (KPT-9661) and KPT-UTSW3 are described in Supplemental Methods.

### Cellular cargo inhibition assay

HeLa Kyoto cells stably expressing the XPO1-dependent NLSSV40-AcGFP-NESPKI reporter^105^ were seeded at 8,000 cells per well in 96-well tissue culture plates. The following day, cells were treated with serial dilutions of selinexor or small SINE analogs for 3 hours, fixed, and counterstained with DAPI. Fluorescence in the blue and green channels was imaged using an ArrayScan XTI High Content Reader (Thermo Fisher Scientific). In each cell, nuclear and cytoplasmic compartments were segmented and their average pixel intensities in the green channel were quantified using HCS Studio software (Thermo Fisher Scientific). A ratio of nuclear to cytoplasmic signal equal to or above 1.25 was considered predominantly nuclear. GraphPad Prism was used for dose-response curve fitting based on the percentage of cells with a predominantly nuclear localization of the reporter cargo. Representative images were acquired using a Leica TCS SP5 confocal microscope (Leica Microsystems) with an HCX PL APO 63× (NA 1.2) water immersion objective.

### Data availability statement

Coordinates and cryoEM density maps have been deposited in the Protein Data Bank (PDB) under accession codes 9OG9, 9OGA, 9OGB, 9OGC, 9OGD, 9OGE and 9OGF, and the Electron Microscopy Data Bank (EMDB) under accession codes EMD-70458, EMD-70459, EMD-70460, EMD-70461, EMD-70462, EMD-70463 and EMD-70464, respectively. Coordinates and structure factors have been deposited in the PDB under accession codes 9OGN and 9OGO. All data are available upon request.

## References

1 Wing, C. E., Fung, H. Y. J. & Chook, Y. M. Karyopherin-mediated nucleocytoplasmic transport. Nat Rev Mol Cell Biol 23, 307–328 (2022). 10.1038/s41580-021-00446-7

2 Azmi, A. S., Uddin, M. H. & Mohammad, R. M. The nuclear export protein XPO1 – from biology to targeted therapy. Nat Rev Clin Oncol 18, 152–169 (2021). 10.1038/s41571-020-00442-4

3 De Cesare, M. et al. Anti-tumor activity of selective inhibitors of XPO1/CRM1-mediated nuclear export in diffuse malignant peritoneal mesothelioma: the role of survivin. Oncotarget 6, 13119–13132 (2015). 10.18632/oncotarget.3761

4 Tai, Y. T. et al. CRM1 inhibition induces tumor cell cytotoxicity and impairs osteoclastogenesis in multiple myeloma: molecular mechanisms and therapeutic implications. Leukemia 28, 155–165 (2014). 10.1038/leu.2013.115

5 Ranganathan, P. et al. Preclinical activity of a novel CRM1 inhibitor in acute myeloid leukemia. Blood 120, 1765–1773 (2012). 10.1182/blood-2012-04-423160

6 Zheng, Y. et al. KPT-330 inhibitor of XPO1-mediated nuclear export has anti-proliferative activity in hepatocellular carcinoma. Cancer Chemother Pharmacol 74, 487–495 (2014). 10.1007/s00280-014-2495-8

7 Kwanten, B. et al. E3 ubiquitin ligase ASB8 promotes selinexor-induced proteasomal degradation of XPO1. Biomed Pharmacother 160, 114305 (2023). 10.1016/j.biopha.2023.114305

8 Bondeson, D. P. & Crews, C. M. Targeted Protein Degradation by Small Molecules. Annu Rev Pharmacol Toxicol 57, 107–123 (2017). 10.1146/annurev-pharmtox-010715-103507

9 Chamberlain, P. P. et al. Structure of the human Cereblon-DDB1-lenalidomide complex reveals basis for responsiveness to thalidomide analogs. Nat Struct Mol Biol 21, 803–809 (2014). 10.1038/nsmb.2874

10 Bussiere, D. E. et al. Structural basis of indisulam-mediated RBM39 recruitment to DCAF15 E3 ligase complex. Nat Chem Biol 16, 15–23 (2020). 10.1038/s41589-019-0411-6

11 Gorlich, D. & Mattaj, I. W. Nucleocytoplasmic transport. Science 271, 1513–1518 (1996). 10.1126/science.271.5255.1513

12 Komiyama, K. et al. Antitumor activity of leptomycin B. J Antibiot (Tokyo) 38, 427–429 (1985).

13 Newlands, E. S., Rustin, G. J. & Brampton, M. H. Phase I trial of elactocin. Br J Cancer 74, 648–649 (1996). 10.1038/bjc.1996.415

14 Parikh, K., Cang, S., Sekhri, A. & Liu, D. Selective inhibitors of nuclear export (SINE)--a novel class of anti-cancer agents. J Hematol Oncol 7, 78 (2014). 10.1186/s13045-014-0078-0

15 Van Neck, T. et al. Inhibition of the CRM1-mediated nucleocytoplasmic transport by N-azolylacrylates: structure-activity relationship and mechanism of action. Bioorg Med Chem 16, 9487–9497 (2008). 10.1016/j.bmc.2008.09.051

16 Daelemans, D. et al. A synthetic HIV-1 Rev inhibitor interfering with the CRM1-mediated nuclear export. Proc Natl Acad Sci U S A 99, 14440–14445 (2002). 10.1073/pnas.212285299

17 Attiyeh, E. F. et al. Pharmacodynamic and genomic markers associated with response to the XPO1/CRM1 inhibitor selinexor (KPT-330): A report from the pediatric preclinical testing program. Pediatr Blood Cancer 63, 276–286 (2016). 10.1002/pbc.25727

18 Etchin, J. et al. Activity of a selective inhibitor of nuclear export, selinexor (KPT-330), against AML-initiating cells engrafted into immunosuppressed NSG mice. Leukemia 30, 190–199 (2016). 10.1038/leu.2015.194

19 Yoshimura, M. et al. Induction of p53-mediated transcription and apoptosis by exportin-1 (XPO1) inhibition in mantle cell lymphoma. Cancer Sci 105, 795–801 (2014). 10.1111/cas.12430

20 Hing, Z. A. et al. Next-generation XPO1 inhibitor shows improved efficacy and in vivo tolerability in hematological malignancies. Leukemia 30, 2364–2372 (2016). 10.1038/leu.2016.136

21 Syed, Y. Y. Selinexor: First Global Approval. Drugs 79, 1485–1494 (2019). 10.1007/s40265-019-01188-9

22 Vercruysse, T. et al. The Second-Generation Exportin-1 Inhibitor KPT-8602 Demonstrates Potent Activity against Acute Lymphoblastic Leukemia. Clin Cancer Res 23, 2528–2541 (2017). 10.1158/1078-0432.CCR-16-1580

23 Abdul Razak, A. R. et al. First-in-Class, First-in-Human Phase I Study of Selinexor, a Selective Inhibitor of Nuclear Export, in Patients With Advanced Solid Tumors. J Clin Oncol 34, 4142–4150 (2016). 10.1200/JCO.2015.65.3949

24 Lapalombella, R. et al. Selective inhibitors of nuclear export show that CRM1/XPO1 is a target in chronic lymphocytic leukemia. Blood 120, 4621–4634 (2012). 10.1182/blood-2012-05-429506

25 Sun, Q. et al. Nuclear export inhibition through covalent conjugation and hydrolysis of Leptomycin B by CRM1. Proc Natl Acad Sci U S A 110, 1303–1308 (2013). 10.1073/pnas.1217203110

26 Etchin, J. et al. Antileukemic activity of nuclear export inhibitors that spare normal hematopoietic cells. Leukemia 27, 66–74 (2013). 10.1038/leu.2012.219

27 Haines, J. D. et al. Nuclear export inhibitors avert progression in preclinical models of inflammatory demyelination. Nat Neurosci 18, 511–520 (2015). 10.1038/nn.3953

28 Walker, J. S. et al. Recurrent XPO1 mutations alter pathogenesis of chronic lymphocytic leukemia. J Hematol Oncol 14, 17 (2021). 10.1186/s13045-021-01032-2

29 Neggers, J. E. et al. Identifying drug-target selectivity of small-molecule CRM1/XPO1 inhibitors by CRISPR/Cas9 genome editing. Chem Biol 22, 107–116 (2015). 10.1016/j.chembiol.2014.11.015

30 Sakakibara, K. et al. CBS9106 is a novel reversible oral CRM1 inhibitor with CRM1 degrading activity. Blood 118, 3922–3931 (2011). 10.1182/blood-2011-01-333138

31 Saito, N. et al. CBS9106-induced CRM1 degradation is mediated by cullin ring ligase activity and the neddylation pathway. Mol Cancer Ther 13, 3013–3023 (2014). 10.1158/1535-7163.MCT-14-0064

32 Mendonca, J. et al. Selective inhibitors of nuclear export (SINE) as novel therapeutics for prostate cancer. Oncotarget 5, 6102–6112 (2014). 10.18632/oncotarget.2174

33 Azmi, A. S. et al. Targeting the Nuclear Export Protein XPO1/CRM1 Reverses Epithelial to Mesenchymal Transition. Sci Rep 5, 16077 (2015). 10.1038/srep16077

34 Kudo, N. et al. Leptomycin B inactivates CRM1/exportin 1 by covalent modification at a cysteine residue in the central conserved region. Proc Natl Acad Sci U S A 96, 9112–9117 (1999). 10.1073/pnas.96.16.9112

35 Wang, S., Han, X., Wang, J., Yao, J. & Shi, Y. Antitumor effects of a novel chromosome region maintenance 1 (CRM1) inhibitor on non-small cell lung cancer cells in vitro and in mouse tumor xenografts. PLoS One 9, e89848 (2014). 10.1371/journal.pone.0089848

36 Liu, Y. et al. Molecular cloning and characterization of the human ASB-8 gene encoding a novel member of ankyrin repeat and SOCS box containing protein family. Biochem Biophys Res Commun 300, 972–979 (2003). 10.1016/s0006-291x(02)02971-6

37 Lumpkin, R. J., Baker, R. W., Leschziner, A. E. & Komives, E. A. Structure and dynamics of the ASB9 CUL-RING E3 Ligase. Nat Commun 11, 2866 (2020). 10.1038/s41467-020-16499-9

38 Kohroki, J., Nishiyama, T., Nakamura, T. & Masuho, Y. ASB proteins interact with Cullin5 and Rbx2 to form E3 ubiquitin ligase complexes. FEBS Lett 579, 6796–6802 (2005). 10.1016/j.febslet.2005.11.016

39 Mahrour, N. et al. Characterization of Cullin-box sequences that direct recruitment of Cul2-Rbx1 and Cul5-Rbx2 modules to Elongin BC-based ubiquitin ligases. J Biol Chem 283, 8005–8013 (2008). 10.1074/jbc.M706987200

40 Huttenhain, R. et al. ARIH2 Is a Vif-Dependent Regulator of CUL5-Mediated APOBEC3G Degradation in HIV Infection. Cell Host Microbe 26, 86–99 e87 (2019). 10.1016/j.chom.2019.05.008

41 Lumpkin, R. J., Ahmad, A. S., Blake, R., Condon, C. J. & Komives, E. A. The Mechanism of NEDD8 Activation of CUL5 Ubiquitin E3 Ligases. Mol Cell Proteomics 20, 100019 (2021). 10.1074/mcp.RA120.002414

42 Kostrhon, S. et al. CUL5-ARIH2 E3-E3 ubiquitin ligase structure reveals cullin-specific NEDD8 activation. Nat Chem Biol 17, 1075–1083 (2021). 10.1038/s41589-021-00858-8

43 Saito, N. & Matsuura, Y. A 2.1-A-resolution crystal structure of unliganded CRM1 reveals the mechanism of autoinhibition. J Mol Biol 425, 350–364 (2013). 10.1016/j.jmb.2012.11.014

44 Monecke, T. et al. Structural basis for cooperativity of CRM1 export complex formation. Proc Natl Acad Sci U S A 110, 960–965 (2013). 10.1073/pnas.1215214110

45 Shaikhqasem, A., Dickmanns, A., Neumann, P. & Ficner, R. Characterization of Inhibition Reveals Distinctive Properties for Human and Saccharomyces cerevisiae CRM1. J Med Chem 63, 7545–7558 (2020). 10.1021/acs.jmedchem.0c00143

46 Dian, C. et al. Structure of a truncation mutant of the nuclear export factor CRM1 provides insights into the auto-inhibitory role of its C-terminal helix. Structure 21, 1338–1349 (2013). 10.1016/j.str.2013.06.003

47 Koyama, M. & Matsuura, Y. An allosteric mechanism to displace nuclear export cargo from CRM1 and RanGTP by RanBP1. EMBO J 29, 2002–2013 (2010). 10.1038/emboj.2010.89

48 Kim, Y. K. et al. Structural basis of intersubunit recognition in elongin BC-cullin 5-SOCS box ubiquitin-protein ligase complexes. Acta Crystallogr D Biol Crystallogr 69, 1587–1597 (2013). 10.1107/S0907444913011220

49 Duda, D. M. et al. Structural insights into NEDD8 activation of cullin-RING ligases: conformational control of conjugation. Cell 134, 995–1006 (2008). 10.1016/j.cell.2008.07.022

50 Jardin, F. et al. Recurrent mutations of the exportin 1 gene (XPO1) and their impact on selective inhibitor of nuclear export compounds sensitivity in primary mediastinal B-cell lymphoma. Am J Hematol 91, 923–930 (2016). 10.1002/ajh.24451

51 Xie, Q. L., Liu, Y. & Zhu, Y. Chromosome region maintenance 1 expression and its association with clinical pathological features in primary carcinoma of the liver. Exp Ther Med 12, 59–68 (2016). 10.3892/etm.2016.3283

52 Hing, Z. A. et al. Exploring the Role of the Recurrent Exportin 1 (XPO1/CRM1) Mutations E571G and E571K in Chronic Lymphocytic Leukemia. Blood 128, 972–972 (2016). 10.1182/blood.V128.22.972.972

53 Navarro-Bailon, A. et al. Exportin-1 E571K mutation is a common finding in patients with classical Hodgkin lymphoma. Hematol Oncol 37, 215–218 (2019). 10.1002/hon.2570

54 Dong, X. et al. Structural basis for leucine-rich nuclear export signal recognition by CRM1. Nature 458, 1136–1141 (2009). 10.1038/nature07975

55 Port, S. A. et al. Structural and Functional Characterization of CRM1-Nup214 Interactions Reveals Multiple FG-Binding Sites Involved in Nuclear Export. Cell Rep 13, 690–702 (2015). 10.1016/j.celrep.2015.09.042

56 Koyama, M., Shirai, N. & Matsuura, Y. Structural insights into how Yrb2p accelerates the assembly of the Xpo1p nuclear export complex. Cell Rep 9, 983–995 (2014). 10.1016/j.celrep.2014.09.052

57 Fung, H. Y., Fu, S. C., Brautigam, C. A. & Chook, Y. M. Structural determinants of nuclear export signal orientation in binding to exportin CRM1. Elife 4 (2015). 10.7554/eLife.10034

58 Fung, H. Y., Fu, S. C. & Chook, Y. M. Nuclear export receptor CRM1 recognizes diverse conformations in nuclear export signals. Elife 6 (2017). 10.7554/eLife.23961

59 Lei, Y. et al. Structure-Guided Design of the First Noncovalent Small-Molecule Inhibitor of CRM1. J Med Chem 64, 6596–6607 (2021). 10.1021/acs.jmedchem.0c01675

60 Cao, S. et al. Defining molecular glues with a dual-nanobody cannabidiol sensor. Nat Commun 13, 815 (2022). 10.1038/s41467-022-28507-1

61 Skaar, J. R., Pagan, J. K. & Pagano, M. Mechanisms and function of substrate recruitment by F-box proteins. Nat Rev Mol Cell Biol 14, 369–381 (2013). 10.1038/nrm3582

62 Sheard, L. B. et al. Jasmonate perception by inositol-phosphate-potentiated COI1-JAZ co-receptor. Nature 468, 400–405 (2010). 10.1038/nature09430

63 Tan, X. et al. Mechanism of auxin perception by the TIR1 ubiquitin ligase. Nature 446, 640–645 (2007). 10.1038/nature05731

64 Ito, T. et al. Identification of a primary target of thalidomide teratogenicity. Science 327, 1345–1350 (2010). 10.1126/science.1177319

65 Fischer, E. S. et al. Structure of the DDB1-CRBN E3 ubiquitin ligase in complex with thalidomide. Nature 512, 49–53 (2014). 10.1038/nature13527

66 Du, X. et al. Structural Basis and Kinetic Pathway of RBM39 Recruitment to DCAF15 by a Sulfonamide Molecular Glue E7820. Structure 27, 1625–1633 e1623 (2019). 10.1016/j.str.2019.10.005

67 Slabicki, M. et al. The CDK inhibitor CR8 acts as a molecular glue degrader that depletes cyclin K. Nature 585, 293–297 (2020). 10.1038/s41586-020-2374-x

68 Slabicki, M. et al. Small-molecule-induced polymerization triggers degradation of BCL6. Nature 588, 164–168 (2020). 10.1038/s41586-020-2925-1

69 Azmi, A. S. et al. Selective inhibitors of nuclear export block pancreatic cancer cell proliferation and reduce tumor growth in mice. Gastroenterology 144, 447–456 (2013). 10.1053/j.gastro.2012.10.036

70 Gravina, G. L. et al. KPT-330, a potent and selective exportin-1 (XPO-1) inhibitor, shows antitumor effects modulating the expression of cyclin D1 and survivin [corrected] in prostate cancer models. BMC Cancer 15, 941 (2015). 10.1186/s12885-015-1936-z

71 Hilliard, M. et al. The anti-inflammatory prostaglandin 15-deoxy-delta(12,14)-PGJ2 inhibits CRM1-dependent nuclear protein export. J Biol Chem 285, 22202–22210 (2010). 10.1074/jbc.M110.131821

72 Ribo-Molina, P. et al. 4-Octyl itaconate reduces influenza A replication by targeting the nuclear export protein CRM1. J Virol 97, e0132523 (2023). 10.1128/jvi.01325-23

73 Waqas, F. H. et al. NRF2 activators inhibit influenza A virus replication by interfering with nucleo-cytoplasmic export of viral RNPs in an NRF2-independent manner. PLoS Pathog 19, e1011506 (2023). 10.1371/journal.ppat.1011506

74 Waqas, F. H. et al. [Pre-print] NRF2 activators restrict coronaviruses by targeting a network involving ACE2, TMPRSS2, and XPO1. bioRxiv, 2025.2002.2024.639813 (2025). 10.1101/2025.02.24.639813

75 Wang, P. et al. Repression of classical nuclear export by S-nitrosylation of CRM1. J Cell Sci 122, 3772–3779 (2009). 10.1242/jcs.057026

76 Boons, E. et al. Human Exportin-1 is a Target for Combined Therapy of HIV and AIDS Related Lymphoma. EBioMedicine 2, 1102–1113 (2015). 10.1016/j.ebiom.2015.07.041

77 Van Hauwenhuyse, J. et al. Abstract 1652: A novel reversible inhibitor of XPO1 with potent efficacy in multiple preclinical mouse models. Cancer Research 83, 1652-1652 (2023). 10.1158/1538-7445.Am2023-1652

78 Dimopoulos, M. A. et al. Carfilzomib and dexamethasone versus bortezomib and dexamethasone for patients with relapsed or refractory multiple myeloma (ENDEAVOR): a randomised, phase 3, open-label, multicentre study. Lancet Oncol 17, 27–38 (2016). 10.1016/S1470-2045(15)00464-7

79 Turner, J. G. et al. XPO1 inhibitor combination therapy with bortezomib or carfilzomib induces nuclear localization of IkappaBalpha and overcomes acquired proteasome inhibitor resistance in human multiple myeloma. Oncotarget 7, 78896–78909 (2016). 10.18632/oncotarget.12969

80 Galinski, B. et al. XPO1 inhibition with selinexor synergizes with proteasome inhibition in neuroblastoma by targeting nuclear export of IkB. Transl Oncol 14, 101114 (2021). 10.1016/j.tranon.2021.101114

81 Schneider, C. A., Rasband, W. S. & Eliceiri, K. W. NIH Image to ImageJ: 25 years of image analysis. Nat Methods 9, 671–675 (2012). 10.1038/nmeth.2089

82 Fung, H. Y. J. & Chook, Y. M. Binding Affinity Measurement of Nuclear Export Signal Peptides to Their Exporter CRM1. Methods Mol Biol 2502, 245–256 (2022). 10.1007/978-1-0716-2337-4_16

83 Keller, S. et al. High-precision isothermal titration calorimetry with automated peak-shape analysis. Anal Chem 84, 5066–5073 (2012). 10.1021/ac3007522

84 Houtman, J. C. et al. Studying multisite binary and ternary protein interactions by global analysis of isothermal titration calorimetry data in SEDPHAT: application to adaptor protein complexes in cell signaling. Protein Sci 16, 30–42 (2007). 10.1110/ps.062558507

85 Brautigam, C. A. Calculations and Publication-Quality Illustrations for Analytical Ultracentrifugation Data. Methods Enzymol 562, 109–133 (2015). 10.1016/bs.mie.2015.05.001

86 Mastronarde, D. N. Automated electron microscope tomography using robust prediction of specimen movements. J Struct Biol 152, 36–51 (2005). 10.1016/j.jsb.2005.07.007

87 Punjani, A., Rubinstein, J. L., Fleet, D. J. & Brubaker, M. A. cryoSPARC: algorithms for rapid unsupervised cryo-EM structure determination. Nat Methods 14, 290-+ (2017). 10.1038/Nmeth.4169

88 Bepler, T. et al. Positive-unlabeled convolutional neural networks for particle picking in cryo-electron micrographs. Nat Methods 16, 1153–1160 (2019). 10.1038/s41592-019-0575-8

89 Pettersen, E. F. et al. UCSF chimera – A visualization system for exploratory research and analysis. J Comput Chem 25, 1605–1612 (2004). 10.1002/jcc.20084

90 Pettersen, E. F. et al. UCSF ChimeraX: Structure visualization for researchers, educators, and developers. Protein Sci 30, 70–82 (2021). 10.1002/pro.3943

91 Adams, P. D. et al. PHENIX: a comprehensive Python-based system for macromolecular structure solution. Acta Crystallogr D 66, 213–221 (2010). 10.1107/S0907444909052925

92 Croll, T. I. ISOLDE: a physically realistic environment for model building into low-resolution electron-density maps. Acta Crystallogr D Struct Biol 74, 519–530 (2018). 10.1107/S2059798318002425

93 Emsley, P. New Tools for Ligand Refinement and Validation in Coot and CCP4. Acta Crystallogr A 74, A390–A390 (2018). 10.1107/S0108767318096101

94 Sanchez-Garcia, R. et al. DeepEMhancer: a deep learning solution for cryo-EM volume post-processing. Commun Biol 4, 874 (2021). 10.1038/s42003-021-02399-1

95 Schrodinger, LLC. The PyMOL Molecular Graphics System, Version 2.*4* (2020).

96 Krissinel, E. & Henrick, K. Inference of macromolecular assemblies from crystalline state. J Mol Biol 372, 774–797 (2007). 10.1016/j.jmb.2007.05.022

97 Jurrus, E. et al. Improvements to the APBS biomolecular solvation software suite. Protein Sci 27, 112–128 (2018). 10.1002/pro.3280

98 Fung, H. Y. J. & Chook, Y. M. Crystallization of Nuclear Export Signals or Small-Molecule Inhibitors Bound to Nuclear Exporter CRM1. Methods Mol Biol 2502, 285–297 (2022). 10.1007/978-1-0716-2337-4_19

99 Moriarty, N. W., Grosse-Kunstleve, R. W. & Adams, P. D. electronic Ligand Builder and Optimization Workbench (eLBOW): a tool for ligand coordinate and restraint generation. Acta crystallographica. Section D, Biological crystallography 65, 1074–1080 (2009). 10.1107/S0907444909029436

100 Afonine, P. V. et al. Towards automated crystallographic structure refinement with phenix.refine. *Acta crystallographica. Section D*, Biological crystallography 68, 352–367 (2012). 10.1107/S0907444912001308

101 Emsley, P., Lohkamp, B., Scott, W. G. & Cowtan, K. Features and development of Coot. Acta crystallographica. Section D, Biological crystallography 66, 486–501 (2010). 10.1107/S0907444910007493

102 Doench, J. G. et al. Optimized sgRNA design to maximize activity and minimize off-target effects of CRISPR-Cas9. Nat Biotechnol 34, 184–191 (2016). 10.1038/nbt.3437

103 Sanjana, N. E., Shalem, O. & Zhang, F. Improved vectors and genome-wide libraries for CRISPR screening. Nat Methods 11, 783–784 (2014). 10.1038/nmeth.3047

104 Jung, M. E. & Buszek, K. R. The stereochemistry of addition of trialkylammonium and pyridinium tetrafluoroborate salts to activated acetylenes. Preparation of novel dienophiles for Diels-Alder reactions. Journal of the American Chemical Society 110, 3965–3969 (1988). 10.1021/ja00220a039

105 Vercruysse, T. et al. Ibetazol, a novel inhibitor of importin beta1-mediated nuclear import. Commun Biol 7, 1560 (2024). 10.1038/s42003-024-07237-8

