## Extended Data for "SINE compounds activate exportin-1 degradation via an allosteric mechanism"

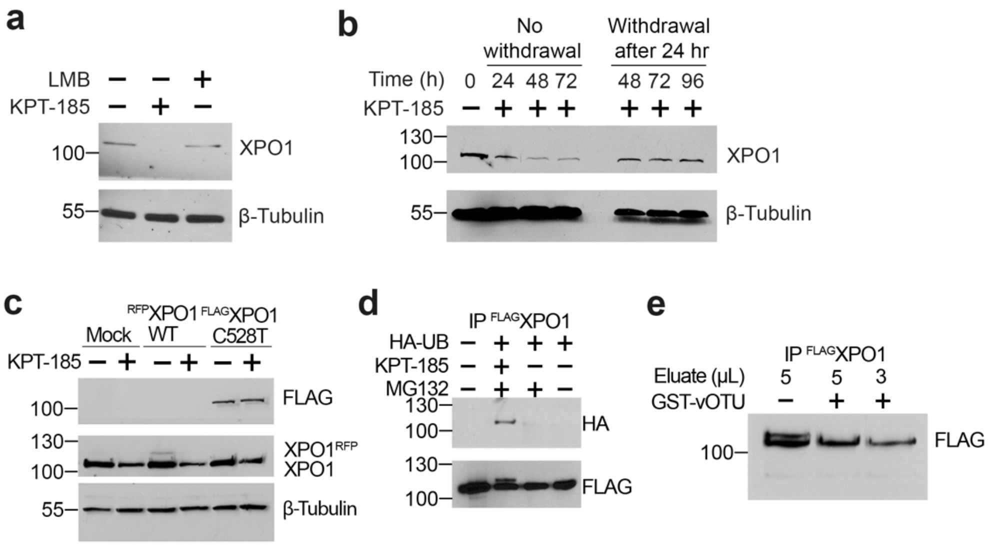


**Extended Data Figure 1. KPT-185-induced XPO1 degradation: specificity and XPO1 ubiquitination.**

**a,** XPO1 levels in HT1080 cells treated with KPT-185, LMB or DMSO detected by Western blot (WB). Loading control in **a-c** is β-tubulin.

**b**, HT1080 cells, treated with DMSO or KPT-185 for 24 h followed by fresh media with or without KPT-185, were collected at 24, 48, 72 and 96 h for detection of XPO1 levels by WB.

**c**, HT1080 cells transfected with $^{RFP}$XPO1$^{WT}$ or $^{FLAG}$XPO1$^{C528T}$ were treated with DMSO or KPT-185; XPO1 detected by WB.

**d,** HEK-293$^{FLAG\text{-}XPO1}$ stable cells transfected with HA-ubiquitin (HA-UB) were treated with DMSO or KPT-185 followed by MG132. $^{FLAG}$XPO1 immunoprecipitated (IP-ed) from cell lysates were subjected to WB with anti-HA antibody.

**e,** $^{FLAG}$XPO1, IP-ed from HEK-293$^{FLAG\text{-}XPO1}$ cells treated with selinexor and MG132, was treated with GST-vOTU deubiquitinase. WB shows disappearance of the higher molecular weight $^{FLAG}$XPO1 band after deubiquitinase treatment.

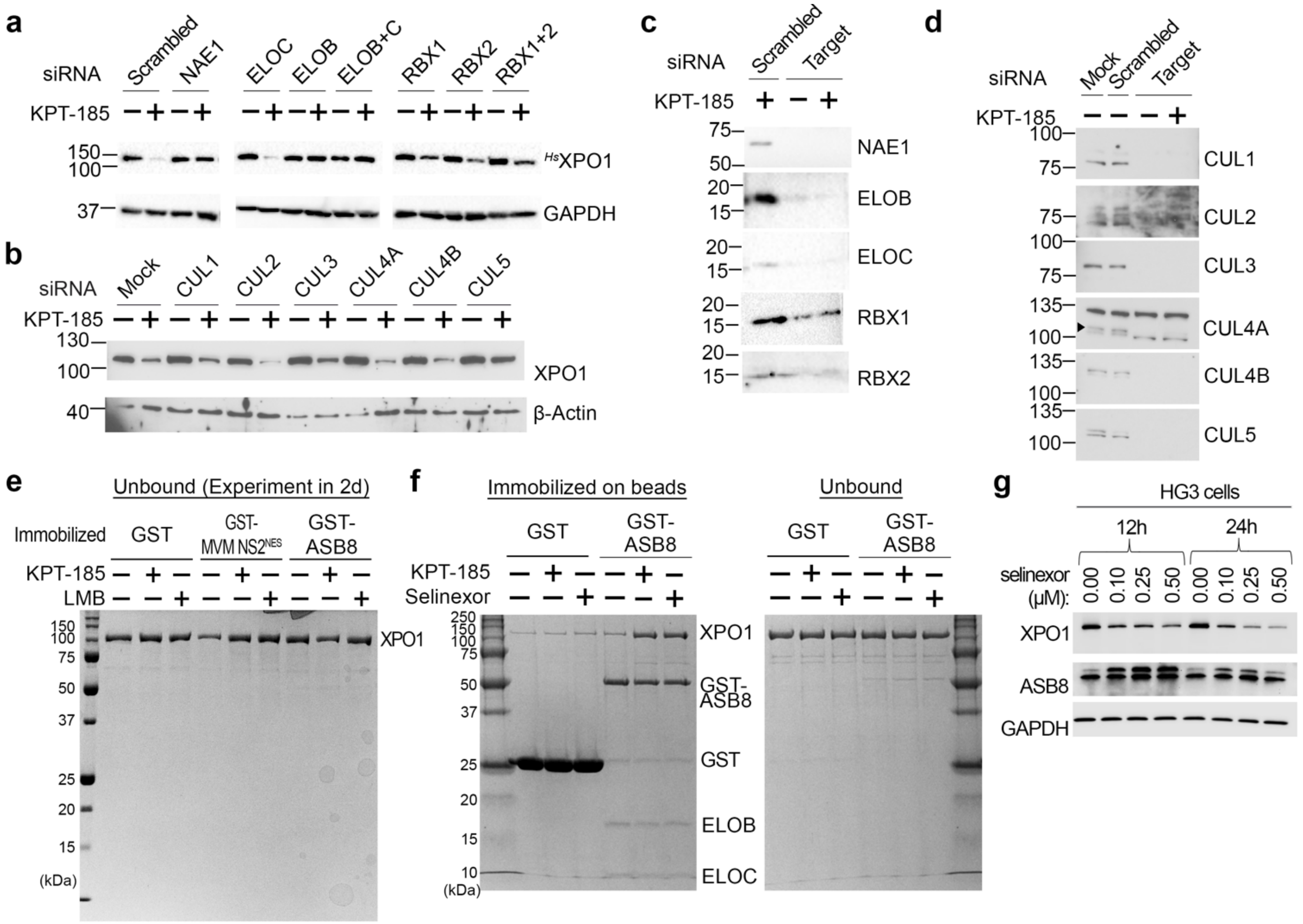


**Extended Data Figure 2. SINE-induced degradation of XPO1 is mediated by CRL5$^{ASB8}$.**

- **a**, XPO1 level in HT1080 cells treated with KPT-185 and siRNAs of various CRL5$^{ASB8}$ components. Detection by WB, loading control is GAPDH and positive control is NAE1 siRNA.
- **b,** HT1080 cells treated with siRNA for several cullins followed by treatment with DMSO or KPT-185. Loading control is β-actin.
- **c,** Cells from **(a)** probed with antibodies for the respective the siRNA targets.
- **d,** Cells from **(b)** probed with antibodies for the respective the siRNA targets.
- **e,** Unbound proteins from pull-down assay shown in **Figure 2c** of XPO1 treated with DMSO, KPT-185, LMB binding to immobilized GST-ASB8-ELOB/C (GST is negative control, and GST-MVM NS2$^{NES}$ is positive control).
- **f,** Pull-down assay of immobilized GST-ASB8•ELOB/C (negative control is GST) with XPO1 pre-treated with KPT-185 or selinexor. Bound and unbound proteins are visualized by Coomassie SDS-PAGE.
- **g,** Expression level of XPO1 and ASB8 by WB in HG3 cells treated with DMSO or selinexor. Cells were treated at 0-0.5 µM selinexor for 12 h or 24 h. Loading control is GAPDH.

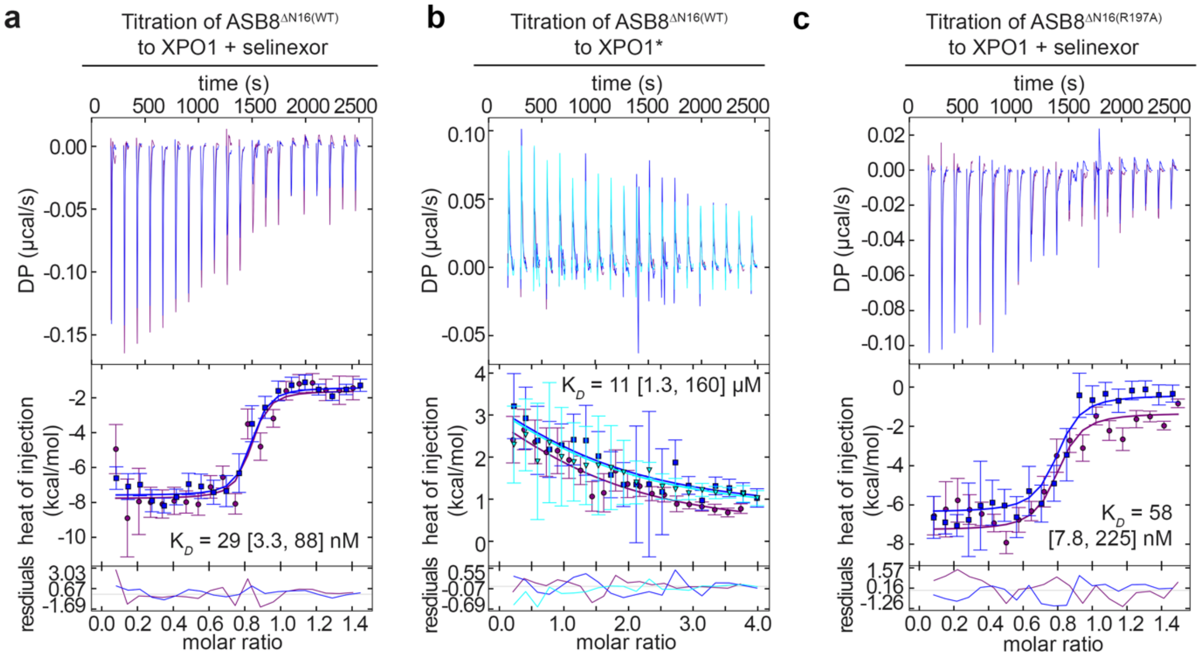


**Extended Data Figure 3. ITC analysis of XPO1 binding to CRL5$^{ASB8}$.**

**a-c,** ITC titration of ASB8$^{\Delta N16(WT)}$ binding to XPO1 in the presence of selinexor (**a**), ASB8$^{\Delta N16(WT)}$ binding to XPO1 in the absence of selinexor (**b**), or ASB8$^{\Delta N16(R197A)}$ binding to XPO1 in the presence of selinexor (**c**). Reactions with selinexor (**a** and **c**) were performed at 20 °C, while reactions without selinexor (**b**) were performed at 10 °C (indicated with an asterisk). The top panels show reconstructed thermograms from NITPIC, the middle panels show binding isotherms and individual fits (error bars are estimated error of peak integration performed by NITPIC), and the bottom panels show the fitting residuals. Dissociation constants ($K_D$s) obtained from global analysis of duplicate (**a** and **c**) or triplicate (**b**) measurements are displayed with 95% confidence intervals in brackets. ASB8$^{\Delta N16(WT)}$ binds to selinexor-XPO1 with a $K_D$ of 29 nM (**a**), while the affinity is much weaker ($K_D$ = 11 μM) in the absence of SINEs (**b**). ASB8$^{\Delta N16(R197A)}$ binds selinexor-XPO1 with a similar affinity as WT ASB8, with a $K_D$ of 58 nM (**c**).

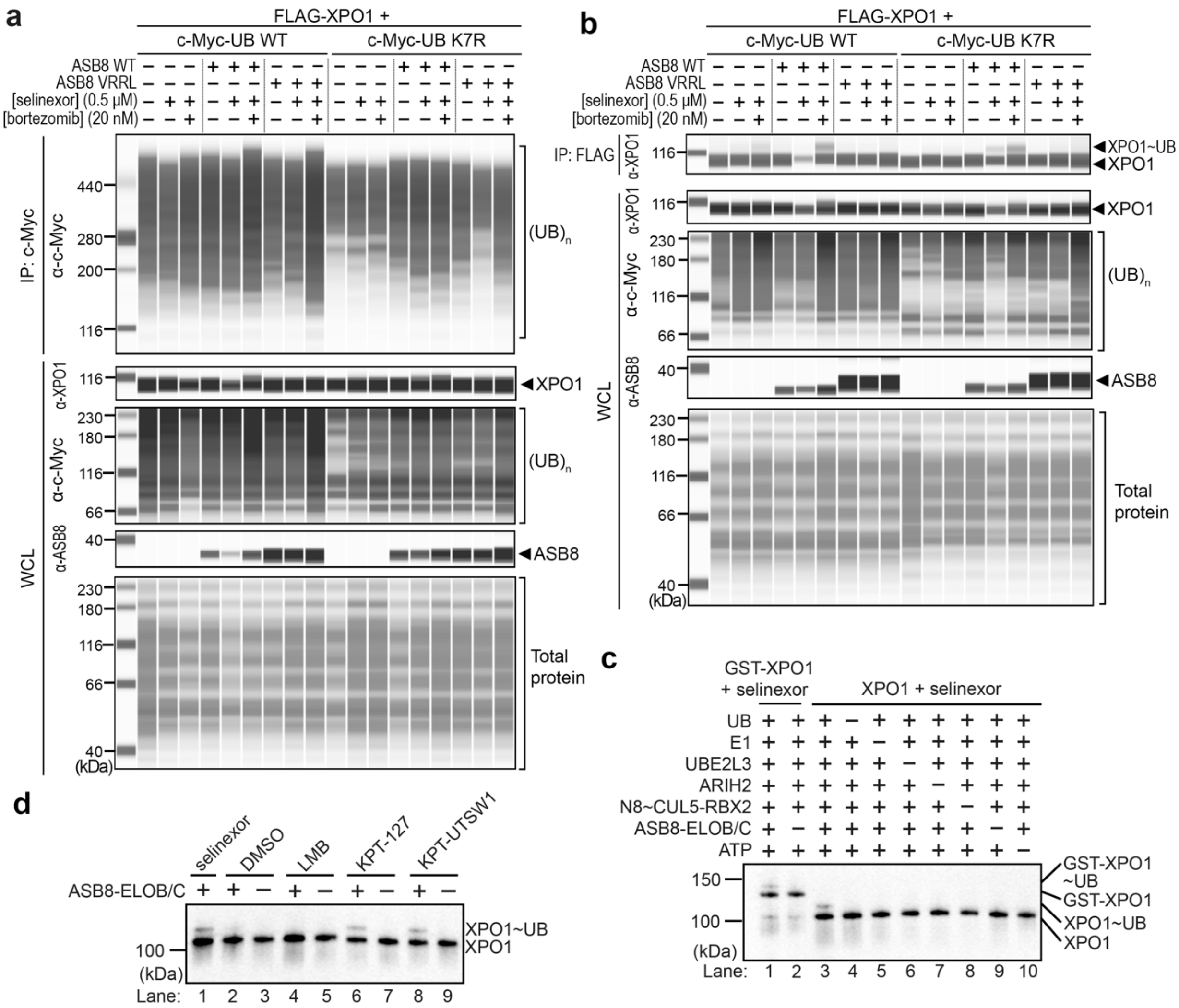


**Extended Data Figure 4. SINE-induced ubiquitination of XPO1 is mediated by $CRL5^{ASB8}$.**

**a,b,** ASB8-mediated ubiquitination of XPO1 was examined by expressing pDNA constructs in HEK293T cells as indicated, followed by 24 h of incubation and 6 h of treatment with selinexor and bortezomib. Cells were collected and lysed (whole-cell lysate, WCL) after which c-Myc (**a**) or FLAG (**b**) fusions were pulled down and detected via WB. Blots serve as supplementary material for **Figure 2d**.

**c,** *In vitro* mono-ubiquitination of XPO1 by ARIH2 and UBE2L3, visualized by XPO1 antibody in via WB. A series of reactions (lanes 3-10) were assembled with ubiquitin, E1, ATP-Mg, UBE2L3, ARIH2, neddylated CUL5-RBX2 (N8~CUL5-RBX2), ASB8(FL)-ELOB/C and XPO1 pre-incubated with selinexor, by omitting each individual component. Absence of XPO1~UB species in lanes 4-10 indicate that all components are necessary for ubiquitination of XPO1-selinexor. Similar reactions with and without ASB8-ELOB/C were also assembled with GST-XPO1 instead of untagged XPO1 (lanes 1-2), indicating that the band shift due to ubiquitination is less than that of GST, which is ~25 kDa, and is likely mono-ubiquitination.

**d,** Like the assay in (**c**), but XPO1 was pre-incubated with selinexor, DMSO, Leptomycin B (LMB), KPT-127 or KPT-UTSW1. There is very minor ubiquitination when no inhibitor is present (lane 2), but LMB, which blocks the entire NES groove, prevents ASB8 interaction and thus ubiquitination (lane 4). Selinexor, KPT-127 and KPT-UTSW1 can all enable high affinity ASB8-binding and cause mono-ubiquitination by ARIH2/UBE2L3.

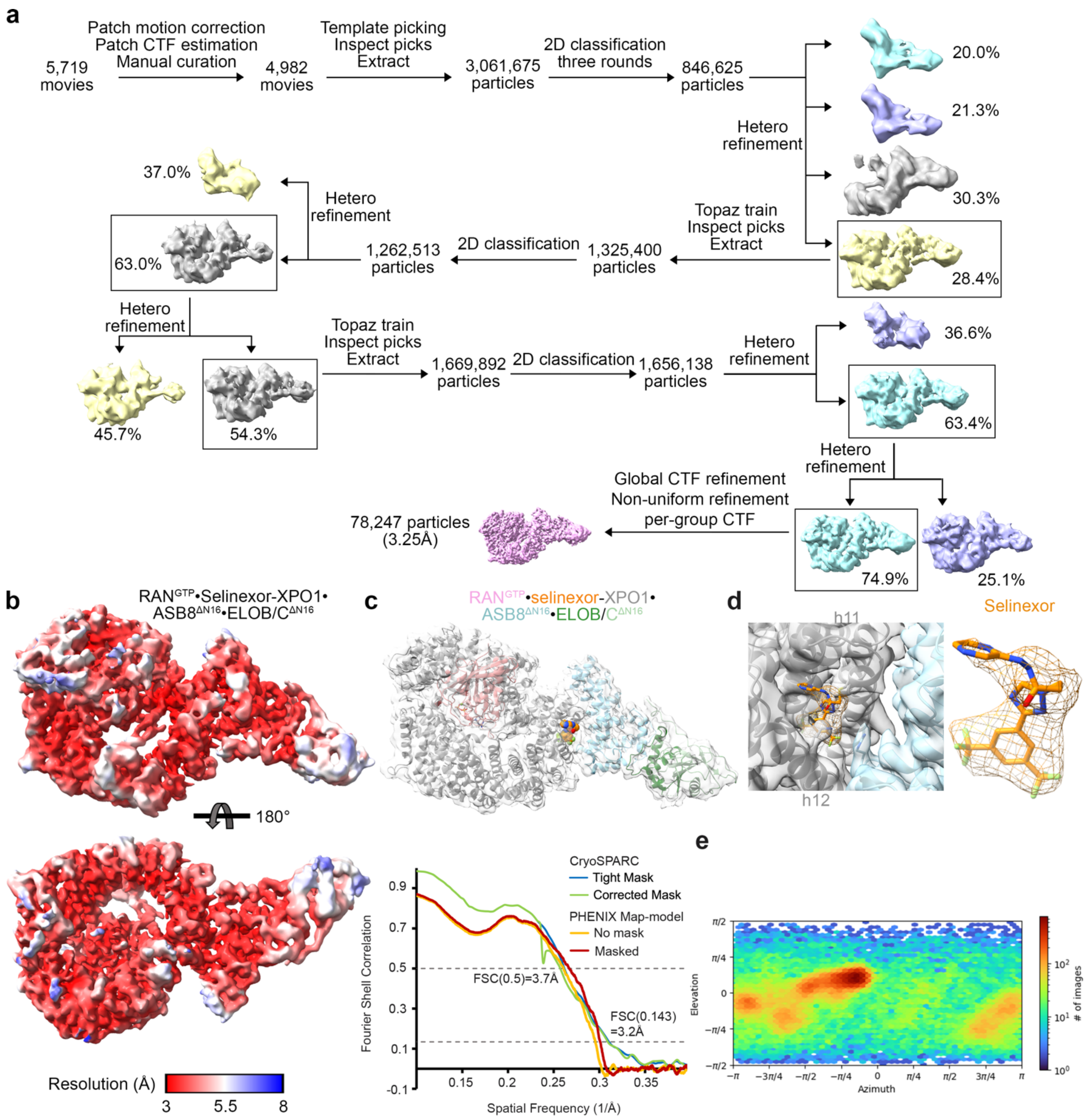


**Extended Data Figure 5. CryoEM maps and statistics for RAN$^{GTP}$•selinexor-XPO1•ASB8$^{\Delta N16}$•ELOB/C**

**a,** The flowchart of single particle analysis of the RAN$^{GTP}$•selinexor-XPO1•ASB8$^{\Delta N16}$•ELOB/C complex.

**b,** The final map of RAN$^{GTP}$•selinexor-XPO1•ASB8$^{\Delta N16}$•ELOB/C colored by local resolution as indicated by the scale.

**c,** The refined RAN$^{GTP}$•selinexor-XPO1•ASB8$^{\Delta N16}$•ELOB/C structure overlayed with the map in (**b**), accompanied by cryoSPARC and PHENIX map-model FSCs on the bottom.

**d,** A view of the selinexor-bound XPO1 groove (gray cartoon and transparent gray map). Selinexor is shown as orange stick with the cryoEM map around the inhibitor shown as mesh. ASB8 cartoon and map in transparent cyan.

**e,** Viewing direction distribution plot for the particles used in reconstruction for the map in (**b**).

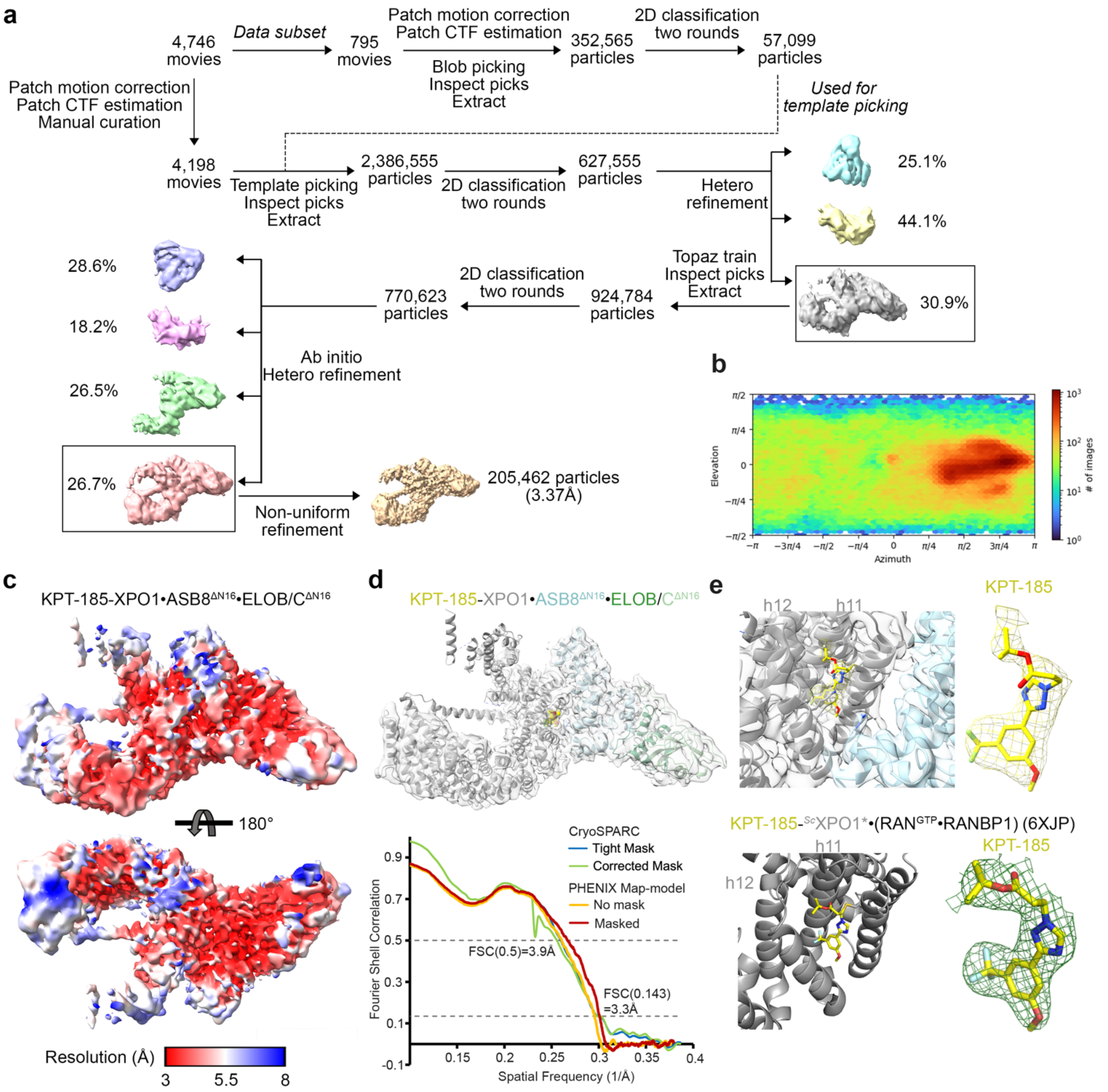


**Extended Data Figure 6. CryoEM map and statistics for KPT-185-XPO1•ASB8$^{\Delta N16}$•ELOB/C.**

**a,** The flowchart of single particle analysis of the KPT-185-XPO1•ASB8$^{\Delta N16}$•ELOB/C complex.

**b,** Viewing direction distribution plot for the particles used in reconstruction for the map in (**c**).

**c,** The final map of KPT-185-XPO1•ASB8$^{\Delta N16}$•ELOB/C colored by local resolution as indicated by the scale.

**d,** The refined KPT-185-XPO1•ASB8$^{\Delta N16}$•ELOB/C structure overlayed with the map in (**c**), accompanied by cryoSPARC and PHENIX map-model FSCs on the bottom.

**e,** Top: a view of the KPT-185-bound groove of XPO1 (gray cartoon and transparent gray map). KPT-185 is shown as yellow stick with the map around shown as mesh. ASB8 cartoon and map in cyan. Bottom: a similar view of the KPT-185-bound groove of XPO1 of the crystal structure of KPT-185-$^{Sc}$XPO1*•RAN$^{GTP}$•RANBP1 (6XJP), with the 2mFo-Fc density at 1σ (green mesh) shown at KPT-185.

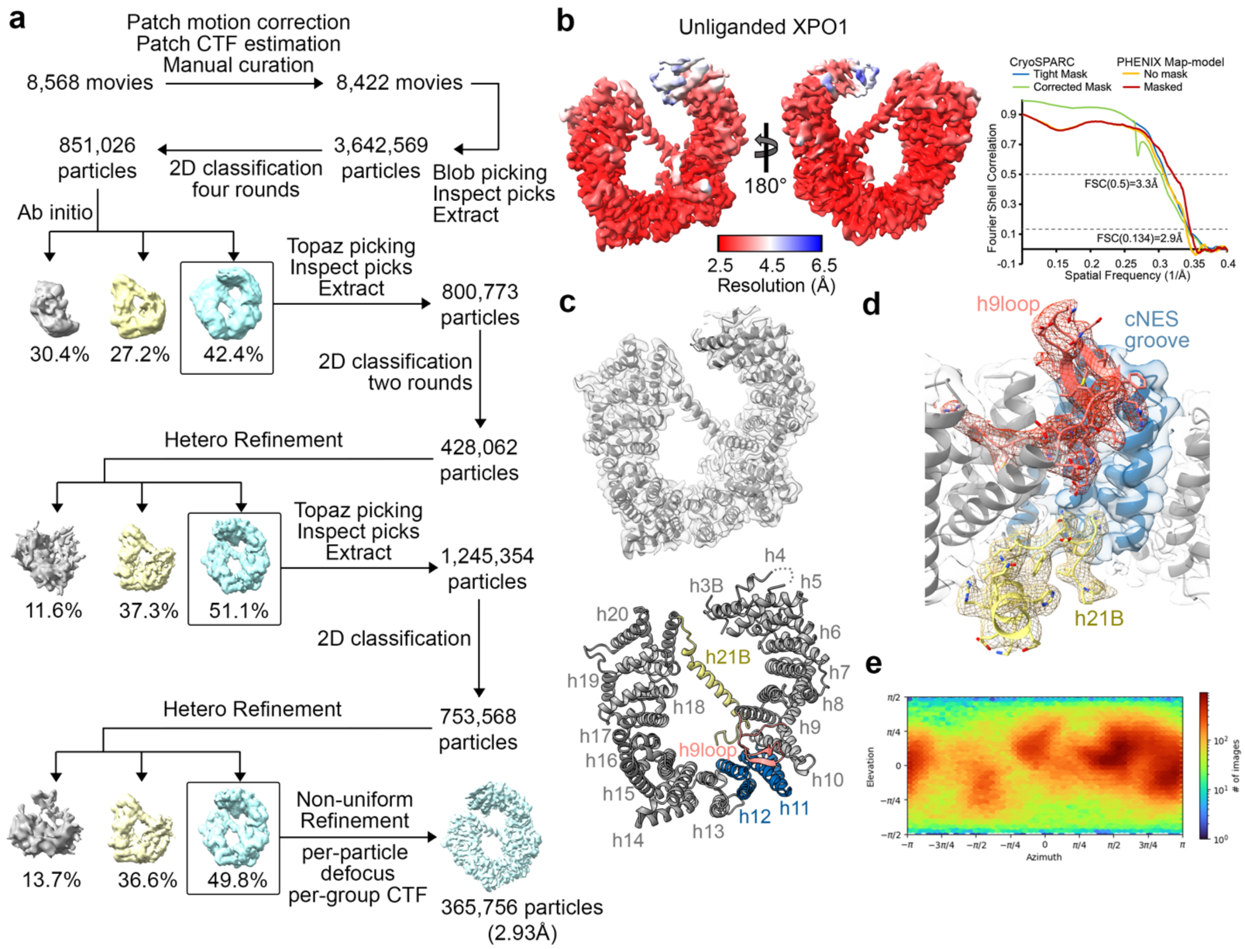


**Extended Data Figure 7. CryoEM maps and statistics for unliganded XPO1.**

**a,** The flowchart of single particle analysis of unliganded XPO1.

**b,** The final map of unliganded XPO1 colored by local resolution as indicated by the scale, with the cryoSPARC and PHENIX generated map-model FSC curves on the right.

**c,** Structure of unliganded XPO1. Top panel, the refined structure of unliganded XPO1 in the final map. Bottom panel, the structure shown as cartoon, with the 21 HEAT repeats labeled.

**d,** Density (mesh) of the h21B helix and C-terminal tail (yellow) and of the h9loop (red). The h21B-C-terminal tail stretch and the h9loop interact with the back (or concave side) of the cNES binding groove (shown in blue) to stabilize the groove in its closed conformation.

**e,** The viewing direction distribution plot for the particles used in reconstruction for the map in (**b**).

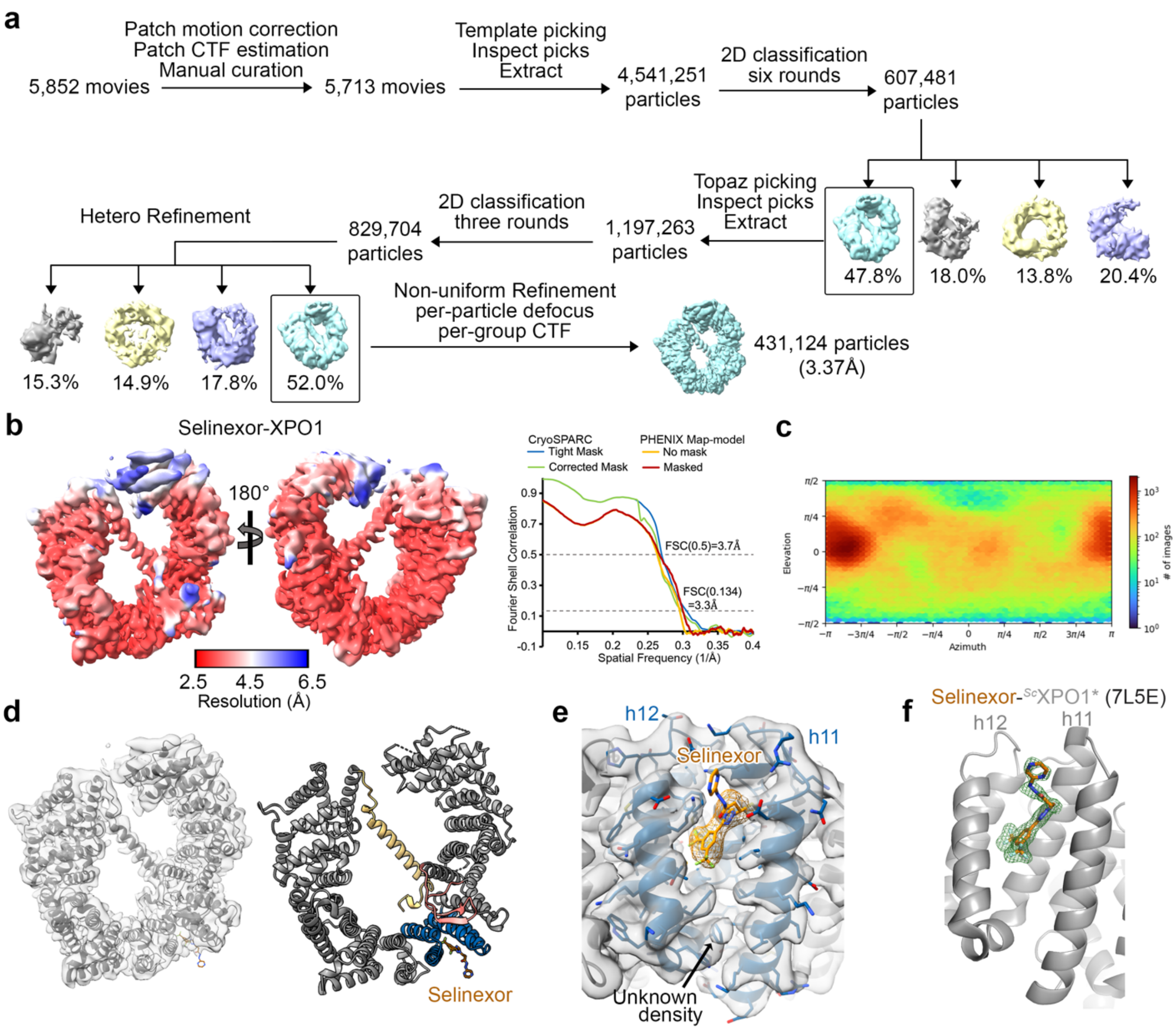


**Extended Data Figure 8. CryoEM maps and statistics for selinexor-XPO1.**

**a,** The flowchart of single particle analysis of selinexor-XPO1.

**b,** The final map of selinexor-XPO1 colored by local resolution, with the cryoSPARC and PHENIX generated map-model FSC curves shown on right.

**c,** Viewing direction distribution plot for the particles used in reconstruction for the map in (**b**).

**d**, The selinexor-XPO1 structure. Left, the refined structure of selinexor-XPO1 in the final map. Right, cartoon of XPO1 and selinexor shown as brown sticks.

**e,** The cryoEM map at the selinexor-bound groove. The density (mesh) for the KPT-185 scaffold of selinexor is well-defined, but density for the pyrazinylpropenehydrazide extension is weak. Extra density in the groove for an unknown molecule (possibly a water molecule or an ion) is marked with a black arrow.

**f,** The crystal structure of the selinexor-$^{Sc}$XPO1*•RAN$^{GTP}$•RANBP1 complex (7L5E), showing the 2mFo-Fc map at 1σ for selinexor (green mesh). The different orientation of the pyrazinylpropenehydrazide arm here compared to in panel **e** above, suggests flexibility.

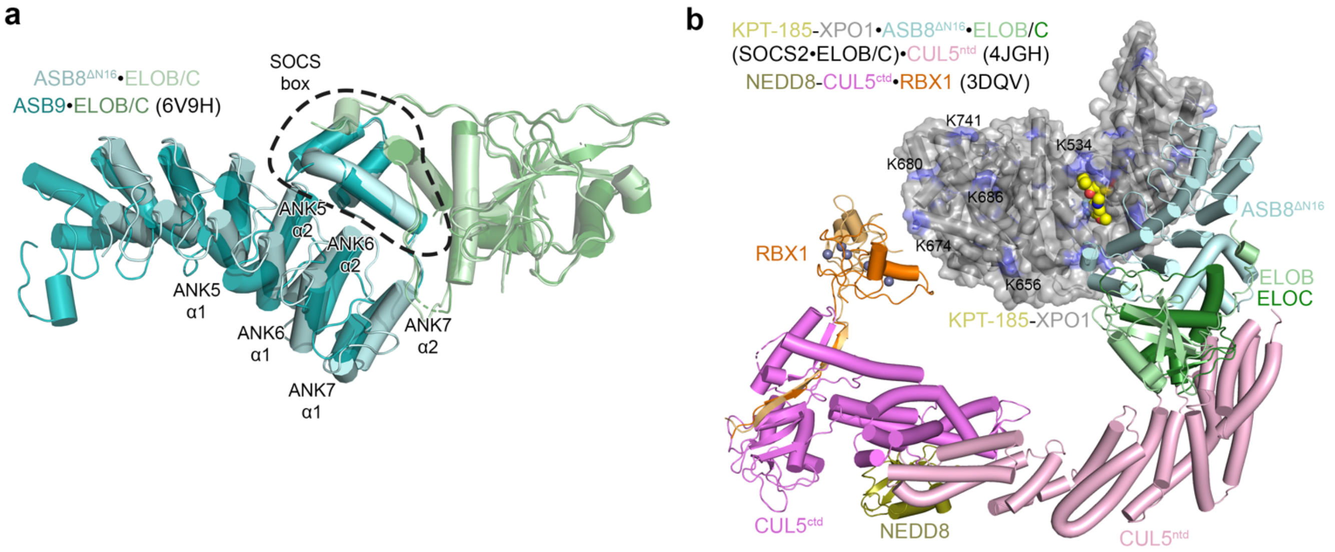


**Extended Data Figure 9. KPT-185-XPO1•ASB8$^{\Delta N16}$•ELOB/C: structural alignments.**

**a,** Alignment of ASB8 and ASB9. ELOB/C (dark green) bound to ASB9 (dark teal) from 6V9H was aligned to ELOB/C (light green) bound to the ASB8 (pale cyan) in our KPT-185-XPO1•ASB8$^{\Delta N16}$•ELOB/C cryoEM structure (0.809 Å r.m.s.d.). The α1 and α2 helices of ANK5-7 are labeled, and the SOCS boxes are marked with a dashed line.

**b,** A model of KPT-185-XPO1•CRL5$^{ASB8}$ generated by a series of alignments. The cryoEM structure of KPT-185-XPO1•ASB8•ELOB/C was first aligned with ELOB/C-CUL5$^{ntd}$ (4JGH) (r.m.s.d 1.151 Å). The overlayed ELOB/C-CUL5$^{ntd}$ was then aligned to CUL5$^{FL}$ (7ONI, r.m.s.d. 0.843 Å) to position the full length CUL5 into the model. Finally, the four-helix bundle and α/β subdomains (CUL5 residues 401-687) of the overlayed CUL5$^{FL}$ (7ONI) and the NEDD8-CUL5$^{ctd}$•RBX1 (3DQV) were aligned (r.m.s.d. 1.143 Å) to model in the neddylated CUL5 and the released/mobile RBX1 (copies from both asymmetric units shown). The mobile RBX1/2 is expected to allow the associated E2 enzyme to reach exposed lysine residues of XPO1.

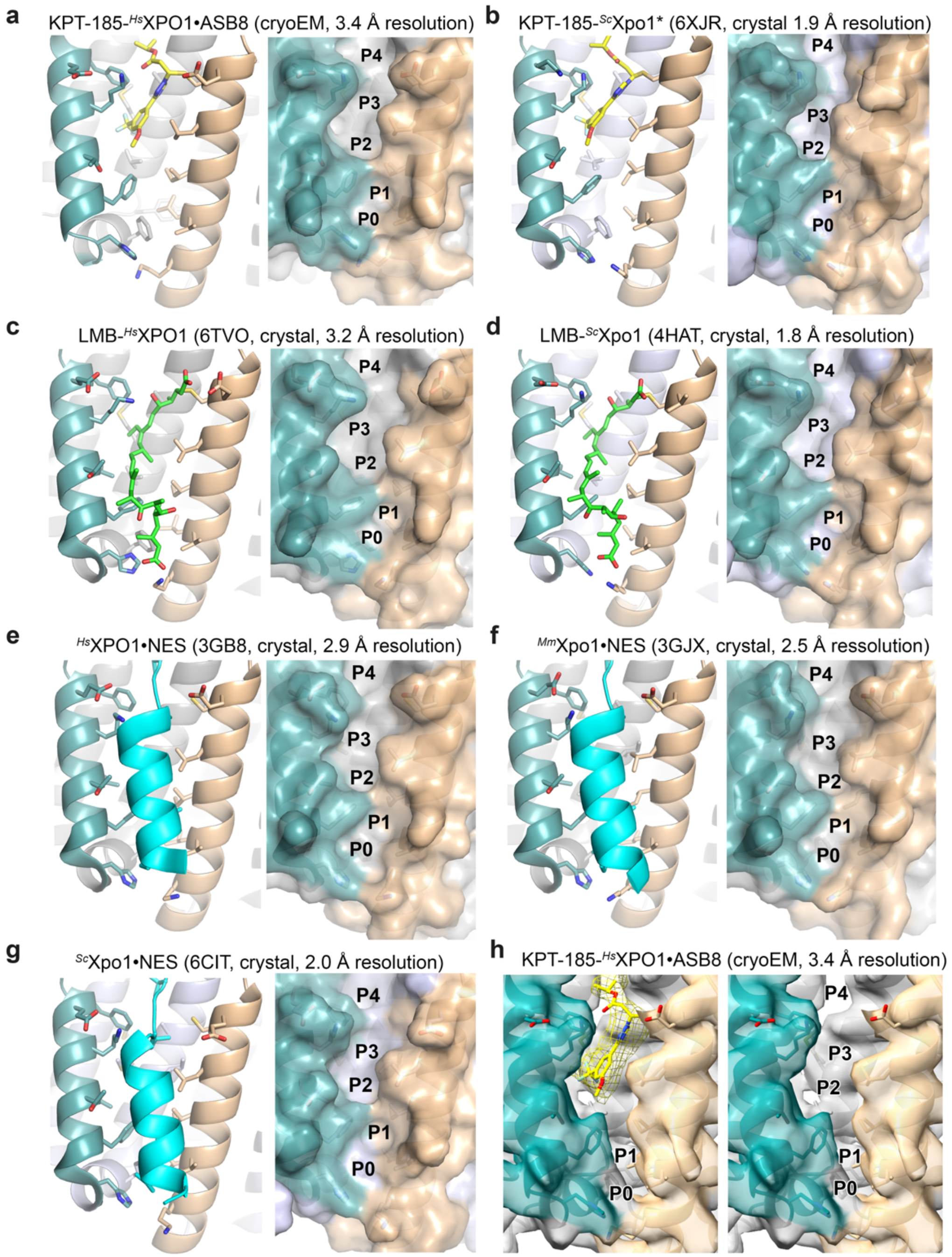


**Extended Data Figure 10. Comparing open XPO1 grooves.**

**a-g,** Cartoons with side chain sticks (left) and surface representations (right) of the XPO1 groove are shown in different colors: h11A and h12A helices (front) are in brown and teal, respectively, while the h11B and h12B helices (back) are in gray. When the groove is open, the gray back helices should be visible. Surface representations are displayed without ligands to show the open grooves, with hydrophobic pockets P0-P4 labeled on the right. (**a-d**) Inhibitor-bound XPO1 grooves**: a,** the KPT-185-$^{Hs}$XPO1•ASB8•ELOB/C cryoEM structure (ASB8 not displayed); **b,** the crystal structure of KPT-185-$^{Sc}$Xpo1* (* refers to humanized groove)**; c,** the crystal structure of LMB-$^{Hs}$XPO1 and **d,** the crystal structure of LMB-$^{Sc}$XPO1**.** (**e-g**) NES-bound XPO1 grooves (only NES peptides is displayed, in cyan)**: e,** crystal structure of cargo SPN1 bound to human XPO1; **f,** the crystal structure of RAN$^{GTP}$•mouse Xpo1•SPN1 complex; **g,** the crystal structure of a RanBP1•Ran$^{GTP}$•$^{Sc}$XPO1*•MVM NS2 NES complex**.**

**h,** CryoEM map of the structure in panel **a** is shown instead of surface representation of the model, colored by the overlayed model in the same coloring scheme as (**a**).

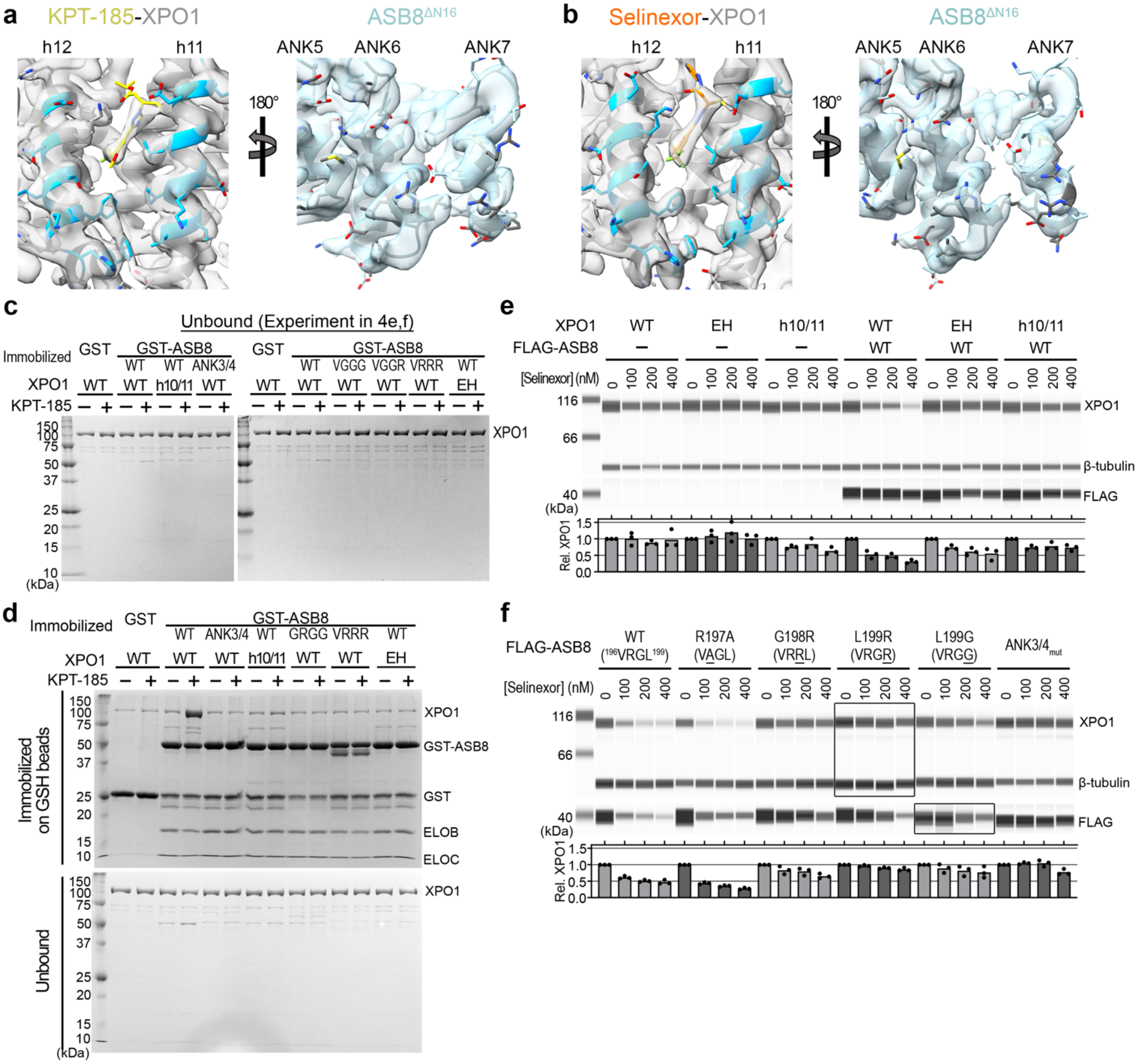


**Extended Data Figure 11. CryoEM Map quality and binding assays of GST-ASB8•ELOB/C with SINE-XPO1 to study the XPO1-ASB8 interfaces.**

**a-b,** CryoEM maps of (**a**) KPT-185-XPO1•ASB8$^{\Delta N16}$•ELOB/C and (**b**) RAN$^{GTP}$•selinexor-XPO1•ASB8$^{\Delta N16}$•ELOB/C shown at the XPO1 (h11/12)-ASB8 (ANK6/7) interfaces of both cryoEM structures, zoomed in, with the same view as in **Figure 3c**. XPO1 maps and cartoons are colored gray, with KPT-185 and selinexor are drawn as yellow and orange sticks, and XPO1 residues that interact with ASB8 as sky blue sticks. ASB8 maps and cartoons are light blue, with residues that interact with XPO1 drawn as dark gray sticks. Interface residues were reported by PDBePISA.

**c,** Unbound proteins from pull-down assay in **Figure 3e,f** of XPO1 (WT and mutants) ± KPT-185 to immobilized GST-ASB8-ELOB/C mutants. Gels were visualized by Coomassie staining.

**d,** Pull-down assay of immobilized GST-ASB8•ELOB/C mutants with XPO1 ± KPT-185, visualized by Coomassie staining. Mutations target the h10/11-ANK3/4 and the h11/12-ANK6/7 interfaces, similar to **Figure 4e,f**, but with ASB8$^{GRGG}$ instead of ASB8$^{VGGG}$ or ASB8$^{VGGR}$.

**e,** XPO1 degradation assay from **Figure 4g** with additional selinexor concentrations. HEK293T cells were transfected with XPO1$^{WT}$, XPO1$^{EH}$ or XPO1$^{h10/11}$ alone or together with ASB8 WT and treated overnight with selinexor. XPO1 degradation was assessed by WB.

**f,** XPO1 degradation assay from **Figure 11f** with additional selinexor concentrations. HEK293T cells were transfected with ASB8 WT ($^{196}$VRGL$^{199}$), ASB8$^{ANK3/4}$ and ASB8 mutants VAGL, VRRL, VRGR or VRGG followed by selinexor treatment overnight and WB analysis of XPO1. Representative WB images in **e** and **f.** Bar plots show the average XPO1/β-tubulin peak area from three independent experiments relative to untreated cells.

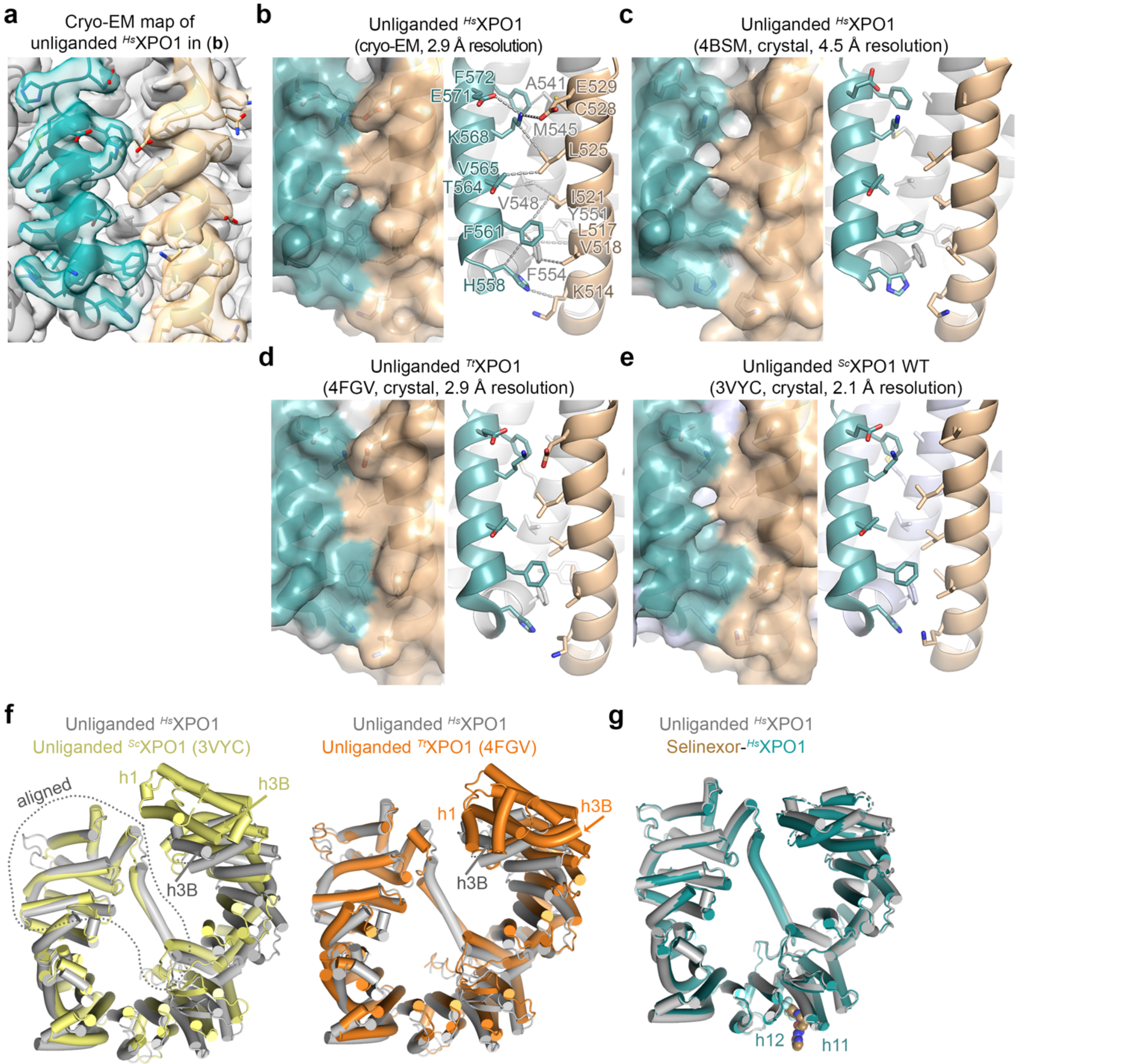


**Extended Data Figure 12. Comparing the closed grooves of unliganded XPO1.** Cartoons with side chain sticks and surface representations, with the four helices of the XPO1 groove in different colors. The h11A and h12A helices (front) in brown and teal, respectively, while the h11B and h12B helices (back) are in gray. The gray back helices should mostly be blocked from view when the groove is closed.

**a-e,** The closed grooves of unliganded XPO1: **a,** Quality of the cryoEM map of the unliganded $^{Hs}$XPO1 groove in this manuscript. **b,** $^{Hs}$XPO1 in (**a**); dashed lines show interactions between groove residues, **c,** crystal structure of $^{Hs}$XPO1, **d,** crystal structure of $^{Ct}$XPO1 and **e,** crystal structure of $^{Sc}$XPO1. The back gray helices in **a-d** are not visible; the grooves are closed.

**f,** Alignment of cryoEM structure of unliganded XPO1 (gray) with: left, the crystal structure of unliganded $^{Sc}$XPO1 (yellow, 3VYC) and right, the crystal structure of unliganded $^{Ct}$XPO1 (orange, 4FGV). The width and pitch of the *Hs* and *Sc* XPO1 rings are similar, while the *Ct* XPO1 ring is larger and slightly more open.

**g,** Alignment of our unliganded XPO1 and selinexor-bound XPO1 cryoEM structures. Both XPO1 rings are similar except for small differences at the N-terminal HEAT repeats and the cNES binding groove (h11/12) is open when bound to selinexor (spheres).

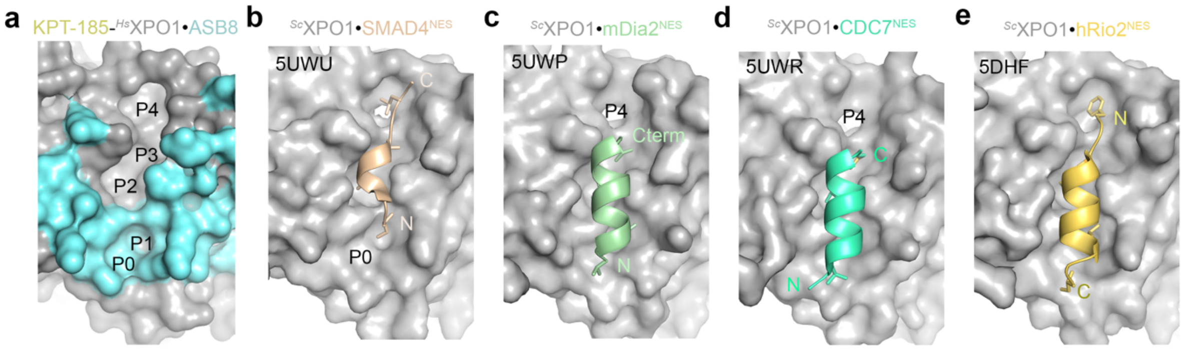


**Extended Data Figure 13. The diverse structures of XPO1-bound NESs.**

**a,** The NES/inhibitor binding groove of XPO1 in the KPT-185-XPO1•ASB8•ELOB/C structure. KPT-185 and ASB8 are not displayed to view the hydrophobic pockets P0-P4 of the XPO1 groove. The XPO1 surface is in gray, with the ASB8 interface colored cyan.

**b-e,** The NES-bound XPO1 grooves from four crystal structures in the PDB: **b,** SMAD4$^{NES}$ (light orange) binds hydrophobic pockets P1-P4, leaving P0 exposed. **c,** mDia2$^{NES}$ (light green) binds P0-P3, leaving P4 exposed, **d,** CDC7$^{NES}$ (aquamarine) also leaves P4 exposed, and **e,** hRio2$^{NES}$ (golden yellow) occupies the entire groove. All XPO1-NES structures in the PDB show that NESs minimally bind P1-P3.

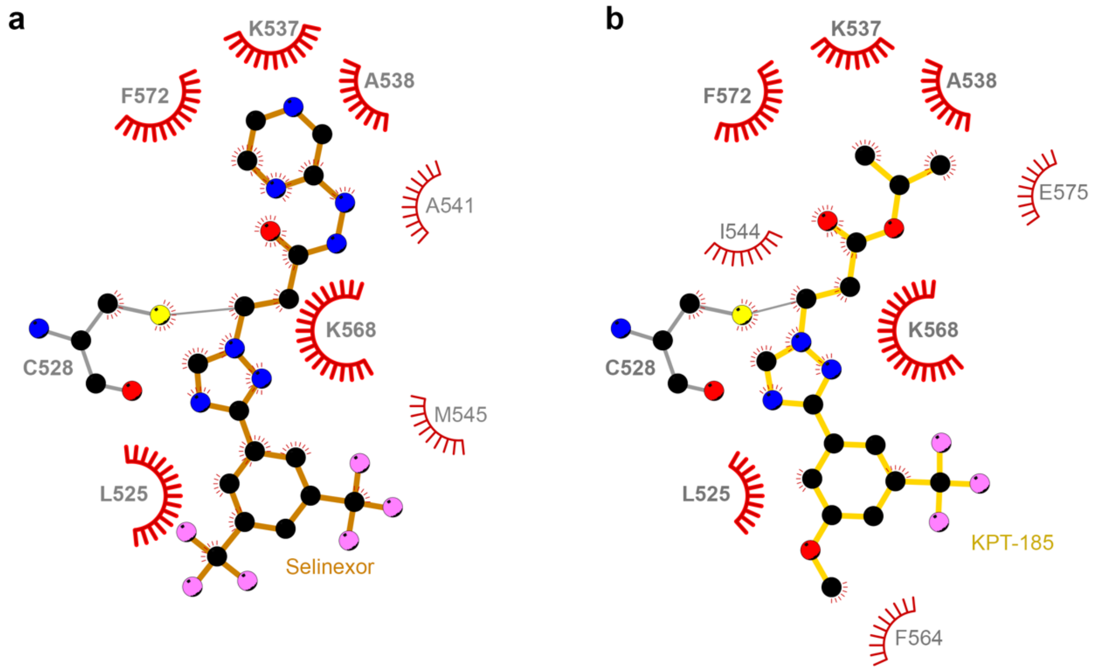


**Extended Data Figure 14. SINE-XPO1 interactions and XPO1-ASB8 interfaces.**

**a-b,** LigPlot+ [1] representation of interactions of selinexor (**a**) or KPT-185 (**b**) in the respective SINE-XPO1•ASB8•ELOB/C cryoEM structures. SINE and the XPO1 C528 side chains are drawn as ball-and-stick (selinexor in orange, KPT-185 in yellow and XPO1 C528 in gray). XPO1 residue labels are in gray. Red arcs with spokes represent nonbonded contacts, and interactions conserved in both the selinexor- and KPT-185-bound structures are labeled in bold. Interactions between SINEs and XPO1 are predominantly hydrophobic. No interactions between the SINEs and ASB8 were detected by LigPlot+.

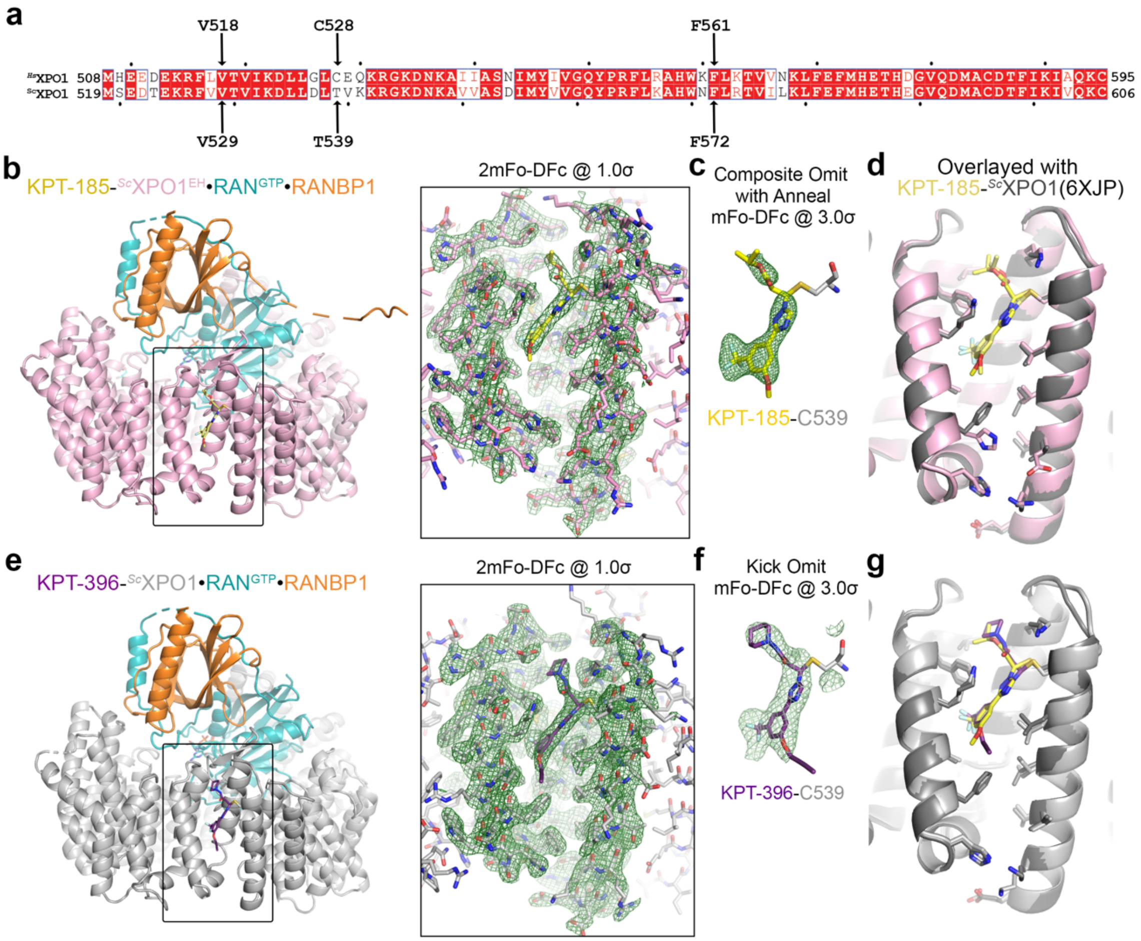


**Extended Data Figure 15. X-ray crystallographic structures of inhibitor-bound $^{Sc}$XPO1*•RAN$^{GTP}$•RANBP1.**

**a,** Sequence alignment of $^{Hs}$XPO1 (top) and $^{Sc}$XPO1 (bottom) at h11/h12. Sites of inhibitor conjugation and of the EH mutation are indicated with arrows.

**b,** Crystal structure of the KPT-185-$^{Sc}$XPO1*$^{EH}$•RAN$^{GTP}$•RANBP1 complex. KPT-185 is shown as yellow sticks. $^{Sc}$XPO1*$^{EH}$ ($^{Sc}$XPO1* is $^{Sc}$XPO1 with humanized groove[23-26]), RAN$^{GTP}$ and RANBP1 are shown as pink, teal and orange cartoons, respectively. 2mFo-DFc map (green mesh) at 1.0σ of the NES groove (highlighted by black box) is shown on the right.

**c,** Composite omit map mFo-DFc (calculated with anneal by omitting the KPT-185 in (**a**)) is shown at 3σ (green mesh).

**d,** KPT-185 bound groove in (**a**), overlayed with the dark gray WT groove of the KPT-185-$^{Sc}$XPO1•RAN$^{GTP}$•RANBP1 crystal structure (PDB ID: 6XJP). The KPT-185-bound grooves of the mutant and WT XPO1 are highly similar except at the sites of the EH mutation.

**e,** Crystal structure of the KPT-396-$^{Sc}$XPO1*•RAN$^{GTP}$•RANBP1 complex. KPT-396 is shown as purple sticks. $^{Sc}$XPO1*, RAN$^{GTP}$ and RANBP1 are shown as light gray, teal and orange cartoons, respectively. 2mFo-DFc map (green mesh) at 1.0σ of the NES groove (highlighted by black box) is shown on the right.

**f,** Kick omit map mFo-DFc (calculated by omitting the KPT-396 in (**e**)) is shown at 3σ (green mesh). Density for the core scaffold of KPT-396 is strong but the density for the propynyl extension is poor. However, the propynyl extension is likely to point out of the groove towards solvent because of the alkyne bond.

**g,** KPT-396 bound groove in (**e**), overlayed with the KPT-185 bound groove (6XJP) as in (**d**).

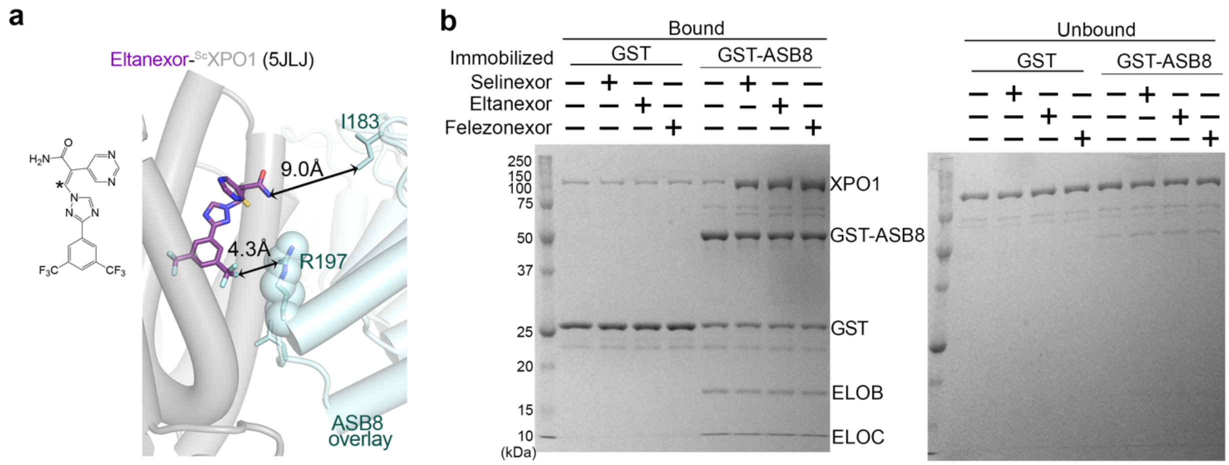


**Extended Data Figure 16. ASB8-XPO1 binding assays with other inhibitors**

**a**, The chemical structure of eltanexor, and the eltanexor-bound XPO1 groove (5JLJ) with ASB8 overlayed.

**b,** Pull-down assay of immobilized GST-ASB8•ELOB/C (negative control is GST) with XPO1 pre-incubated with selinexor, eltanexor or felezonexor. Bound and unbound proteins are visualized by Coomassie SDS-PAGE.

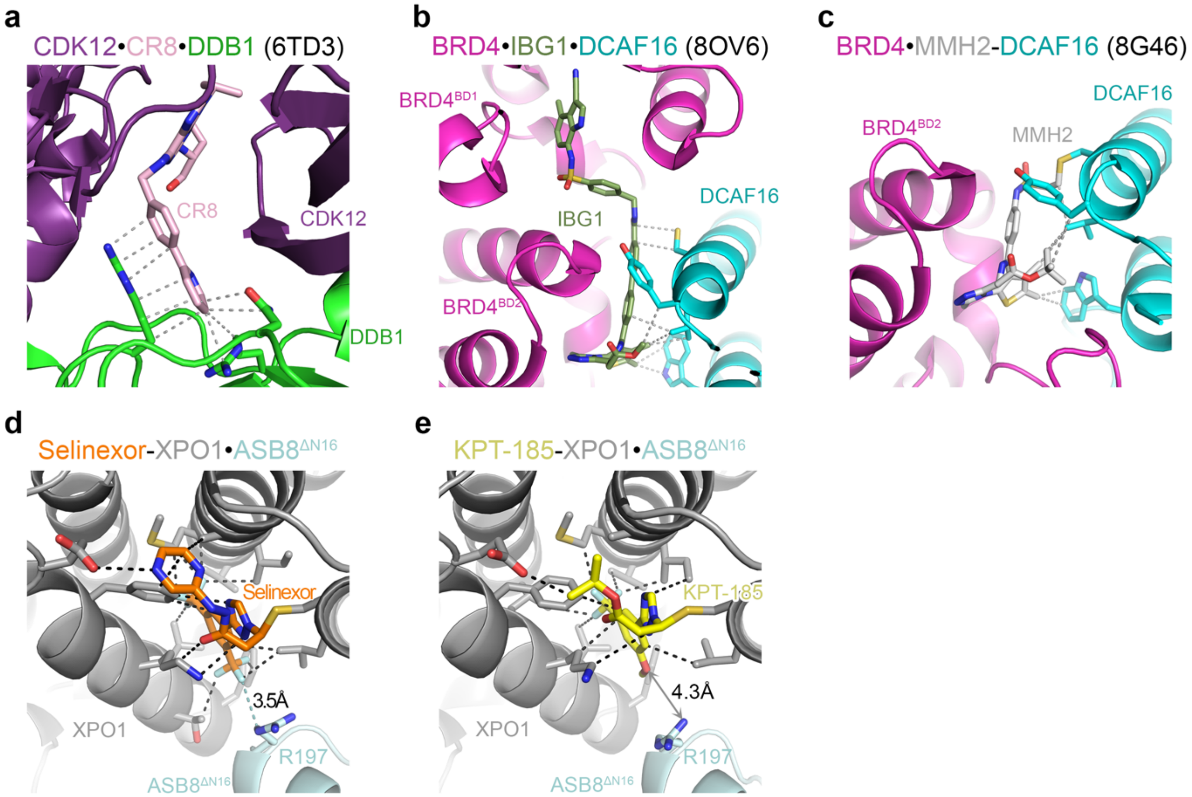


**Extended Data Figure 17. Comparison of substrate-degrader-E3 interactions.**

**a,** The structure of CR8 (light pink) bound CUL4 adaptor protein DDB1 (green) and neosubstrate receptor CDK12 (purple) (6TD3)[2]. Dashed lines indicate contacts within 4 Å. Contacts between CR8 and CDK12 are not displayed, for clarity and focus on CR8-E3 interactions.

**b,** The structure of IBG1 (light green) bound to the CUL4 substrate receptor DCAF16 (cyan) and both bromodomains (BD1 and BD2) of neosubstrate BRD4 (pink) (8OV6)[3]. Contacts between IBG1 and BRD4 are not displayed.

**c,** The structure of MMH2 (light gray) bound to DCAF16 and BD2 of BRD4 (8G46)[4]. Gray dashed lines indicate interactions between MMH2 and DCAF16. Contacts between MMH2 and $BRD4^{BD2}$ are not displayed.

**d-e,** Selinexor (orange, **d**) and KPT-185 (yellow, **e**) bound to XPO1 (dark gray) and $ASB8^{\Delta N16}$ (pale cyan). Black dashed lines indicate SINE-XPO1 contacts, while teal dashed line indicated the single selinexor-ASB8 contact. (**d**). KPT-185 does not contact ASB8 (**e**).

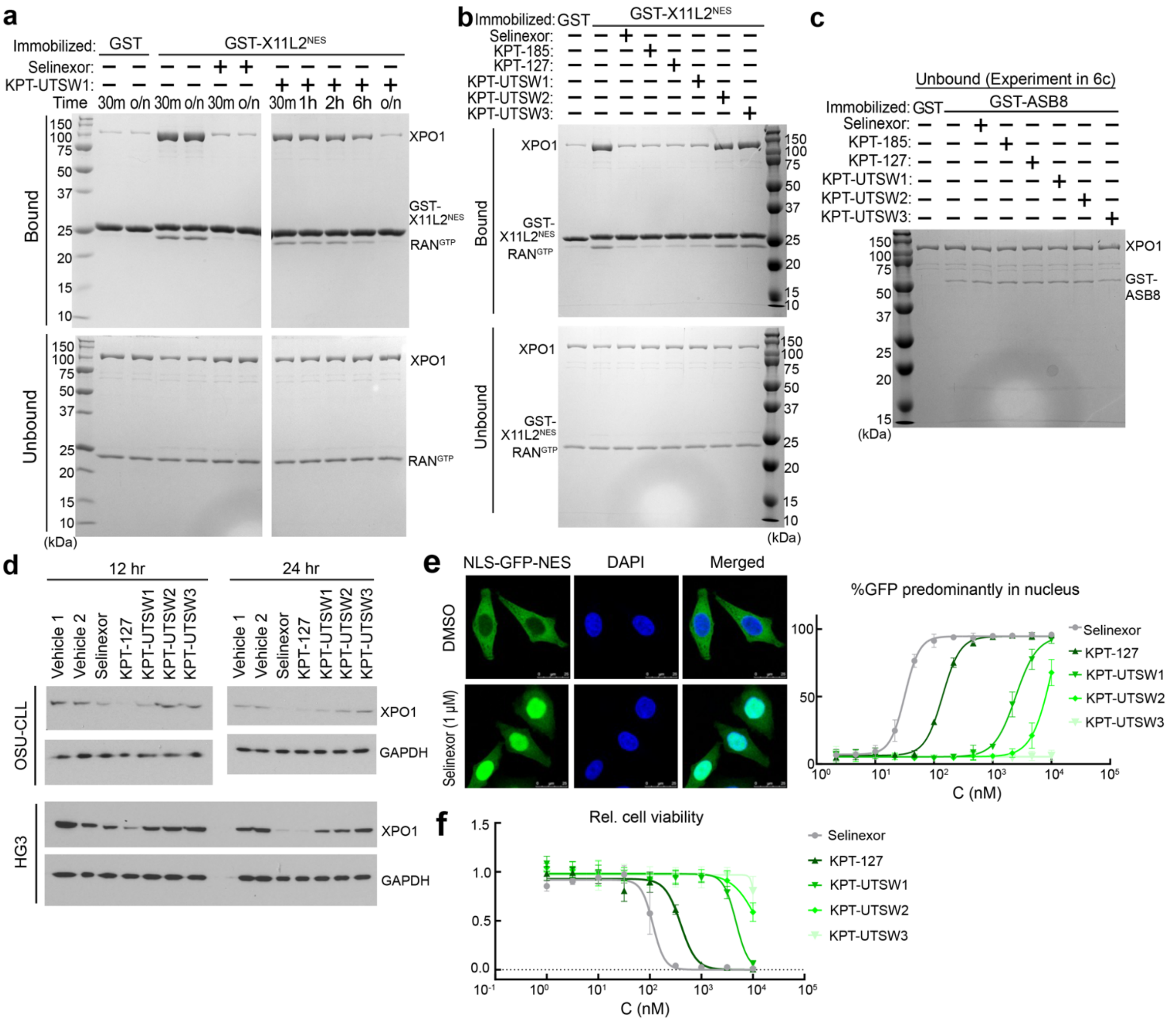


**Extended Data Figure 18. Small SINE analogs: XPO1 conjugation, ASB8 binding and XPO1 degradation.**

**a,** Pull-down assay of immobilized GST-X11L2$^{NES}$ (negative control is GST) with XPO1 pre-incubated with selinexor or KPT-UTSW1 for the indicated time. Bound and unbound proteins are visualized by Coomassie SDS-PAGE. Selinexor effectively inhibits XPO1 (through covalent conjugation) after 30 min of incubation, while KPT-UTSW1 required overnight incubation, likely due to poorer binding to the XPO1 groove, to achieve inhibition of NES binding as effective as selinexor.

**b,** Pull-down assay of immobilized GST-X11L2$^{NES}$ (as in (**a**)) with XPO1 pre-treated with selinexor, KPT-185 or KPT-127 for 30 minutes, or KPT-UTSW1, KPT-UTSW2 or KPT-UTSW3 overnight. KPT-UTSW2 and KPT-UTSW3 do not inhibit NES binding, likely due to decreased efficiency of conjugation to XPO1.

**c,** Unbound proteins from pull-down assay shown in **Figure 6c** to assess different SINE-XPO1 complexes binding to ASB8, visualized by Coomassie staining.

**d,** Degradation of XPO1 by 0.5 μM selinexor or 1 μM of other indicated compounds in CLL cell lines (OSU-CLL and HG3) at 12 h or 24 h. Whole cell lysates were probed by XPO1 or GAPDH (loading control) antibody via WB. Vehicles were negative controls. KPT-127, KPT-UTSW1 and KPT-UTSW2 successfully induced XPO1 degradation in cells.

**e,** Cargo localization assay with an NLS-GFP-NES$^{PKI}$ reporter that is stably expressed in HeLa cells. Representative confocal microscopy images show localization in the cytoplasm under steady state while treatment with selinexor results in nuclear accumulation (left). For quantitative analysis, the percentage of cells in which the fluorescent reporter was primarily located in the nucleus after treatment with SINE analogs for 3 h, was determined (right). The experiment was performed three times. Data are presented as mean ±SD. KPT-127, KPT-UTSW1 and KPT-UTSW2 inhibited nuclear export by XPO1, with a progressive decrease in inhibition (KPT-127 > KPT-UTSW1 > KPT-UTSW2). The progressive decrease correlates with diminished *in vitro* inhibition of XPO1–NES binding shown in (**b**), as well as XPO1 degradation in cells shown in (**c**), which indicates reduced efficiency of covalent conjugation to XPO1 across the series. Decreased conjugation efficiency thus resulted in progressively less XPO1 degradation.

**f,** MTS viability assay in HAP1 cells following 72 h treatment with a dilution series of SINE analogs. Data were normalized to untreated cells and represent mean ±SD of three separate experiments performed in triplicate.

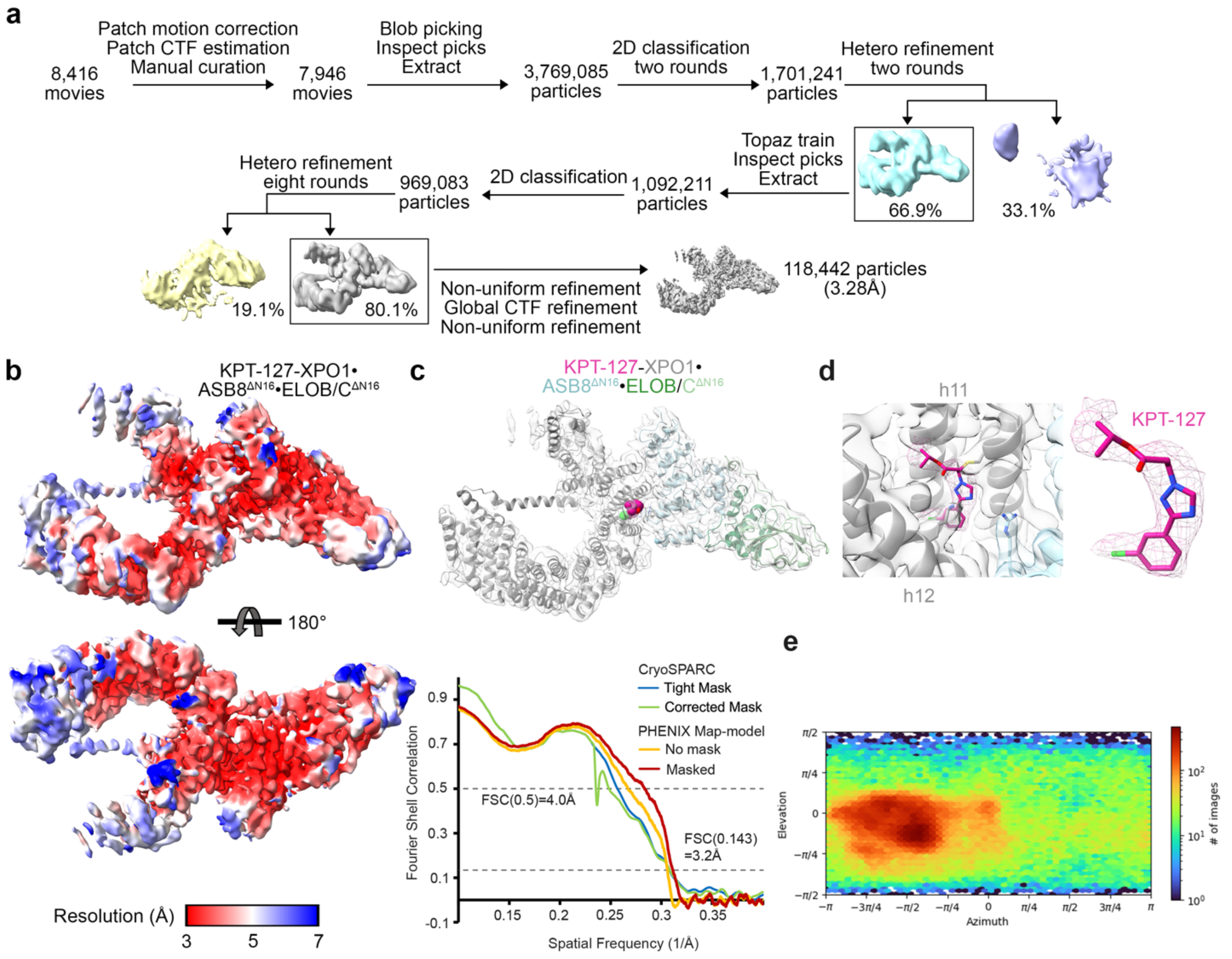


**Extended Data Figure 19. CryoEM maps and statistics for KPT-127-XPO1•ASB8$^{\Delta N16}$• ELOB/C.**

**a,** The flowchart of single particle analysis of the KPT-127-XPO1•ASB8$^{\Delta N16}$•ELOB/C complex.

**b,** The final map of KPT-127-XPO1•ASB8$^{\Delta N16}$•ELOB/C colored by local resolution as indicated by the scale.

**c,** The refined KPT-127-XPO1•ASB8$^{\Delta N16}$•ELOB/C structure overlayed with map in (**b**), accompanied by cryoSPARC and PHENIX map-model FSCs on the bottom.

**d,** A view of the KPT-127(dark pink stick and mesh)-binding groove of XPO1 (gray surface and cartoon). ASB8 cartoon and map in cyan.

**e,** Viewing direction distribution plot for the particles used in reconstruction for map in (**b**).

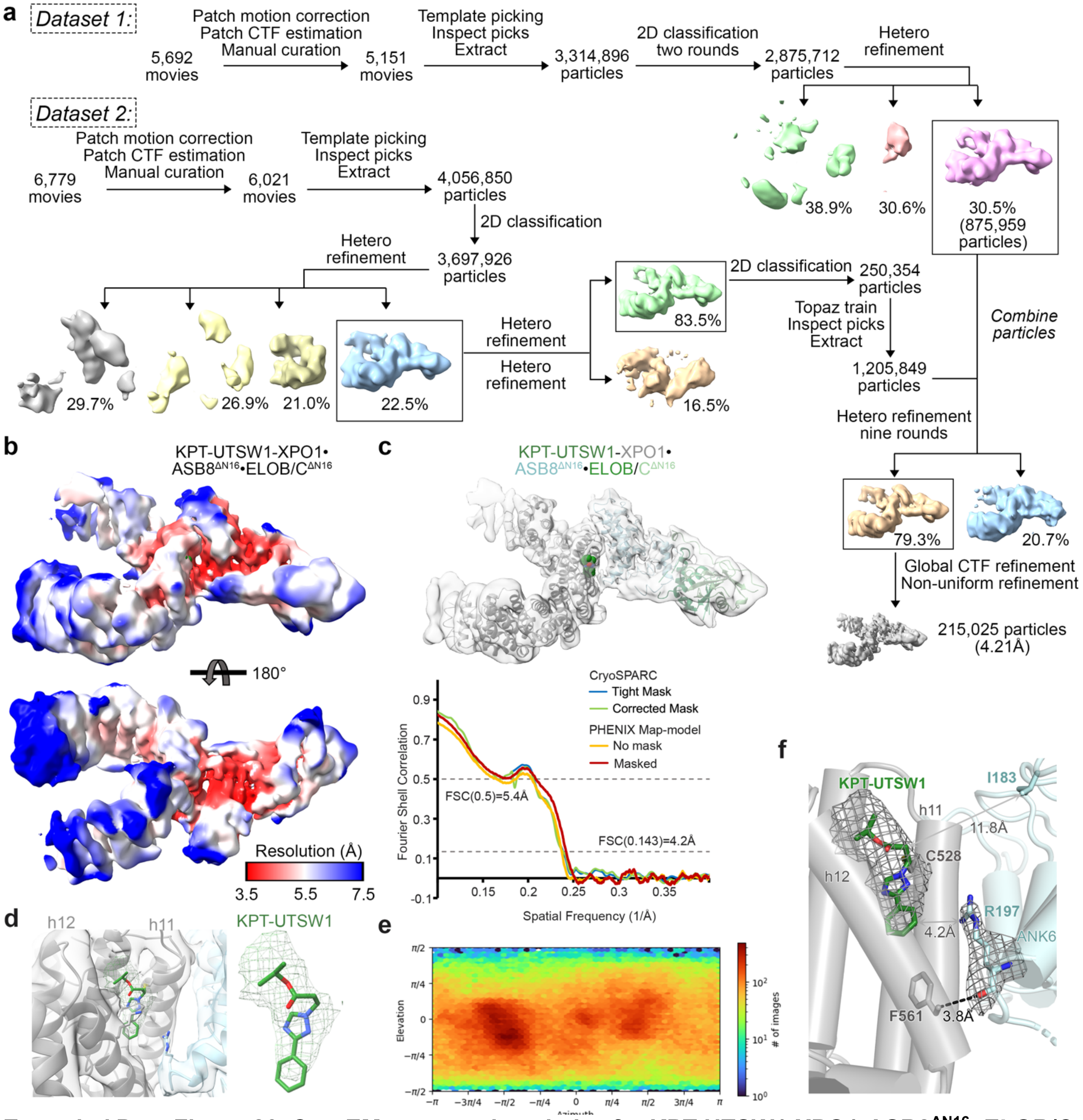


**Extended Data Figure 20. CryoEM maps and statistics for KPT-UTSW1-XPO1•ASB8$^{\Delta N16}$• ELOB/C.**

**a,** The flowchart of single particle analysis of the KPT-UTSW1-XPO1•ASB8$^{\Delta N16}$•ELOB/C complex.

**b,** The final map of KPT-UTSW1-XPO1•ASB8$^{\Delta N16}$•ELOB/C colored by local resolution as indicated by the scale.

**c,** The refined KPT-UTSW1-XPO1•ASB8$^{\Delta N16}$•ELOB/C structure overlayed with map in (**b**), accompanied by cryoSPARC and PHENIX map-model FSCs on the bottom.

**d,** A view of the KPT-UTSW1(green stick and mesh)-binding groove of XPO1 (gray surface and cartoon). ASB8 cartoon and map in cyan.

**e,** Viewing direction distribution plot for the particles used in reconstruction for map in (**b**).

**f,** Side view of the KPT-UTSW1-XPO1•ASB8$^{\Delta N16}$ interface. CryoEM map shown at KPT-UTSW1 and ASB8 R197.

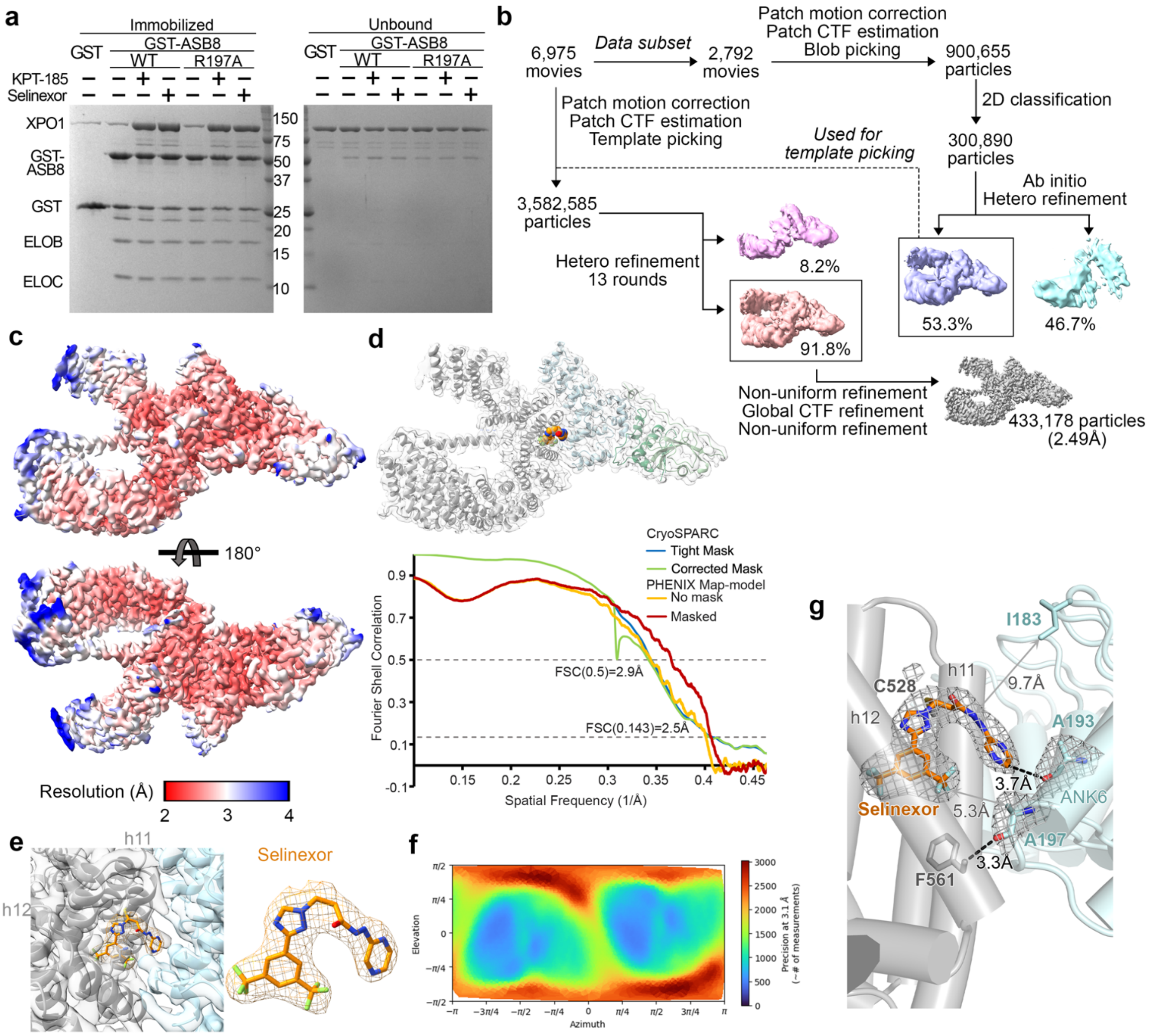


**Extended Data Figure 21. ASB8$^{R197A}$ binds selinexor-XPO1 similarly to ASB8$^{WT}$.**

**a,** A pull-down assay of XPO1 by immobilized GST-ASB8•ELOB/C, in the absence/presence of KPT-185 or selinexor. ASB8$^{R917A}$ binds SINE-XPO1 similarly to ASB8$^{WT}$.

**b,** The flowchart of single particle analysis of the selinexor-XPO1•ASB8$^{\Delta N16/R197A}$•ELOB/C complex.

**c,** The final map of selinexor-XPO1•ASB8$^{\Delta N16/R197A}$•ELOB/C colored by local resolution as indicated by the scale.

**d,** The refined selinexor-XPO1•ASB8$^{\Delta N16/R197A}$•ELOB/C structure overlayed with map in (**c**), accompanied by cryoSPARC and PHENIX map-model FSCs on the bottom.

**e,** A view of the selinexor(orange stick and mesh)-binding groove of XPO1 (gray surface and cartoon). ASB8 cartoon and map in cyan.

**f,** Viewing direction distribution plot for the particles used in reconstruction for map in (**c**).

**g,** Side view of the interface in selinexor-XPO1•ASB8$^{\Delta N16/R197A}$•ELOB/C. Map density of selinexor, ASB8 R197A and A193 is overlayed and shown as gray mesh. The selinexor density shows its pyrazinylpropenehydrazide arm clearly pointed away from the XPO1 groove and towards ASB8 A193 to occupy the space introduced by the ASB8 R197A mutation.

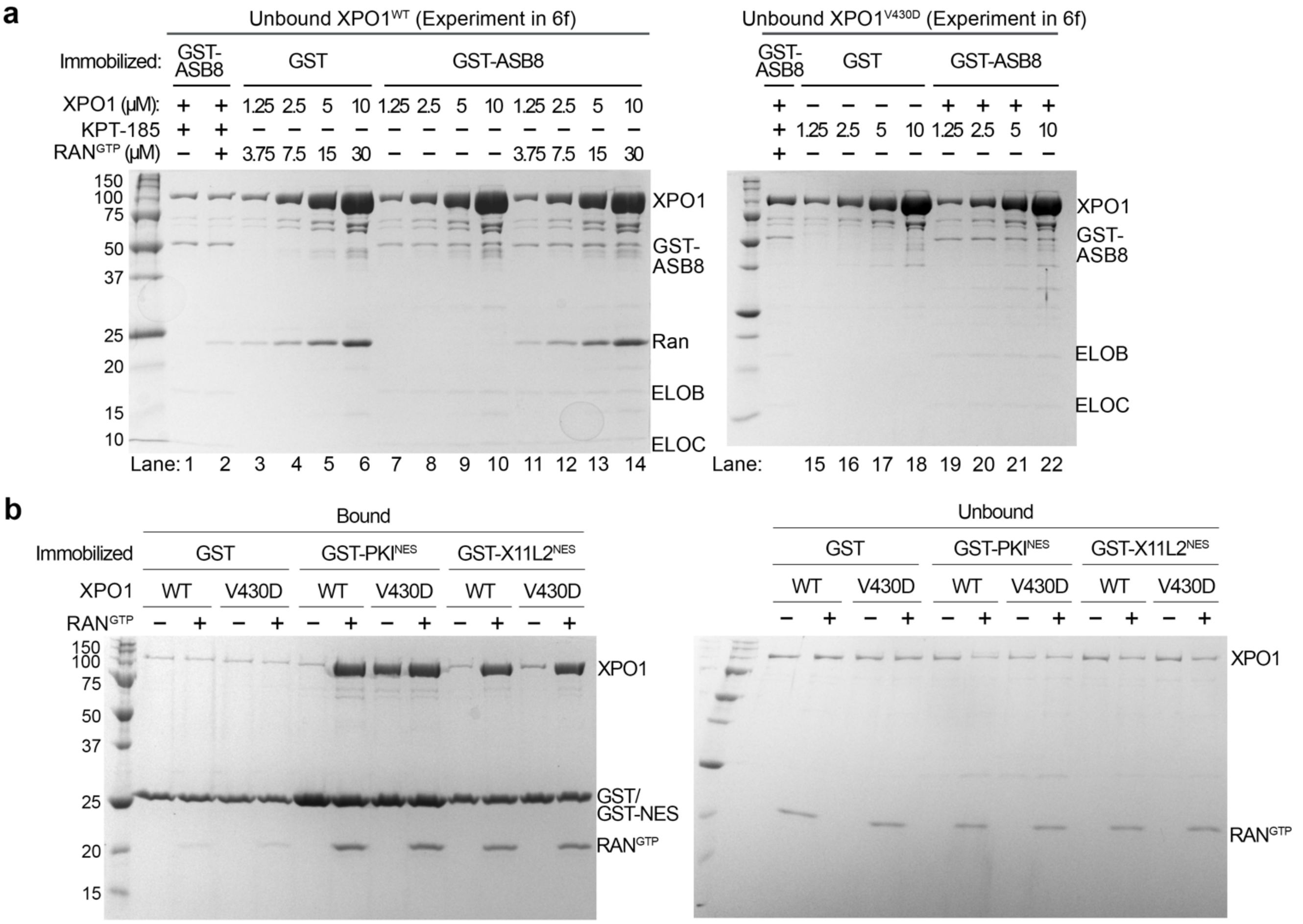


**Extended Data Figure 22. Controls for binding assays of GST-ASB8•ELOB/C with XPO1 in absence of KPT-185.**

**a**, Unbound proteins from the direct pull-down assay shown in **Figure 6g** of XPO1 binding to immobilized GST-ASB8•ELOB/C, in the absence/presence of KPT-185, $RAN^{GTP}$. $XPO1^{WT}$ shown on left, and $XPO1^{V430D}$ shown on right.

**b,** Pull-down assay of immobilized GST, $GST\text{-}PKI^{NES}$ or $GST\text{-}X11L2^{NES}$ with $XPO1^{WT}$ or $XPO1^{V430D}$, with and without $RAN^{GTP}$. $PKI^{NES}$ has a higher affinity than $X11L2^{NES}$ for XPO1 and can bind $XPO1^{V430D}$ without $RAN^{GTP}$, indicating that the V430D mutation shifts XPO1 to the groove open state.

**a**

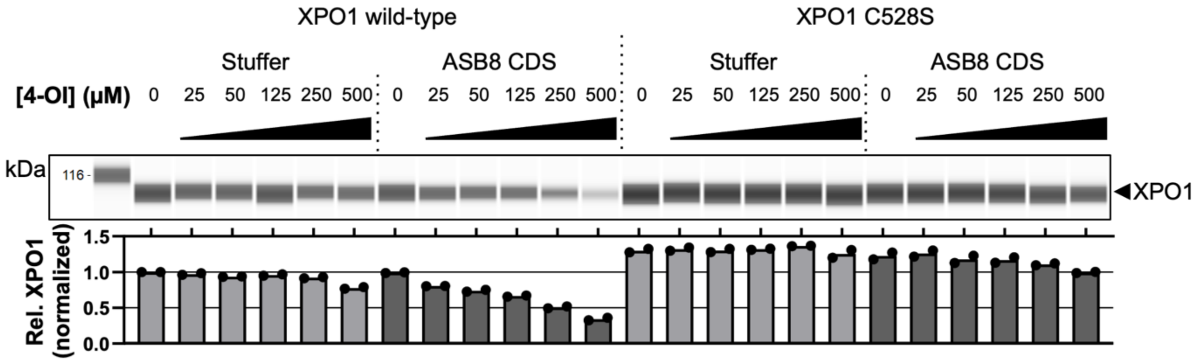


**b**

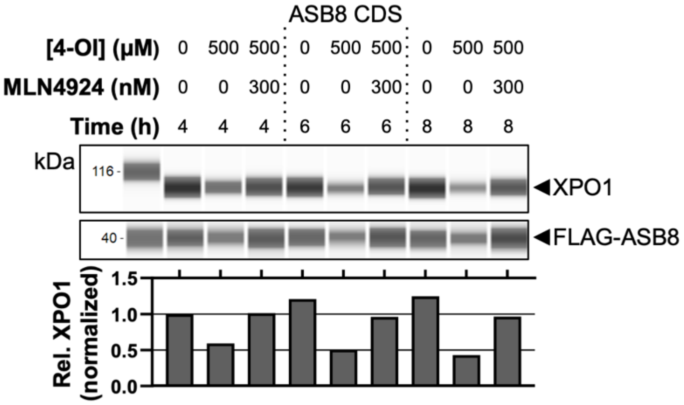


**Extended Data Figure 23. The metabolite 4-octyl itaconate engages XPO1 and promotes ASB8-mediated proteasomal degradation**

**a,** Degradation of XPO1 in HAP1 cells stably transduced with a safe-targeting versus *ASB8* targeting sgRNA or with a non-coding RNA sequence (stuffer) versus ASB8 coding sequence. Cells were treated overnight with increasing concentrations of 4-OI followed by lysis and immunodetection of XPO1. XPO1 peak area was normalized to total protein content per lane and further divided by the untreated control in lane 1. Data collected from two independent experiments.

**b,** Time-of-drug-addition assay in HAP1 cells stably overexpressing ASB8. 4-OI induced XPO1 degradation was observed after 4, 6 and 8 h and could effectively be rescued by addition of MLN4924. XPO1 peak area was normalized to total protein content per lane and further divided by the untreated control in lane 1. Data collected from a single experiment.

**Extended Data Table 1. CryoEM data collection and structure refinement**

| | $RAN^{GTP}$•selinexor-XPO1•$ASB8^{ΔN16}$•ELOB/C (EMD-70460) (PDB 9OGB) | KPT-185•XPO1•$ASB8^{ΔN16}$•ELOB/C (EMD-70461) (PDB 9OGC) | Selinexor•XPO1 (EMD-70459) (PDB 9OGA) | Unliganded XPO1 (EMD-70458) (PDB 9OG9) |
|---|---|---|---|---|
| **Data collection and processing** | | | | |
| Microscope and detector | Titan Krios/ Gatan K3 | Titan Krios/ Gatan K3 | Titan Krios/ Gatan K3 | Titan Krios/ Falcon 4 |
| Magnification | 105,000 | 105,000 | 105,000 | 165,000 |
| Voltage (kV) | 300 | 300 | 300 | 300 |
| # of movies | 4982 | 4746 | 5852 | 8568 |
| Electron exposure ($e^-/Å^2$) | 60 | 50 | 60 | 60 |
| Defocus range (μm) | -0.9 to -2.4 | -1.1 to -2.4 | -0.9 to -2.2 | -0.9 to -2.2 |
| Pixel size (Å) | 0.417 | 0.42 | 0.415 | 0.738 |
| Symmetry imposed | C1 | C1 | C1 | C1 |
| # of particles extracted | 1,821,140 | 1,066,266 | 1,197,263 | 1,245,354 |
| # of final particles | 78,241 | 205,462 | 431,664 | 365,756 |
| Map resolution (Å) | 3.25 | 3.37 | 3.37 | 2.93 |
| FSC threshold | 0.143 | 0.143 | 0.143 | 0.143 |
| $CC_{mask}/CC_{volume}$ | 0.79/0.79 | 0.78/0.78 | 0.80/0.80 | 0.85/0.85 |
| **Refinement and validation** | | | | |
| Initial model used | XPO1 (6TVO), $RAN^{GTP}$ (3M1I), $ASB8^{ΔN16}$•ELOB/C | Selinexor•XPO1, ASB8 (AF-Q9H765-F1 ELOB/C (6V9H) | Unliganded XPO1 | XPO1 (3GB8) |
| Model resolution (Å) | 3.1/3.2/3.8 | 3.2/3.3/3.7 | 3.2/3.3/3.7 | 2.8/2.9/3.1 |
| FSC threshold | 0/0.143/0.5 | 0/0.143/0.5 | 0/0.143/0.5 | 0/0.143/0.5 |
| Model composition | | | | |
| Nonhydrogen atoms | 13122 | 9758 | 7421 | 7321 |
| Protein residues | 1629 | 1222 | 912 | 902 |
| Ligand | 3 (V6A, GTP, MG) | 1 (K85) | 1 (V6A) | 0 |
| R.m.s. deviations | | | | |
| Bond lengths (Å) | 0.005 | 0.003 | 0.003 | 0.002 |
| Bond angles (°) | 0.823 | 0.479 | 0.588 | 0.466 |
| Mean B-factors [ligand] | 173.58 [92.5] | 183.84 [136.08] | 148.94 [130.38] | 124.05 |
| MolProbity score | 1.79 | 1.39 | 1.41 | 1.04 |
| Clashscore | 9.14 | 6.75 | 7.41 | 2.59 |
| Poor rotamers (%) | 0.97 | 0 | 0 | 0 |
| Ramachandran plot | | | | |
| Favored (%) | 95.67 | 97.92 | 98.12 | 98.65 |
| Allowed (%) | 4.02 | 2.08 | 1.77 | 1.35 |
| Outliers (%) | 0.31 | 0 | 0.11 | 0 |

**Extended Data Table 2. X-ray crystallography data collection and refinement statistics.**

| **Data collection** | | |
|---|---|---|
| | **KPT-185-$^{Sc}$XPO1*$^{EH}$• RAN$^{GTP}$•RANBP1** | **KPT-396-$^{Sc}$XPO1*• RAN$^{GTP}$•RANBP1** |
| Space group | $P4_32_12$ | |
| Cell dimensions a=b, c (Å) | 106.08, 303.93 | 105.58,304.56 |
| Resolution range (Å) | 50.00 – 2.37 (2.41 – 2.37) | 50.00 – 2.41 (2.45 – 2.41) |
| Unique reflections | 71232(3485) | 66485 (3182) |
| Multiplicity | 8.7(8.1) | 8.3 (7.9) |
| Data completeness (%) | 99.8 (99.3) | 99.1 (95.9) |
| $R_{merge}$[a] /$R_{pim}$[b] (%) | 8.9 (>100) / 3.4 (46.8) | 9.8 (>100) / 3.4 (34.0) |
| $I/\sigma(I)$ | 19.4 (1.3) | 19.2 (1.6) |
| $CC_{1/2}$ (last resolution shell) | 0.509 | 0.772 |
| **Refinement statistics** | | |
| Resolution range (Å) | 47.0 – 2.37 (2.43 – 2.37) | 49.9 – 2.41 (2.47 – 2.41) |
| No. of reflections $R_{work}$/$R_{free}$ | 66477/2000 (2819/87) | 60841/2000 (2296/78) |
| Data completeness (%) | 93.2 (58.0) | 90.5 (86.0) |
| Atoms (non-H protein/ligand & ions/$H_2O$) | 10791/65/420 | 10869/84/403 |
| $R_{work}$/$R_{free}$ (%) | 19.2/23.7 (28.3/29.9) | 18.6/22.4 (27.8/24.0) |
| R.m.s.d. Bond length (Å)/angle (°) | 0.003/0.483 | 0.002/0.462 |
| Mean B-value (Å$^2$) Protein/ligands & ions/$H_2O$ | 47.4/38.4/37.1 | 46.2/44.4/36.7 |
| Ramachandran plot (%)[b] (favored /disallowed) | 97.5/0.08 | 97.4/0.15 |
| ML coordinate error | 0.31 | 0.28 |
| Missing residues | A: 1-7, 188-193; B: 62, 69-77, 201; C: 970-985, 1054-1058 | A: 1-7, 187-194; B: 62-78, 201; C: 686-689, 1054-1058 |
| PDB code | | |

Data for the outermost shell are given in parentheses.

[a] $R_{merge} = 100\ \Sigma_h\Sigma_i|I_{h,i} - \langle I_h\rangle|/\Sigma_h\Sigma_i\langle I_{h,i}\rangle$, where the outer sum (h) is over the unique reflections and the inner sum (i) is over the set of independent observations of each unique reflection.

[a] $R_{pim} = 100\ \Sigma_h\Sigma_i\,[1/(n_h - 1)]^{1/2}|I_{h,i} - \langle I_h\rangle|/\Sigma_h\Sigma_i\langle I_{h,i}\rangle$, where $n_h$ is the number of observations of reflections **h**.

[b] As defined by the validation suite MolProbity in PHENIX (Chen, V.B., Arendall, W.B.A., Headd, J.J., Keedy, D.A., Immormino, R.M., Kapral, G.J., Murray, L.W., Richardson, J.S., Richardson, D.C. (2010) *MolProbity*: all-atom structure validation for macromolecular crystallography. *Acta Cryst*. **D66**, 12-21.).

**Extended Data Table 3. CryoEM data collection and structure refinement**

| | Selinexor-XPO1•ASB8$^{\Delta N16(R197A)}$•ELOB/C (EMD-70462) (PDB 9OGD) | KPT-127-XPO1•ASB8$^{\Delta N16}$•ELOB/C (EMD-70463) (PDB 9OGE) | KPT-UTSW1-XPO1•ASB8$^{\Delta N16}$•ELOB/C (EMD-70464) (PDB 9OGF) |
|---|---|---|---|
| **Data collection and processing** | | | |
| Microscope and detector | Titan Krios/Falcon 4 | Titan Krios/Falcon 4i | Titan Krios/Gatan K3 |
| Magnification | 165,000 | 165,000 | 105,000 |
| Voltage (kV) | 300 | 300 | 300 |
| # of movies | 6,256 | 7,946 | 11,172 |
| Electron exposure ($e^{-}$/Å$^2$) | 55 | 50 | 60 |
| Defocus range (µm) | -0.9 to -2.4 | -0.9 to -2.4 | -0.9 to -2.2 |
| Pixel size (Å) | 0.738 | 0.7296 | 0.413 |
| Symmetry imposed | C1 | C1 | C1 |
| # of particles extracted | 3,582,585 | 1,092,211 | 2,081,808 |
| # of final particles | 433,178 | 118,442 | 215,025 |
| Map resolution (Å) | 2.49 | 3.28 | 4.21 |
| FSC threshold | 0.143 | 0.143 | 0.143 |
| $CC_{mask}$/$CC_{volume}$ | 0.84/0.85 | 0.80/0.80 | 0.77/0.76 |
| **Refinement and validation** | | | |
| Initial model used | XPO1•ASB8$^{\Delta N16}$•ELOB/C | XPO1•ASB8$^{\Delta N16}$•ELOB/C | XPO1•ASB8$^{\Delta N16}$•ELOB/C |
| Model resolution (Å) | 2.4/2.5/2.7 | 3.1/3.2/3.5 | 4.0/4.1/4.8 |
| FSC threshold | 0/0.143/0.5 | 0/0.143/0.5 | 0/0.143/0.5 |
| Model composition | | | |
| Nonhydrogen atoms | 9,978 | 9,260 | 7,738 |
| Protein residues | 1,244 | 1,154 | 964 |
| Ligand | 1 (V6A) | 1 (KPT-127) | 1 (KPT-UTSW1) |
| R.m.s. deviations | | | |
| Bond lengths (Å) | 0.003 | 0.004 | 0.003 |
| Bond angles (°) | 0.469 | 0.698 | 0.548 |
| Mean B-factors [ligand] | 102.26 [85.15] | 121.89 [92.5] | 271.13 [285.75] |
| MolProbity score | | | |
| Clashscore | 3.05 | 9.60 | 6.24 |
| Poor rotamers (%) | 0 | 0.1 | 0 |
| Ramachandran plot | | | |
| Favored (%) | 97.89 | 96.75 | 96.26 |
| Allowed (%) | 2.11 | 3.07 | 3.74 |
| Outliers (%) | 0 | 0.18 | 0 |

### References


1 Laskowski, R. A. & Swindells, M. B. LigPlot+: multiple ligand-protein interaction diagrams for drug discovery. *J Chem Inf Model* **51**, 2778-2786 (2011). https://doi.org:10.1021/ci200227u

2 Slabicki, M. *et al.* The CDK inhibitor CR8 acts as a molecular glue degrader that depletes cyclin K. *Nature* **585**, 293-297 (2020). https://doi.org:10.1038/s41586-020-2374-x

3 Hsia, O. *et al.* Targeted protein degradation via intramolecular bivalent glues. *Nature* **627**, 204-211 (2024). https://doi.org:10.1038/s41586-024-07089-6

4 Li, Y. D. *et al.* Template-assisted covalent modification underlies activity of covalent molecular glues. *Nat Chem Biol* **20**, 1640-1649 (2024). https://doi.org:10.1038/s41589-024-01668-4
