## Supplemental Methods for "SINE compounds activate exportin-1 degradation via an allosteric mechanism"

**General Information:** All NMR experiments were recorded on Bruker Ascend-400 spectrometer. Data for 1H and 13C NMR spectra are reported as follows: chemical shift (δ, ppm), integration, multiplicity (s = singlet, d = doublet, t = triplet, q = quartet, dd = doublet of doublets, ddd = doublet of doublets of doublets, dt = doublet of triplets and m = multiplet) and coupling constant (Hz). The 7.21 resonance of residual $CHCl_3$ for proton spectra and the 77.04 ppm resonance of $CDCl_3$ for carbon spectra were used as internal references. ISCO flash chromatography was used for purifications. Thin layer chromatography (TLC) was performed on silica gel 60 F254 pre- coated glass plates (0.25 mm) purchased from E. Merck. Visualization of TLC plates was performed using ultraviolet light (254 nm), phosphomolybdic acid (PMA), p-anisaldehyde or KMnO4 stain and heat as developing agents. Mass spectra were acquired on an Agilent technologies 1200 series LC/MS using indicated ionization methods.

**Materials:** Chemicals were purchased from Aldrich or Combi-block and used without purification unless otherwise noted. Deuterated solvents were purchased from Cambridge Isotope Laboratories, Inc. and Sigma Aldrich. All reactions were conducted in oven- or flame-dried glassware under an inert atmosphere of nitrogen.

**Experimental Procedures:**

**I. General Procedure:**

In a flame-dried round bottom flask triazole derivative (1.0 equiv.) was dissolved in anhydrous acetonitrile and cooled at 0 $^{o}$C. Then, $Et_3N$ (1.5 equiv) was added followed by isopropyl propiolate[1] (1.1 equiv) in cooling condition under a nitrogen atmosphere. The reaction mixture was heated at reflux overnight at 90°C, and completion of reaction was monitored on TLC using ethyl acetate: hexane (2:8) as mobile phase and by LCMS analysis. Upon completion, acetonitrile was removed under reduced pressure. The crude reaction mixture was extracted with water (5.0 mL) and dichloromethane (10.0 mL). The organic layer was separated, and the aqueous layer was extracted with dichloromethane (3 x 5.0 mL). The combined organic layers were dried over anhydrous $Na_2SO_4$, filtered, and concentrated in vacuo. The resulting crude mixture was purified by ISCO flash chromatography with EtOAc in *n*-Hexane to afford the corresponding product.

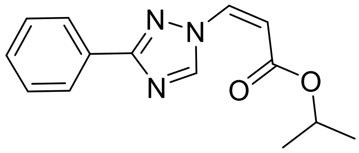

**Isopropyl (*Z*)-3-(3-phenyl-1H-1,2,4-triazol-1-yl)acrylate (KPT-UTSW1):**
Prepared according to the general procedure above with 3-Phenyl-1H-1,2,4-triazole [cas no: 3357-42-4] (80 mg, 0.55 mmol, 1 equiv), $Et_3N$ (110 μL, 0.83 mmol, 1.5 equiv) and isopropyl propiolate (68 μL, 0.61 mmol, 1.1 equiv). The crude product was purified by ISCO flash chromatography (Hexanes: EtOAc, 90:10) to afford the product as white solid (45 mg, 32% yield). **TLC:** $\mathbf{R_f}$ = 0.25 (Hexanes: EtOAc, 90:10); **$^1$H NMR** (400 MHz, $CDCl_3$) δ 9.63 (s, 1H), 8.11 – 8.02 (m, 2H), 7.42 – 7.32 (m, 3H), 7.20 (d, *J* = 10.8 Hz, 1H), 5.58 (d, *J* = 10.9 Hz, 1H), 5.05 (hept, *J* = 6.2 Hz, 1H), 1.24 (d, *J* = 6.3 Hz, 6H); **$^{13}$C NMR** (101 MHz, $CDCl_3$) δ 164.0, 162.4, 147.6, 133.4, 129.9, 129.9, 128.7, 126.8, 107.5, 68.8, 21.8; **ESI MS** for $C_{14}H_{15}N_3O_2$ m/z $[M+H]^+$: calculated: 258.1, found: 258.2.

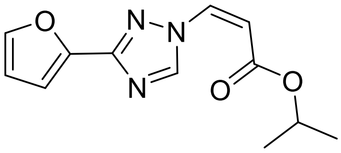

**Isopropyl (*Z*)-3-(3-(furan-2-yl)-1H-1,2,4-triazol-1-yl)acrylate (KPT-UTSW2/KPT-9661):**
Prepared according to the general procedure above with 3-(2-Furyl)-1h-1,2,4-triazole [cas no: 23195-65-5] (50 mg, 0.37 mmol, 1 equiv), $Et_3N$ (77 μL, 0.56 mmol, 1.5 equiv) and isopropyl propiolate (46 μL, 0.41 mmol, 1.1 equiv). The crude product was purified by ISCO flash chromatography (Hexanes: EtOAc, 80:20) to afford the product as white solid (12 mg, 13% yield). **TLC:** $\mathbf{R_f}$ = 0.3 (Hexanes: EtOAc, 80:20); **$^1$H NMR** (400 MHz, $CDCl_3$) δ 9.67 (s, 1H), 7.49 (dd, *J* = 1.8, 0.8 Hz, 1H), 7.20 (d, *J* = 10.9 Hz, 1H), 7.01 (dd, *J* = 3.4, 0.8 Hz, 1H), 6.47 (dd, *J* = 3.4, 1.8 Hz, 1H), 5.60 (d, *J* = 11.0 Hz, 1H), 5.05 (hept, *J* = 6.3 Hz, 1H), 1.24 (d, *J* = 6.3 Hz, 6H); 13C NMR (101 MHz, $CDCl_3$) δ 164.0, 152.7, 147.7, 145.5, 144.0, 133.3, 111.7, 111.2, 107.8, 68.9, 21.8; **ESI MS** for $C_{12}H_{13}N_3O_3$ m/z $[M+H]^+$: calculated: 248.1, found: 248.1.

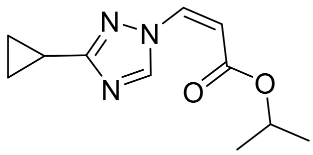


**Isopropyl (*Z*)-3-(3-cyclopropyl-1H-1,2,4-triazol-1-yl)acrylate (KPT-UTSW3):** Prepared according to the modified procedure above with 3-Cyclopropyl-1H-1,2,4-triazole [cas no: 1211390-33-8] (30 mg, 0.27 mmol, 1 equiv), $Et_3N$ (57 μL, 0.41 mmol, 1.5 equiv) and isopropyl propiolate (34 μL, 0.30 mmol, 1.1 equiv). The crude product was purified by ISCO flash chromatography (Hexanes: EtOAc, 85:15) to afford the product as white solid (23 mg, 38% yield). **TLC: R*f*** = 0.4 (Hexanes: EtOAc, 80:20); **$^1$H NMR** (400 MHz, $CDCl_3$) δ 9.46 (s, 1H), 7.05 (d, *J* = 10.9 Hz, 1H), 5.49 (d, *J* = 11.0 Hz, 1H), 5.02 (hept, *J* = 6.3 Hz, 1H), 2.05 – 1.94 (m, 1H), 1.22 (d, *J* = 6.2 Hz, 6H), 0.99 – 0.90 (m, 4H); **$^{13}$C NMR** (101 MHz, $CDCl_3$) δ 166.7, 164.1, 147.0, 133.2, 106.4, 68.6, 21.8, 8.7, 8.0; **ESI MS** for $C_{11}H_{15}N_3O_2$ m/z $[M+H]^+$: calculated: 222.1, found: 222.1.

**Spectral Data:**

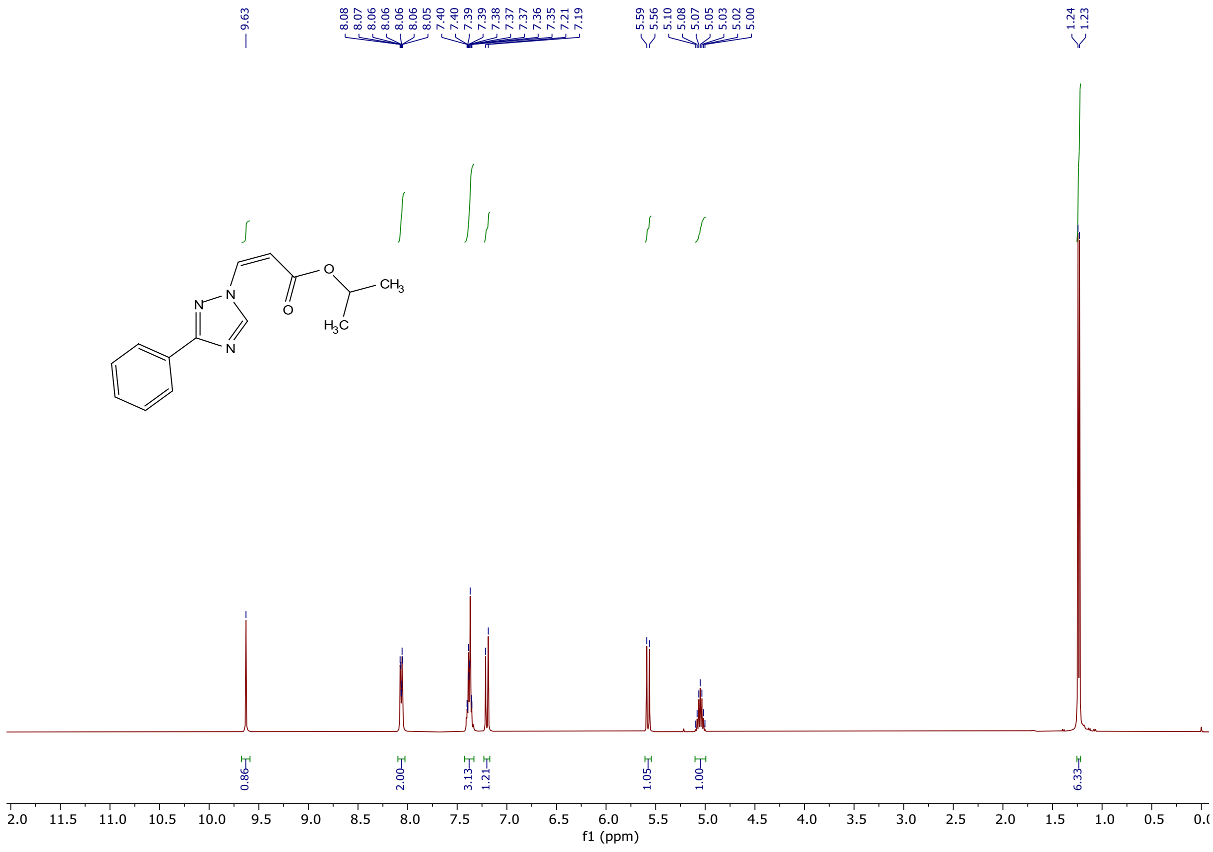

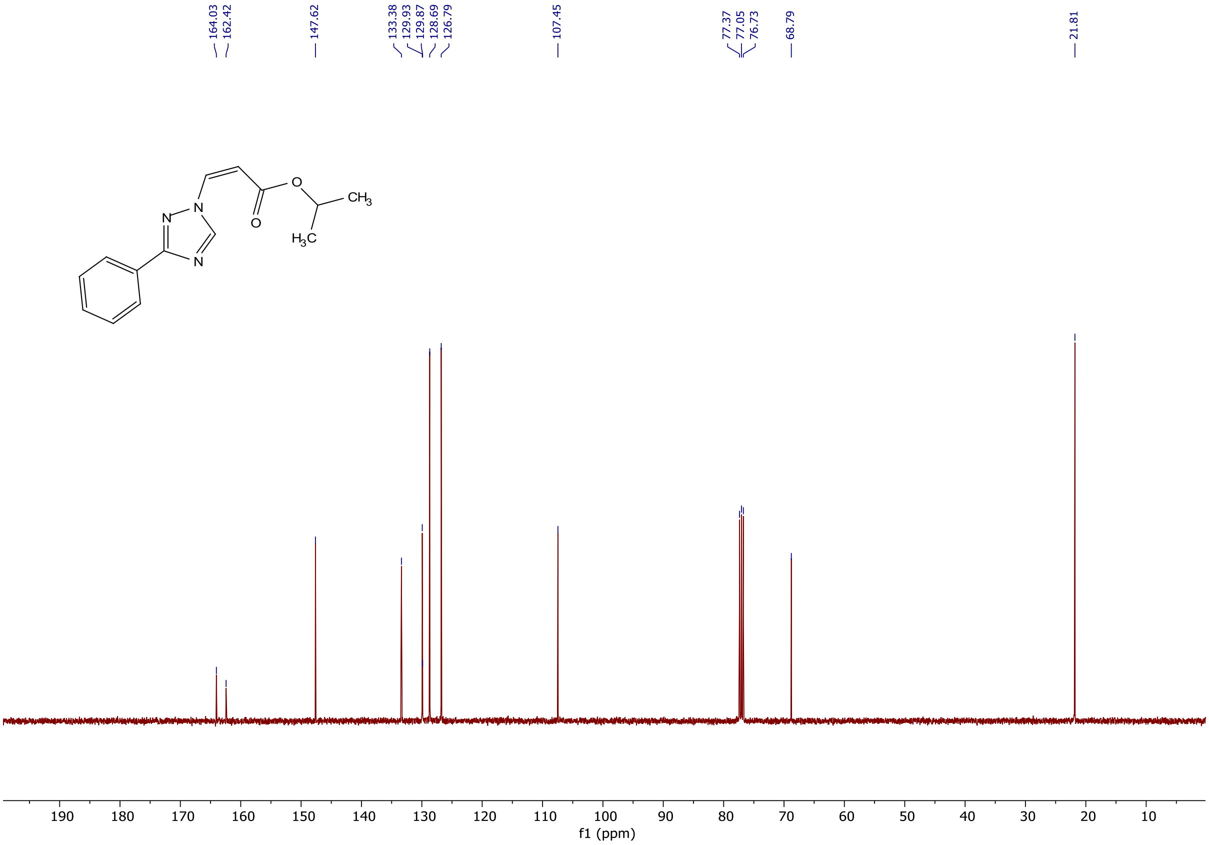
164.03
162.42
147.62
133.38
129.93
129.87
128.69
126.79
107.45
77.37
77.05
76.73
68.79
21.81
N
N
N
O
O
CH3
H3C
190 180 170 160 150 140 130 120 110 100 90 80 70 60 50 40 30 20 10
f1 (ppm)

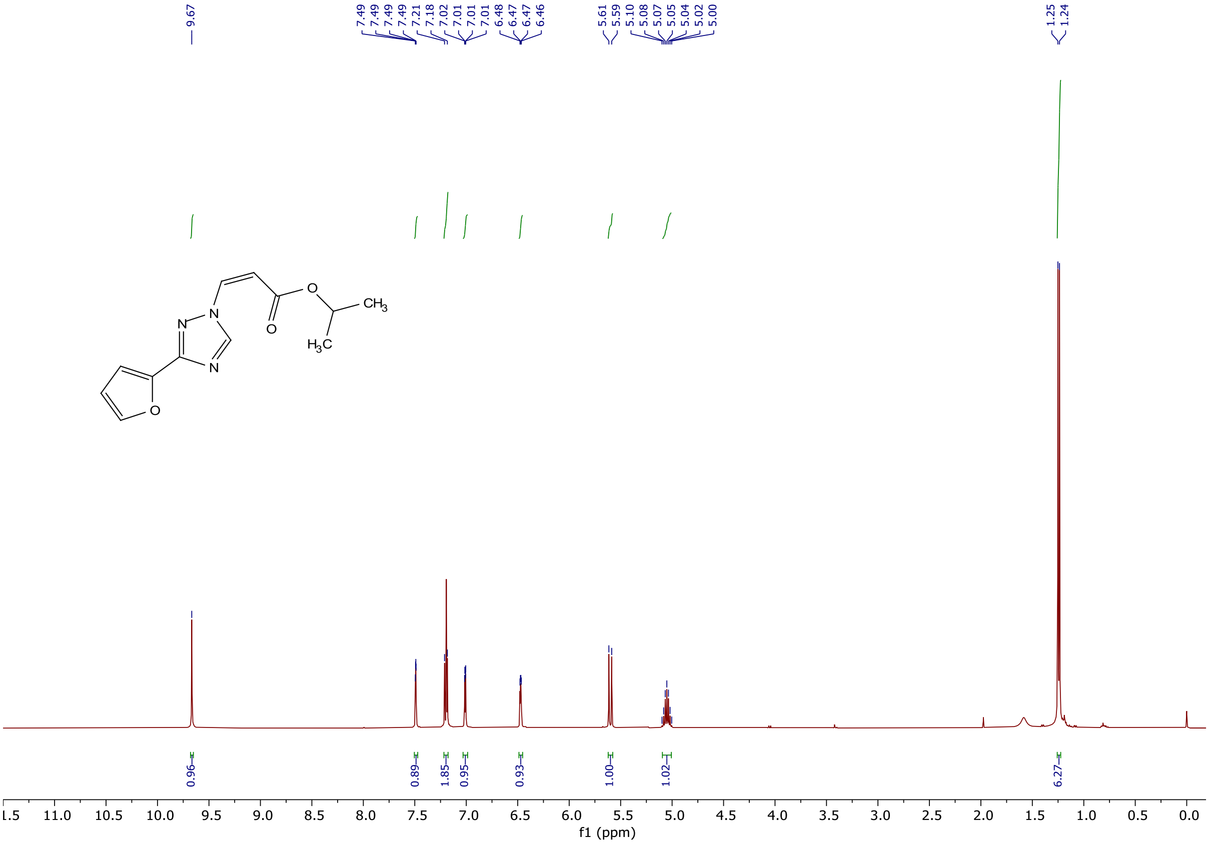
9.67
7.49
7.49
7.49
7.49
7.21
7.18
7.02
7.01
7.01
7.01
6.48
6.47
6.47
6.46
5.61
5.59
5.10
5.08
5.07
5.05
5.04
5.02
5.00
1.25
1.24
N
N
N
O
O
O
CH3
H3C
0.96
0.89
1.85
0.95
0.93
1.00
1.02
6.27
11.0 10.5 10.0 9.5 9.0 8.5 8.0 7.5 7.0 6.5 6.0 5.5 5.0 4.5 4.0 3.5 3.0 2.5 2.0 1.5 1.0 0.5 0.0
f1 (ppm)

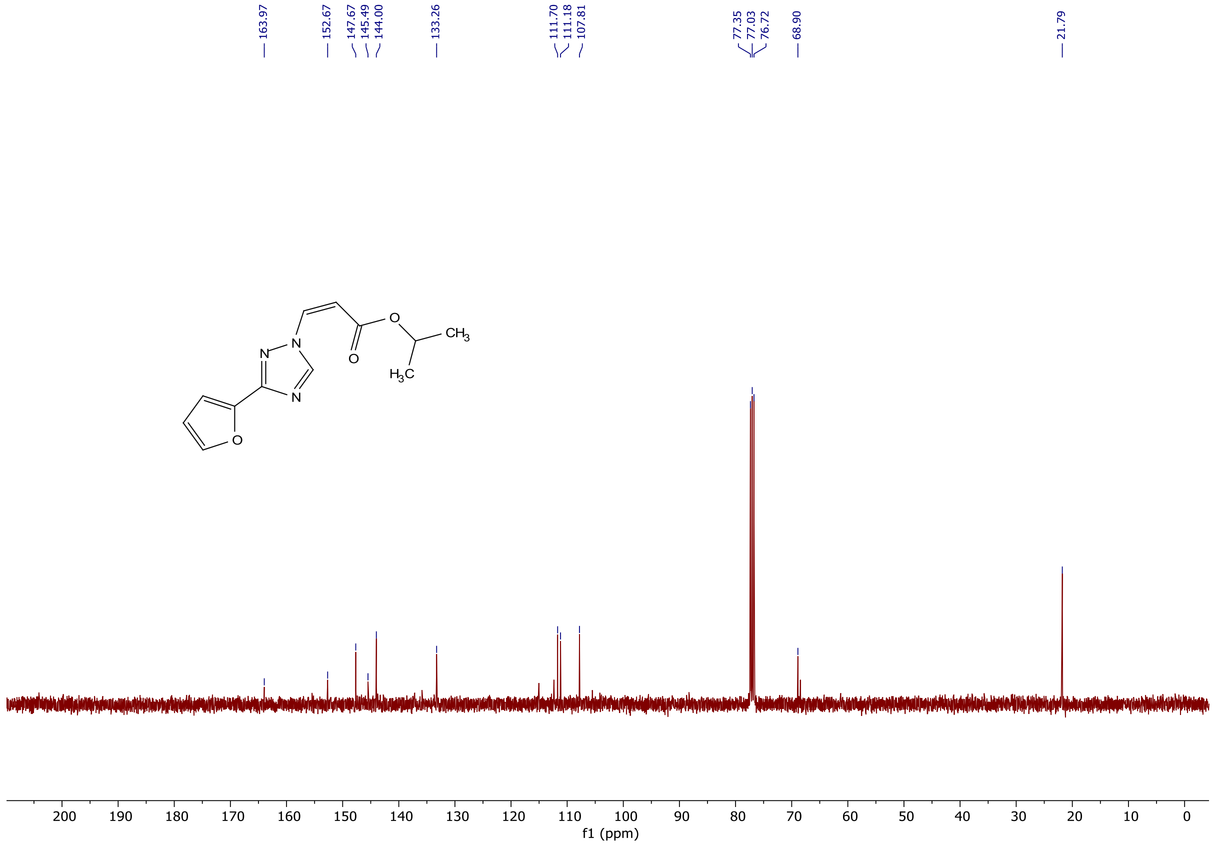
163.97
152.67
147.67
145.49
144.00
133.26
111.70
111.18
107.81
77.35
77.03
76.72
68.90
21.79
N
N
N
O
O
O
$CH_3$
$H_3C$
200 190 180 170 160 150 140 130 120 110 100 90 80 70 60 50 40 30 20 10 0
f1 (ppm)

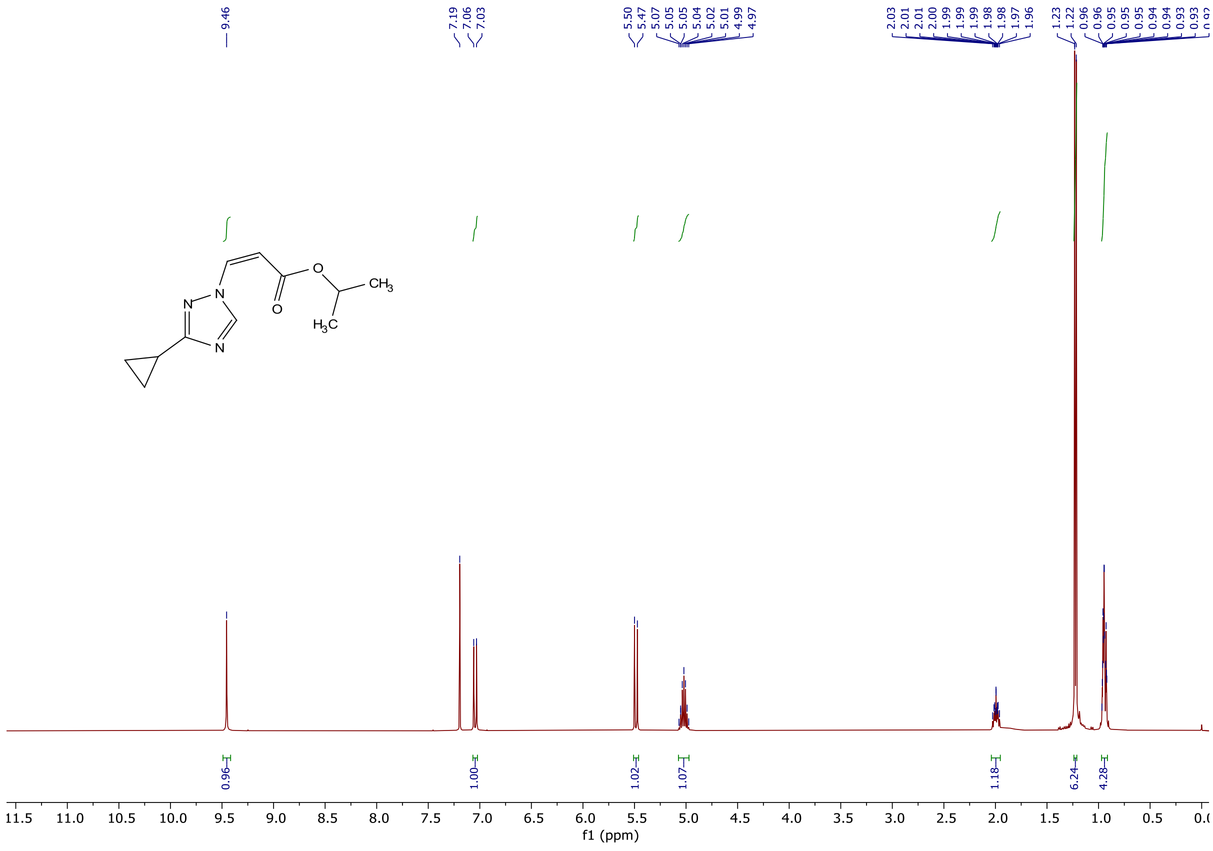
9.46
7.19
7.06
7.03
5.50
5.47
5.07
5.05
5.05
5.04
5.02
5.01
4.99
4.97
2.03
2.01
2.01
2.00
1.99
1.99
1.99
1.98
1.97
1.96
1.23
1.22
0.96
0.96
0.95
0.95
0.95
0.94
0.94
0.93
N
N
N
O
O
$CH_3$
$H_3C$
0.96
1.00
1.02
1.07
1.18
6.24
4.28
11.5 11.0 10.5 10.0 9.5 9.0 8.5 8.0 7.5 7.0 6.5 6.0 5.5 5.0 4.5 4.0 3.5 3.0 2.5 2.0 1.5 1.0 0.5
f1 (ppm)

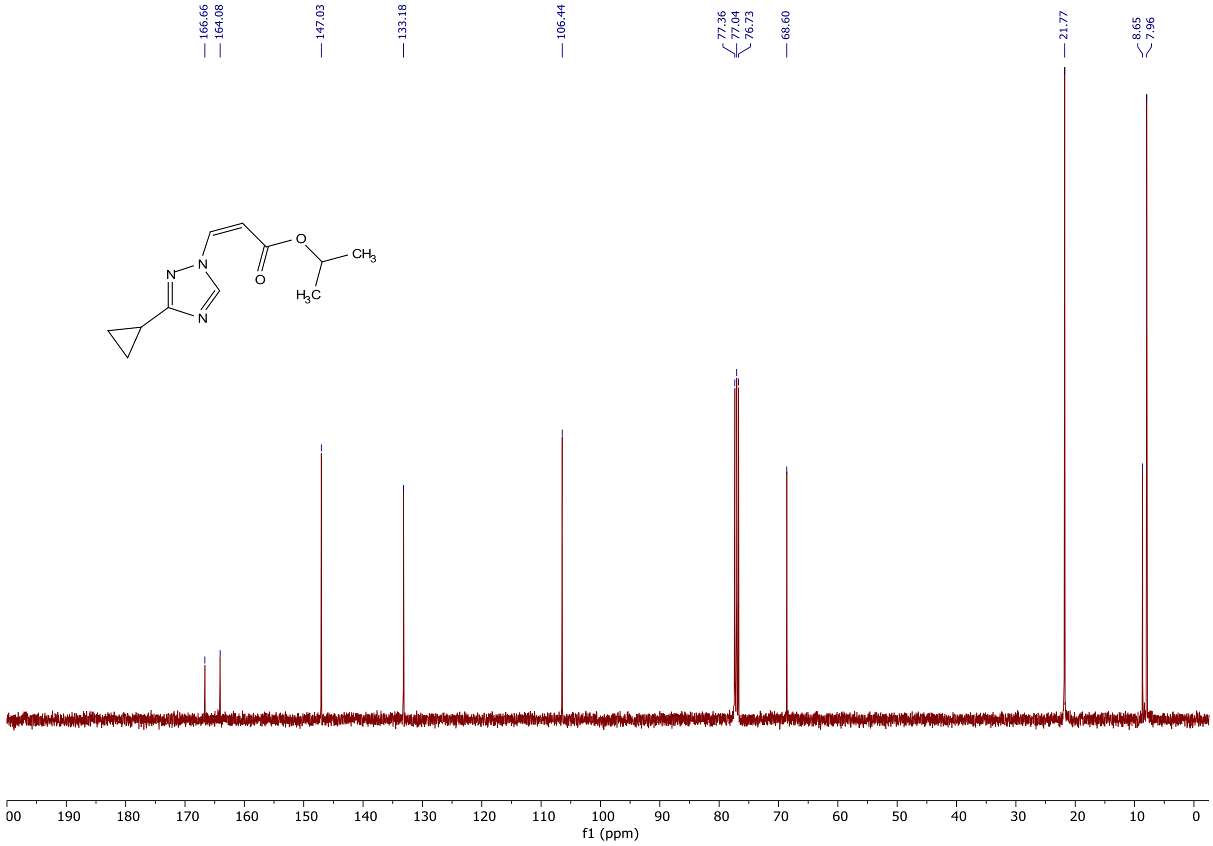


### References:

1. Jung, M. E.; Buszek, K. R., The stereochemistry of addition of trialkylammonium and pyridinium tetrafluoroborate salts to activated acetylenes. Preparation of novel dienophiles for Diels-Alder reactions. *J. Am. Chem. Soc.* **1988***, 110*, 3965–3969.
